# Archaeogenomic Distinctiveness of the Isthmo-Colombian Area

**DOI:** 10.1101/2020.10.30.350678

**Authors:** Marco Rosario Capodiferro, Bethany Aram, Alessandro Raveane, Nicola Rambaldi Migliore, Giulia Colombo, Linda Ongaro, Javier Rivera, Tomás Mendizábal, Iosvany Hernández-Mora, Maribel Tribaldos, Ugo Alessandro Perego, Hongjie Li, Christiana Lyn Scheib, Alessandra Modi, Alberto Gòmez-Carballa, Viola Grugni, Gianluca Lombardo, Garrett Hellenthal, Juan Miguel Pascale, Francesco Bertolini, Gaetano Grieco, Cristina Cereda, Martina Lari, David Caramelli, Luca Pagani, Mait Metspalu, Ronny Friedrich, Corina Knipper, Anna Olivieri, Antonio Salas, Richard Cooke, Francesco Montinaro, Jorge Motta, Antonio Torroni, Juan Guillermo Martín, Ornella Semino, Ripan Singh Malhi, Alessandro Achilli

## Abstract

The recently enriched genomic history of Indigenous groups in the Americas is still meagre concerning continental Central America. Here, we report ten pre-Hispanic (plus two early colonial) genomes and 84 genome-wide profiles from seven groups presently living in Panama. Our analyses reveal that pre-Hispanic demographic changes and isolation events contributed to create the extensive genetic structure currently seen in the area, which is also characterized by a distinctive Isthmo-Colombian Indigenous component. This component drives these populations on a specific variability axis and derives from the local admixture of different ancestries of northern North American origin(s). Two of these ancestries were differentially associated to Pleistocene Indigenous groups that also moved into South America leaving heterogenous footprints. An additional Pleistocene ancestry was brought by UPopI, a still unsampled population that remained restricted to the Isthmian area, expanded locally during the early Holocene, and left genomic traces up to the present.

## INTRODUCTION

Archaeological and genetic evidence suggest that the peopling of sub-arctic America started from Beringia before, during and immediately after late Glacial times (Achilli et al., 2018; Ardelean et al., 2020; Becerra-Valdivia and Higham, 2020; Braje et al., 2017; Skoglund and Reich, 2016; Waters, 2019; Yu et al., 2020). Initial settlement attempts were followed by a more widespread occupation that reached southern South America as early as ∼15 thousand years ago (kya) (Dillehay et al., 2017). Recent studies of ancient and modern genomes describe a complex scenario prior to European contact, with multiple migrations from Beringia, as initially suggested also by mitochondrial DNA (mtDNA) data (Achilli et al., 2013; Brandini et al., 2018; Gómez-Carballa et al., 2018; Llamas et al., 2016; Perego et al., 2009; Perego et al., 2010; Tamm et al., 2007), as well as demographic spreads and admixture events along the two continents (Flegontov et al., 2019; Moreno-Mayar et al., 2018; Posth et al., 2018; Scheib et al., 2018; Schroeder et al., 2018). The great majority of ancestries in early Native Americans (NA, here used to indicate Indigenous groups) derive from an ancestral Beringian population(s) that differentiated sometime between ∼22 and ∼18 kya and likely exhibited genetic sub-structure that may explain the initial late Glacial migration(s) as well as the spread of the so-called UPopA (unknown population in the Americas) whose legacy reappears in Central America ∼8.7 kya, leaving signs in the gene pool of the Mixe (Moreno-Mayar et al., 2018). In unglaciated North America, the first peoples split into two branches called Northern Native American (NNA, or ANC-B) and Southern Native American (SNA, or ANC-A). The most ancient representatives of SNA were individuals who were living on both sides of the Rocky Mountains more than 10 kya: the Clovis-associated Anzick-1 and the Spirit Cave individual associated with Western Stemmed technology. Ancient individuals carrying SNA ancestries crossed the Panama land bridge and entered South America. The fast spread along the southern continent is evidenced by the earliest archaeological human presence in the Southern Cone at 14.6 kya and by ancient human genomes dating more than 9 kya on both sides of the continent: at Cuncaicha (Peru) and Los Rieles (Chile) on the Pacific, Lapa do Santo and Lagoa Santa (Brazil) on the Atlantic. Another unknown population (UPopY) with Australasian ancestry may have contributed to the early peopling of South America as recognized in one sample from the Lagoa Santa site and in some isolated Amazonian groups (e.g. Surui and Karitiana) (Moreno-Mayar et al., 2018; Skoglund et al., 2015).

However, the demographic dynamics underlying many of these events are still uncharacterized, especially at the regional level before and after European contact (Lindo et al., 2017; Nakatsuka et al., 2020; Nägele et al., 2020). The Panamanian land bridge lies between the Atlantic and Pacific Oceans and connects the two American continents. It was the only land bridge during the initial peopling of South America and has remained a crossroads of goods, technologies, ideas and peoples throughout history including more recent colonial times (Cooke, 2005; Cooke et al., 2019; Hernández Mora et al., 2020). In light of Panama’s geographic location, the archaeogenomic study of its past can reveal its demographic history, including movements between North and South America.

### Ethics and Community Engagement

This study involves international collaborative efforts that bring together archaeologists, geneticists, historians, anthropologists, and computer engineers to incorporate existing knowledge with new genomic information about pre-Hispanic, as well as modern Indigenous individuals from the Isthmus of Panama. It was possible with the support of local authorities and Indigenous peoples of Panama and centrally involved local co-authors with years of experience in the Isthmo-Colombian region (JR, TM, MT, JMP, RC, JM and JGM). Samples from the ancient individuals were collected for the ArtEmpire ERC project (CoG 648535) in collaboration with the *Patronato Panamá Viejo* as established by the *Convenio Específico de colaboración entre el Patronato Panamá Viejo de la República de Panamá y la Universidad Pablo de Olavide, de Sevilla, España* signed on January 20^th^, 2016. Excavations were undertaken with the permission of the Republic of Panama’s *Instituto Nacional de Cultura, Dirección Nacional de Patrimonio Histórico, Resolución* DNPH No. 139-16 of November 11^th^, 2016 and *Resolución* DNPH No. 006-18 of January 8^th^, 2018. Selected samples from bone and teeth were exported to Pavia (Italy) and Mannheim (Germany) in accordance with the Permission of the Republic of Panama’s *Instituto Nacional de Cultura, Dirección Nacional de Patrimonio Histórico, Resolución* No. 080-17 DNPH of April 19^th^, 2017 and *Resolución* No. 304-18 DNPH of September 26^th^, 2018. To our knowledge, none of Panama’s present-day Indigenous communities identifies Old Panama’s pre-Hispanic inhabitants as their ancestors.

The collection of the modern Indigenous samples was approved by the *Comité de Bioética de la investigación del Instituto Conmemorativo Gorgas* and undertaken by the *Instituto Conmemorativo Gorgas de Estudios de la Salud* (ICGES, Gorgas Memorial Institute for Health Studies) of Panama. The ICGES explained the project to community leaders in their native languages and sent biological samples to the Department of Biology and Biotechnology of the University of Pavia for DNA extraction and analysis in agreement with the memorandum of understanding (written in English and Spanish) signed on August 9^th^, 2016. All experimental procedures and individual written informed consent forms were also reviewed and approved by the Ethics Committee for Clinical Experimentation of the University of Pavia, Board minutes of April 11^th^, 2013 and October 5^th^, 2010.

The project has been designed to maximize opportunities for public engagement, as testified by ArtEmpire’s first public meeting*, La Primera Globalización en un Arteria del Imperio, 1513-1671,* held in Panama City on April 20-22, 2017, when local researchers emphasized the importance of further attention to the pre-Hispanic population from a genetic perspective. Specific meetings with local interest groups are being organized to discuss the results of this research, as previously done by Prof. A. Achilli, Dr. U.A. Perego and Dr. R. Cooke through the open conference *Descifrando el genoma panameño* at the BIOMUSEO of Panama City (October 16^th^, 2012) to present the results of their previous analyses of uniparental markers (Perego et al., 2012). In order to increase its social impact, the archaeological data on the ancient samples are publicly available in ArtEmpire’s database, translated also in Spanish to increase accessibility for non-English speakers (https://artempire.cica.es/) (Aram et al., 2020), and a documentary has been also released (https://www.youtube.com/watch?v=5BmxppS4oks). The Uninorte hosted an open discussion in Spanish, with the participation of BA, JM and TM, before the launching of the documentary on May 9^th^ 2020.

We are particularly grateful to and acknowledge the people who have shared the ancient and present-day DNA analyzed.

### Panama: archaeology and history

Paleoecological and archaeological data point to a continuous human inhabitation of the Isthmo-Colombian area from approximately 16 kya (Cooke et al., 2013; Ranere and Cooke, 2020). Clear evidence for the cultivation of domesticated plants, including maize (*Zea mays*), manioc (*Manihot esculenta*) and squash (*Cucurbita moschata*), dates back to more than 8-4.5 kya (Linares, 1977a; Linares, 1977b; Linares and Ranere, 1980; Linares et al., 1975; Piperno, 2011; Ranere and Cooke, 2020), while Panama’s first pottery (*Monagrillo Ware*) appeared about 4.5 kya (Martín et al., 2016; Martín et al., 2015). By 3 kya, the western region possesses all the characteristics of coherent historical unit (Greater Chiriquí, which extended into present-day Costa Rica), while this consensus is not available for the central and eastern regions, often termed Greater Coclé and Greater Darién (see specific section in Supplementary Information for further details).

From approximately 500 BCE through 1500 CE relations among neighbors oscillated between cooperation driven by exchange and trade, and conflict over land and resources. Although polities often called chiefdoms expanded and contracted, there is no evidence for aggressive empire-building as in Mexico or the Andes (Helms, 2014). On the eve of the Spanish invasion, historians and most archaeologists agree that much of the central and eastern isthmus was inhabited by Indigenous polities that spoke languages in the Nuclear Chibchan family (with variants of languages in the Chocoan family probably also spoken on the Pacific side) and used the “language of Cueva” as a *lingua franca* (either a trade language or a group of vernaculars) in a linguistically complex region, much as the Huëtar did in Costa Rica (Cooke 2016; Costenla, 2012; Romoli, 1987). Based on partial and fragmentary data, historians have ventured estimates of the pre-Hispanic population of those areas where the language of Cueva was spoken at European contact from 130 to 240 thousand people and archaeologists have identified specific villages capable of sustaining up to 2,400 inhabitants (Cooke, 2005; Cooke et al., 2019; Romoli, 1987). After nearly one millennium of less destructive war and trade among neighbors, European incursions provoked a rapid decline in the region’s Indigenous populations. However, not all of the Indigenous groups experienced simultaneous demographic decline. Historical records suggest that an expansion among the Guna followed the reduction of other Indigenous groups (Castillero Calvo, 2017).

The uneven impact of European colonization and the upheaval it induced, in addition to subsequent migrations, caution against the automatic identification of the territories occupied by specific Indigenous groups before 1500 CE with those they inhabit today (Torres De Arauz, 1999). Although goods also crossed the isthmus in pre-colonial times, the process intensified after the early sixteenth century, when the Spanish established settlements on both sides of the Isthmus to forge a highway for the global transit (and often forced mobility) of persons and goods between the Atlantic and Pacific Oceans.

In order to assess change and continuity in the most radically transformed area of the Isthmus, the under-studied eastern region, most of the ancient remains analyzed in this paper were obtained from the archaeological site of *Panamá Viejo* (Panama City), an area of pre-Hispanic inhabitation and the site of the colonial city from 1519 through 1671 CE (Hernández Mora et al., 2020). Modern sampling, on the other hand, took place in Panama City as well as in the provinces and Indigenous territories. In 2010, Indigenous polities encompassed about 12% of the 3.4 million inhabitants of Panama (*Census 2010*). The most numerous are individuals who identify as the Ngäbe (62.3%), Guna (19.3%) and Emberá (7.5%), followed by smaller polities such as the Chocoan-speaking Wounaan (the Noanomá of Spanish chronicles), and the Buglé, Bribri, and Naso Djërdi (formerly known as Teribe). The contemporary population also includes important numbers of self-identified Afro-Panamanians (“Morenos”) and individuals of mixed Hispano-Indigenous (“Mestizo”) ancestry.

### Panama: genetics

To date uniparental systems have been examined to assess the genetic history of Panama: mtDNA data identified specific lineages predating the Clovis technological horizon (13.2 kya), while the comparison with Y-chromosome data revealed a sex bias in post-colonial times consistent with “more native men perishing or being deprived of reproductive rights than women” (Grugni et al., 2015; Perego et al., 2012). Similar to mtDNA data, patterns of regional genetic continuity in some Indigenous American (IA) communities have been inferred from the analysis of nuclear genomes from continental Central America (Reich et al., 2012), but without ancient DNA (aDNA) data from the lower Isthmian land bridge.

To refine the human genetic history of the Isthmus, we have directly tested and analyzed for the first time autosomal markers of both pre-Hispanic human remains and contemporary Indigenous groups from Panama. Twenty ancient individuals (from 13 pre-Hispanic and seven colonial individuals) were sampled from seven different archaeological excavations along the Pacific coast of Panama City (Figure 1A and Table S1), located within a 2 km radius and extending from the residential area of *Coco del Mar* to the remnants of Old Panama’s Cathedral in *Panamá Viejo*. To examine the demographic history and genetic relationships among the Indigenous populations of Panama, the data from these ancient individuals were compared to newly generated genome-wide data obtained from 84 modern individuals (76 self-identified as associated with five different Indigenous groups plus four self-designated “Moreno” and four self-identified “Mestizo” individuals) sampled in different regions of the Panamanian Isthmus (Figure 1A, Table S2).

**Figure 1.**
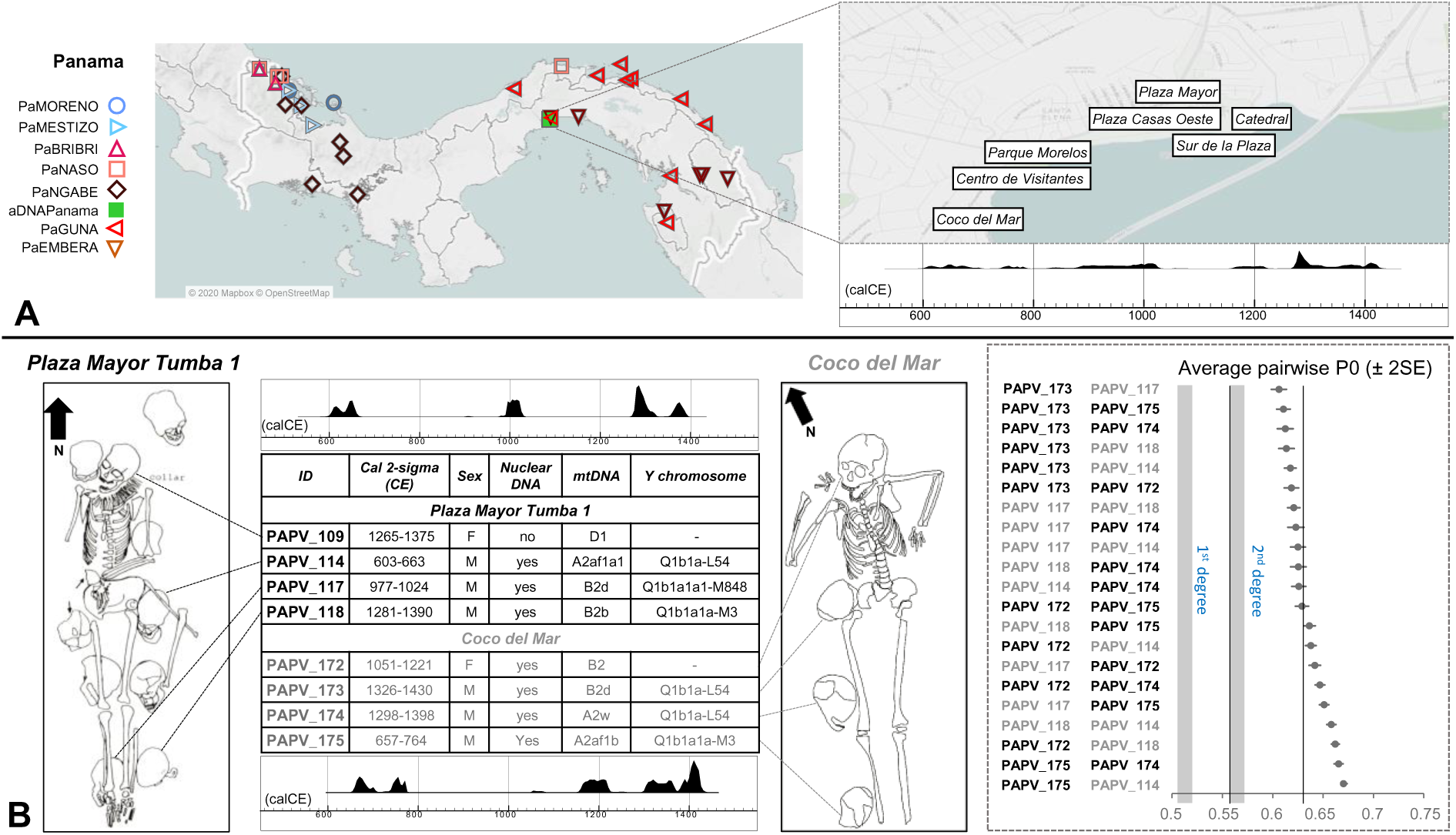
Geographic Locations and Time Ranges of Modern and Ancient Panamanians. (**A**) Map showing the geographic origin of our Panamanian individuals; the inset represents the locations of the archaeological excavations. (**B**) Schematic drawings of *Tumba 1* in the Plaza Mayor site of *Panamá Viejo* and the burial at the *Coco del Mar* site. The table reports mtDNA and Y-chromosome haplogroup affiliations, molecular sex determination and 14C calibrated dates (CE). The sum distributions of all ages combined are shown, separately for the two sites, above (*Tumba 1*) and below (*Coco del Mar*) the table. Calibration dataset: IntCal20. Calibration software: OxCal 4.4.2. The inset on the right shows kinship relationships (extracted from Figure S4) among individuals buried together. *Plaza Mayor Tumba 1* and *Coco del Mar.* IDs (and additional information) are indicated in black and grey, respectively.

## RESULTS AND DISCUSSION

Although the tropical environment and the proximity of the excavation sites to the ocean, with recurrent flooding, challenge the possibility of DNA preservation, we were able to obtain some of the first reliable ancient DNA data from the Isthmus. We assembled low-coverage (≥0.01X) ancient genomes from 12 unrelated individuals (one female and eleven males), including ten from pre-Hispanic times (radiocarbon dated from 603 to 1430 CE). Molecular decay analyses demonstrated the poor preservation of endogenous DNA, but error rate and validation tests confirmed the reliability of the retrieved genomic data, which are comparable with those from other ancient individuals from the Americas (Table S1; Figures S1-S4). In order to characterize the genetics of Panamanian individuals with the greatest possible spatial range and temporal depth, the 12 ancient genomes were compared to genome-wide data from 74 unrelated modern Panamanians (Table S2) and to available modern and ancient data by assembling different datasets (Supplementary Information; Tables S3-S5; Figures S5, S6): 1) a worldwide dataset of modern samples (rWD1560); 2) two datasets of (nearly) “unadmixed” Indigenous Americans (uIA217 and uIA89) obtained by removing individuals with signatures of non-Indigenous genetic contributions using different stringent criteria; and 3) a larger dataset of “admixed” individuals where only genetic fragments inherited from Indigenous individuals were retained, masking (removing) variants not belonging to genetic haplotypes inherited by Indigenous groups (mIA417), to focus on pre-Hispanic interactions. The individuals used for the latter dataset were considered pseudo-haploid in allele frequency analyses, while in haplotype-based methods the individual chromosome pairs were jointly analyzed. Moreover, some of the samples with less than 50% Indigenous ancestry were removed (rmIA311).

### Uniparental lineages of pre-Hispanic Panamanians

The evaluation of uniparental markers revealed the presence of the “pan-American” mtDNA haplogroups A2 and B2 in the pre-Hispanic samples (Figure S7), while two haplogroups, H1j1a and L2a1c2a, typical of Europeans and sub-Saharan Africans, respectively, were identified in the samples taken from colonial ancient individuals (Table S1).

The most represented mtDNA haplogroup of pre-Hispanic Panamanians, A2af1, was previously identified (as A2af) at high frequencies among present-day Panamanians, mainly in the Comarca of Kuna Yala (Perego et al., 2012). It is characterized by the so-called ‘‘Huëtar deletion’’, a peculiar 6-bp control-region deletion initially detected in the Chibchan-speaking *Hu*ë*tar* from Costa Rica (Santos et al., 1994).

The eight pre-contact Y chromosomes are positive for the L54 marker, which characterizes all the Native American branches of haplogroup Q (Figure S8). Two subjects (PAPV118, PAPV175) were further sub-classified as Q1b1a1a-M3 and one (PAPV117) as Q1b1a1a1-M848, the most frequent haplogroups among Indigenous peoples of the Americas (Grugni et al., 2019; Pinotti et al., 2019).

### Archaeological and anthropological significance of two burials in *Panamá Viejo* and *Coco del Mar*

An initial evaluation of the ancient low-coverage genomes made it possible to address long-standing anthropological and archaeological questions regarding the possible genetic relationships among individuals buried together. These cases included the ten human remains (one adult, nearly complete, female skeleton with nine adult male skulls beneath and around her) recovered from a pre-Hispanic burial denominated *Tumba 1* underneath the Plaza Mayor of *Panamá Viejo*, and a similar burial at *Coco del Mar* (approximately 1 km to the west of *Panamá Viejo*), where a female skeleton was found accompanied by three male crania (Figure 1B). Crania interred with prestigious individuals have been interpreted as evidence either of ancestor veneration or of human sacrifice with the ostentation of trophy heads (Mendizabal, 2004; Smith-Guzmán and Cooke, 2018). Arguments in either case draw on presumed (recent or ancestral) tribal and biological relationships. We can now exclude any genetic relatedness among the individuals using genome-wide data. Moreover, the two females exhibit different mtDNA haplogroups (D1 and B2, respectively) with respect to the surrounding male crania (A2af1a1, B2b and B2d in *Tumba 1*; A2af1b, A2w and B2d in *Coco del Mar*) (Figure 1B).

In combination with these genetic results, radiocarbon dates obtained for the pre-Hispanic individuals sampled from *Panamá Viejo* and *Coco del Mar* point toward a more complex and nuanced interpretation. The two female figures, PAPV109 (1265-1375 CE) and PAPV172 (1051-1221 CE), were interred with crania dated from 603 through 1390 CE (2 sigma) in the first case and from 657 to 1430 CE (2 sigma) in the second (Figure 1B). Hence skulls spanning over 700 years, including the area’s earliest and latest pre-Hispanic remains recovered to date, accompanied each of the main individuals. Seven of the skulls that accompanied PAPV109 belonged to individuals who pre-dated her by hundreds of years, and the other two were roughly contemporary. One of the crania buried with PAPV172 belonged to a male individual who lived roughly 500 years before her, and the other two to male individuals deceased and buried over the subsequent 300 years. Drawing upon on Cueva as well as Guna ethnography (Castillero Calvo, 2017: pp. 26, 87, 281-2, 476-8; Fernández de Oviedo, 1853; Fortis, 2013), crania kept for hundreds of years or even deposited after the main interment probably pertained to enemy chiefs whose death in battle guaranteed their spirits’ eternal repose. Their skulls may have provided sorcerers and healers, in this case female seers or *tequina*, a gateway to knowledge about enemies as well as the afterlife. These women entered the next world with the tools of their trade (the skulls), like other individuals interred in *Panamá Viejo* (a “musician” poised as if playing her instrument, and a seated adolescent with flint blades and stingray tail barbs) or mentioned in Spanish chronicles (farmers buried with corn) (Fernández de Oviedo, 1853: Vol. 2, pp. 125-154). The heads of prestigious enemies obtained in warfare would have facilitated the seers’ access to their strength and knowledge, as well as their ability to communicate with other worlds. The different mtDNA lineages of these individuals might also support their origins from various pre-Hispanic groups, since we have found significant differences in the haplogroup distribution among the modern Indigenous populations analyzed here (*p-value* <0.0001). Although the literature contains reference to wives, slaves and loyal servants sacrificed with their chiefs (Fernández de Oviedo, 1853; Romoli, 1987), these burials illustrate a different practice. Their arrangement stands out among a great diversity of pre-Hispanic interments over a wide zone of the Panamanian Pacific, some of these including offerings in local pottery, metal, lapidary, shell and bone-work, and many burial modes, including bones deposited in urns or bundles (Martín, 2002a, b). PAPV172’s head was found beneath a ceramic offering, while five ceramic pots with offerings accompanied PAPV109, who also wore necklace fashioned from thorny oyster shells (*Spondylus* spp.). Within the variety of pre-Hispanic burial patterns observed to date, individuals PAPV109 and 172 appear unique on a local as well as a regional level.

### Deciphering genomic variation in the Isthmus

The analyses conducted in this study facilitate a microgeographic and diachronic assessment of indigenous autosomal variation in this strategic region. The characterization of the pre-colonial genetic histories is clouded by the impacts of colonization. This is evident in the worldwide Principal Component Analyses (PCA) (Figure S9) for some modern populations (e.g. the Colombian “CLM” and the Peruvian “PEL” groups), as well as for some Indigenous Panamanians (i.e. three Bribri individuals), scattered between Indigenous and non-Indigenous populations. However, the colonial genetic impact is even more evident in the distinctive genetic profiles that differentiate the current gene pool of all the Indigenous Panamanian groups as obtained by the ADMIXTURE analyses (Figure 2A).

**Figure 2.**
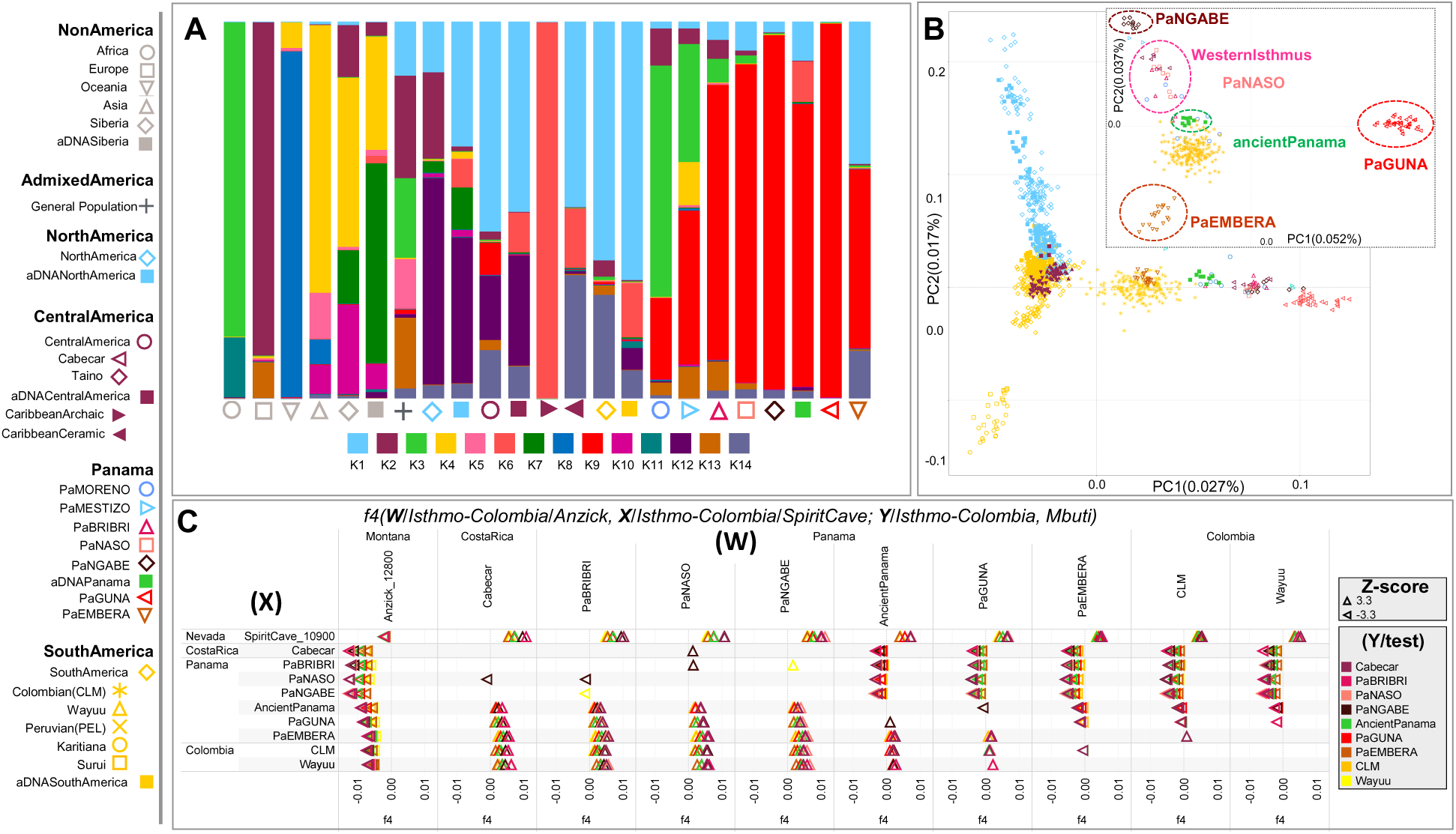
Overview of the Genetic Structure of Ancient and Modern Panamanians. (**A**) ADMIXTURE plot for K = 14, each bar shows the average ancestry proportion of individuals within the same group considering the rWD1,560 dataset plus the American and Siberian ancient individuals. (**B**) Indigenous American PCA analysis including the mIA417 dataset and ancient genomes projected onto uIA217 variability. The inset shows a specific Isthmo-Colombian PCA. (**C**) *f4-statistics* in the form of *(**W**/Isthmo-Colombia/Anzick, **X**/Isthmo-Colombia/SpiritCave; **Y**/Isthmo-Colombia, Mbuti)* considering the uIA89 and mIA417 datasets plus ancient Panamanians (all SNPs). The *f4* values are reported in abscissa. Each tested population (Y) is shown (with triangles pointing to X or W population) only when the initial conformation of the tree is rejected (*p-value∼0.001*, depending on the Z-score), thus visually pointing to the closer population (X or W) in each comparison.

The two groups that experienced a history of admixture, the self-identified “Moreno” and “Mestizo”, reveal large proportions of their genomes not derived from Indigenous peoples of the Americas. Both show a comparable proportion of ancestry predominant in Europeans (K2) and the component common to Africans (K3) is more prevalent in the “Moreno” group; the “Mestizo” are characterized by a component also identified in Asians (K4). The coexistence of different continental genomic ancestries is common in the Americas, due to complex admixture that started during the colonial period (Ongaro et al., 2019). In Panama, where European colonization began in 1502 CE, this is particularly evident in the “Mestizo”, but it is also revealed by individuals, who self-identified as Indigenous and genealogically unadmixed, showing variable amounts of African and European ancestries in their genomes, with the lowest average values in the Guna, followed by the Ngäbe.

The modern and ancient Panamanians are also characterized by a specific Indigenous component, which has been identified considering only modern individuals (K6, Figure S10) as well as with the addition of ancient individuals (K9, Figure 2A). As for the latter analyses, the ancient genomes were projected on the modern variability (Figure S11), as well as explored without projection (Figures 2A, S12). This ancestry drives the Isthmo-Colombian axis, depicted by the first (main) component in the PCA of Indigenous groups (Figure 2B), which includes ancient and modern Panamanians together with the Cabécar from southern Costa Rica and two populations from northern Colombia (the Wayuu and the admixed CLM). The Indigenous groups from the pre-colonial Greater Chiriquí cultural area form two closely related western clusters (one with the Ngäbe and the other including Bribri, Naso and Cabécar). The pre-Hispanic individuals group together in the middle of the Isthmo-Colombian genetic landscape and create a distinct branch in the outgroup *f3-statistics* hierarchical tree, together with a few self-identified “Moreno” (Figure S13), suggesting the integration of pre-Hispanic individuals into newly forming multi-cultural colonial groups. The genetic closeness of the pre-Hispanic individuals is possibly expected when considering the geographic proximity of the archaeological excavations but less expected when taking into account the radiocarbon dates, from 603 to 1430 CE, thus revealing a genetic continuity for almost one thousand years (Table S1). The modern Indigenous populations from the putative Greater Darién unit to the east of the region (Guna, Emberá and northern Colombians) create distinct groups in the PCA plot, well separated from the Greater Chiriquí populations in the west. The details of this genetic sub-structure in the Isthmus became apparent by analyzing the nearly unadmixed Indigenous haplotypes (uIA217 dataset) with fineSTRUCTURE (Figures 3A, S14). Among the five genetic clusters, four are specific of Indigenous Panamanian groups (PaNASO, PaNGABE, PaEMBERA, PaGUNA), while the Bribri individuals form a separate cluster (here called Western Isthmus) with the Cabécar from Costa Rica. The latter branch, together with Naso and Ngäbe, forms a macro-group that might be associated with the geographic region of the pre-colonial Greater Chiriquí cultural area. Genetically distinct are the Emberá and the Guna, suggesting a wider genetic variation in the putative Greater Darién cultural region. The Guna also show the highest level of similarity in both intra- and inter-cluster comparisons (Figure S15), analogous only to two very isolated Amazonian tribes, the Surui and Karitiana (Figure 3B), which preserved an ancient Australasian-related ancestry of the so-called population Y (UPopY) (Moreno-Mayar et al., 2018; Skoglund et al., 2015). We formally looked for UPopY variants in the Isthmus with *f4-statistics* in the form *f4*(Panama, Mixe; Australasia, Mbuti) without finding any significant sign of admixture or gene flow (Figure S16A). The same statistics were also used to formally test the average correlation in allele frequency differences (mixture of ancestries) within the Isthmo-Colombian area (Figure 2C). This analysis provides statistical support to the genetic interactions in the western-isthmian area, eventually extended to Cabécar, Naso and Ngäbe. On the other hand, it reveals close relationships among Emberá and northern Colombians (CLM and Wayuu). Finally, the pre-Hispanic communities inhabiting the Pacific coast in the area of Panama City left significant genetic traces in the Guna when compared with other Isthmo-Colombian populations, except for the Ngäbe.

**Figure 3.**
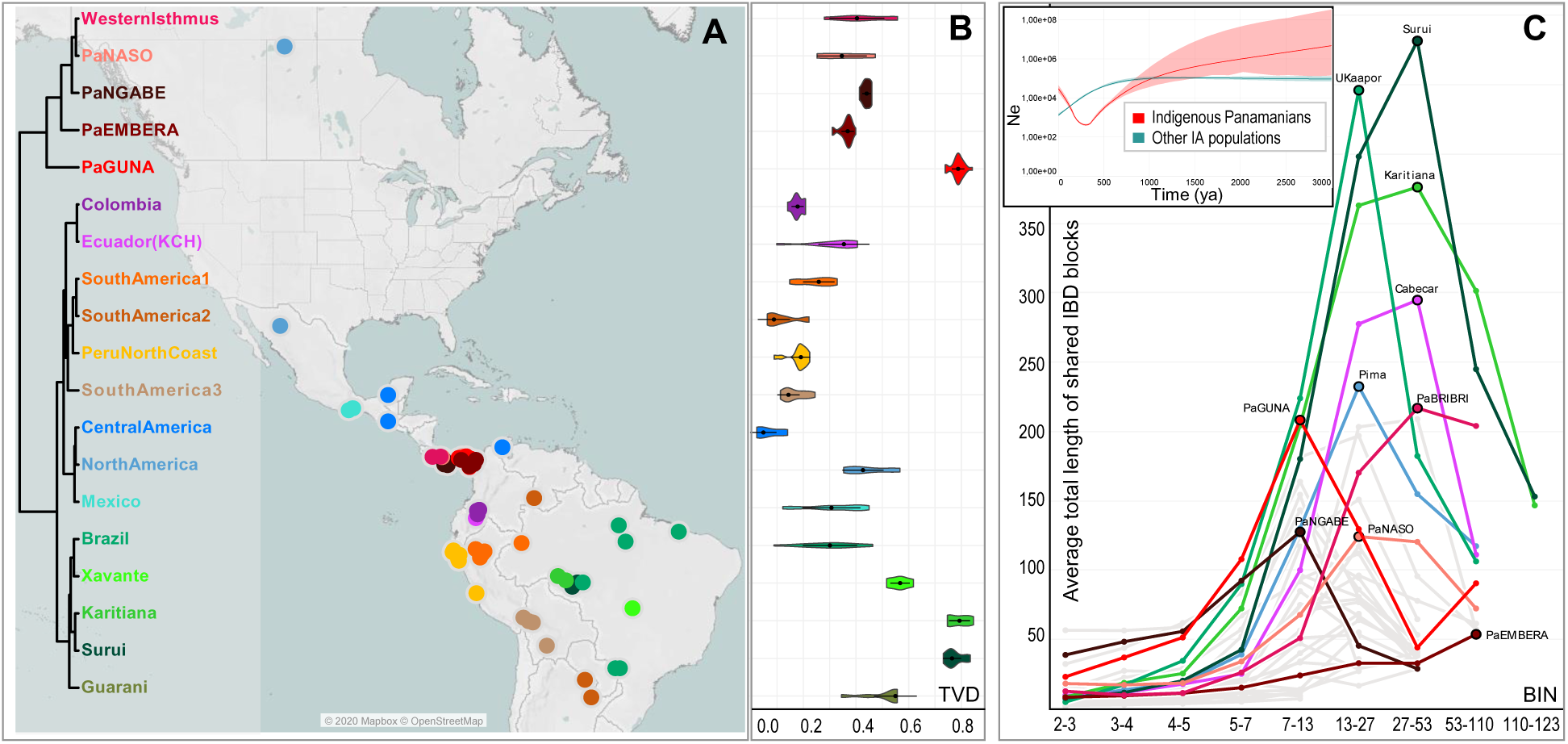
Population Genetic Structure as Revealed by Haplotype Analysis of Modern Panamanian and Indigenous American Populations. (**A**) FineSTRUCTURE dendrogram showing the 19 identified Indigenous clusters and the geographic distributions of the individuals in the nearly unadmixed Indigenous American (uIA217) dataset. (**B**) Violin plot showing cluster self-copy lengths (fragments copied from members of their own cluster) in the uIA217 dataset; higher values are for more isolated groups. (**C**). Density of the IBD segments by length of bins (chunks or artificially created genome fragments) in the Panamanian and non-Panamanian Indigenous groups of the uIA217 dataset. The inset shows the estimation of changes in effective population size (Ne) over time based on IBD segments. A drastic reduction of Ne was experienced by Panamanians since approximately one thousand years ago, considering a generation time of 25 years.

### Deciphering genomic connections outside the Isthmus

Previous studies (Gnecchi-Ruscone et al., 2019; Moreno-Estrada et al., 2013; Reich et al., 2012) have already provided hints of genetic patterns in the Isthmo-Colombian region, eventually extended to other groups that speak Chibchan languages. Here we first confirmed that the previously-discussed Isthmian component is also detectable in additional Chibchan-speaking populations genotyped with a different array (Arhuaco and Kogi from Colombia; Guaymí, Cabécar, Teribe, Bribri, Huëtar, and Maleku from Costa Rica) and that its highest legacy can be detected in the eastern Isthmian land-bridge consisting of the present-day countries of Panama and Costa Rica (Figure S17). We have now detailed patterns of genomic variation of the area’s core population(s), represented by pre-Hispanic and modern Panamanians, underlining their unique features in the overall genetic landscape of the Americas. The *f-statistics* tests detected higher levels of shared genetic history between ancient Panamanians and the present-day Isthmian area extended to Costa Rica (Cabécar) and northern Colombia (CLM and Wayuu) in comparison to other ancient and modern populations (Figures 4, S16B, S18).

**Figure 4.**
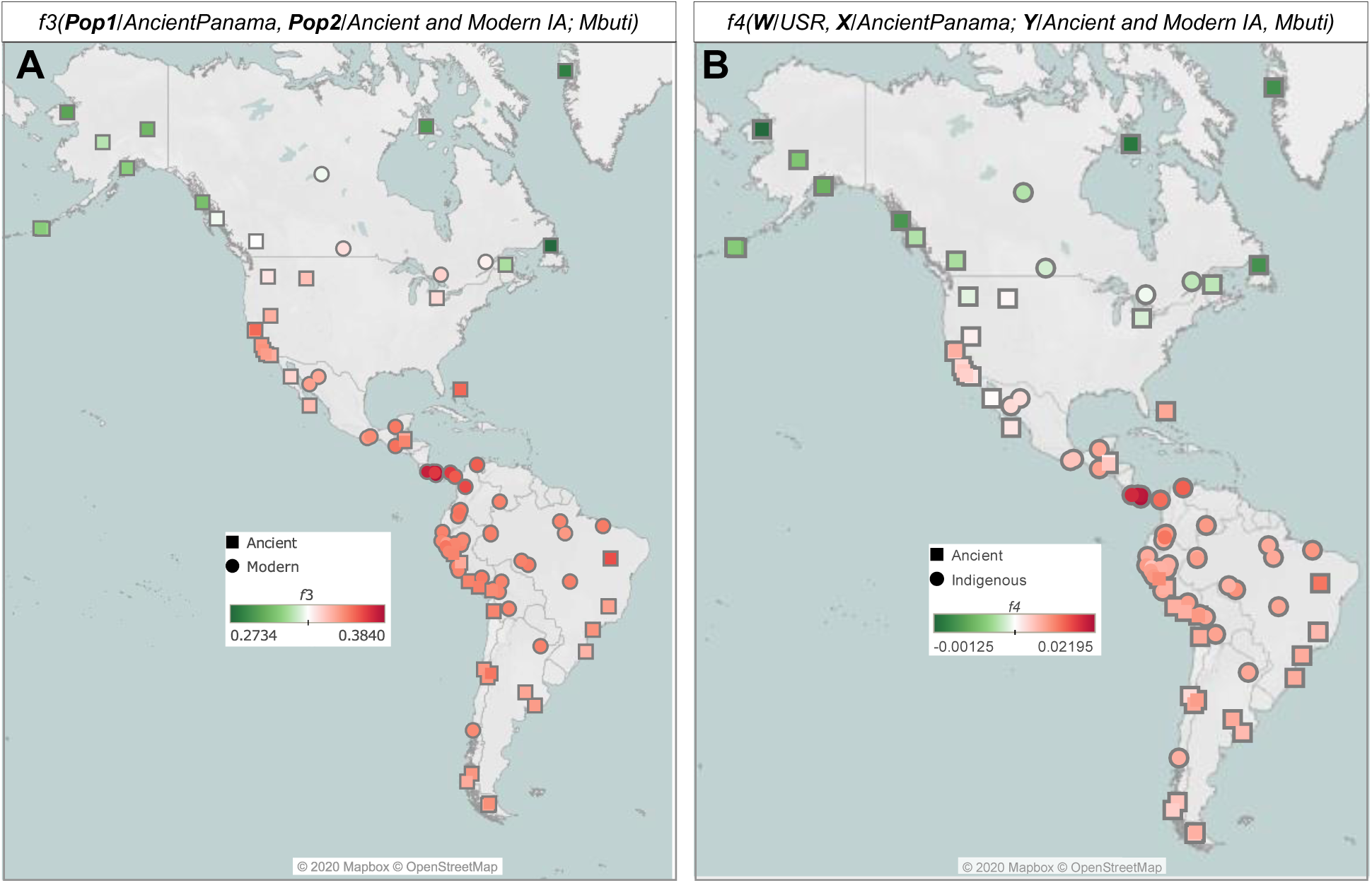
*f-statistics* of Ancient Panamanians in Comparison to Indigenous American Groups. (**A**) Outgroup *f3-statistics* on the uIA89 plus mIA417 dataset (only transversions) in the form *(AncientPanama, Ancient and Modern IA; Mbuti)* showing the longer shared genetic history among Isthmo-Colombian populations. (**B**) Same results confirmed with *f4-statistics* in the form *(**W**/USR, **X**/AncientPanama; **Y**/Ancient and Modern NA, Mbuti)*. Each tested population (Y) is represented by squares (Ancient) and circles (Modern).

An early Isthmo-Colombian branch has been also detected in the neighbor-joining tree (based on outgroup *f3-statistics* and rooted with an ancient Beringian genome, Figure S19) of ancient and modern IA groups. This graph identifies different branches (with a geographic pattern) that largely overlap with the haplotype-based clusters identified in the American-wide uIA217 dataset (Figure 3A). In the latter dendrogram, an Isthmian macro-branch is well separated from the one that encompasses all other IA populations. An ancient origin of the Isthmo-Colombian distinctiveness is suggested by our analysis of effective population size based on the identical-by-descent (IBD) segments. A drastic reduction of the Panamanian population size probably began during pre-colonial times (∼1 kya, inset Figure 3C), thus before the one experienced by other IA population groups. This trend is mainly driven by the Guna, suggesting that their demographic contraction not only preceded that of other Indigenous groups but predated European contact (Figures S20, S21). It is plausible that the current gene pool of the Guna was shaped by a more ancient isolation event than those experienced by the other Indigenous populations (in and outside the Isthmus), as suggested by the observed enrichment of shorter IBD segments, around the 7-13 Mb range for the Guna and Ngäbe, while the peaks of other isolated Indigenous groups are for longer fragments (Figure 3C). The comparison of the IBD fragments shared between our Panamanian groups and other IA populations (Figure S22) reveals ancient (since at least 2,500 years ago) interactions within the Isthmo-Colombian area, much stronger and temporally extended among the western populations (Cabécar, Bribri and Naso), currently living in geographic region associated with the pre-colonial Greater Chiriquí cultural area. On the eastern part of the isthmus, the Guna also show a number of short (older) blocks shared with the Maya from Mexico, while the Emberá share shorter blocks with South American populations. Thus, the former probably received ancient genetic inputs from the north, while the latter admixed with external southern sources (Figure S22). The Guna also show a direct connection with the ancestors of North and Central American populations in the TreeMix ML tree when two migration edges (gene flows) are added (Figures 5A, S23). Such an ancient legacy is also confirmed by the Panamanian mtDNA tree. The most represented haplogroups among ancient and modern Panamanian mitogenomes belong to the four main pan-American founding lineages (A2, B2, C1 and D1; Figure 5B). We also identified four Isthmo-specific sub-branches, the most represented one (A2af1) is dated at 15.82±4.09 kya (Figure S7). Finally, the BSP plot of Panamanian mtDNAs shows an increase in population size starting in the early Holocene (∼10 kya).

**Figure 5.**
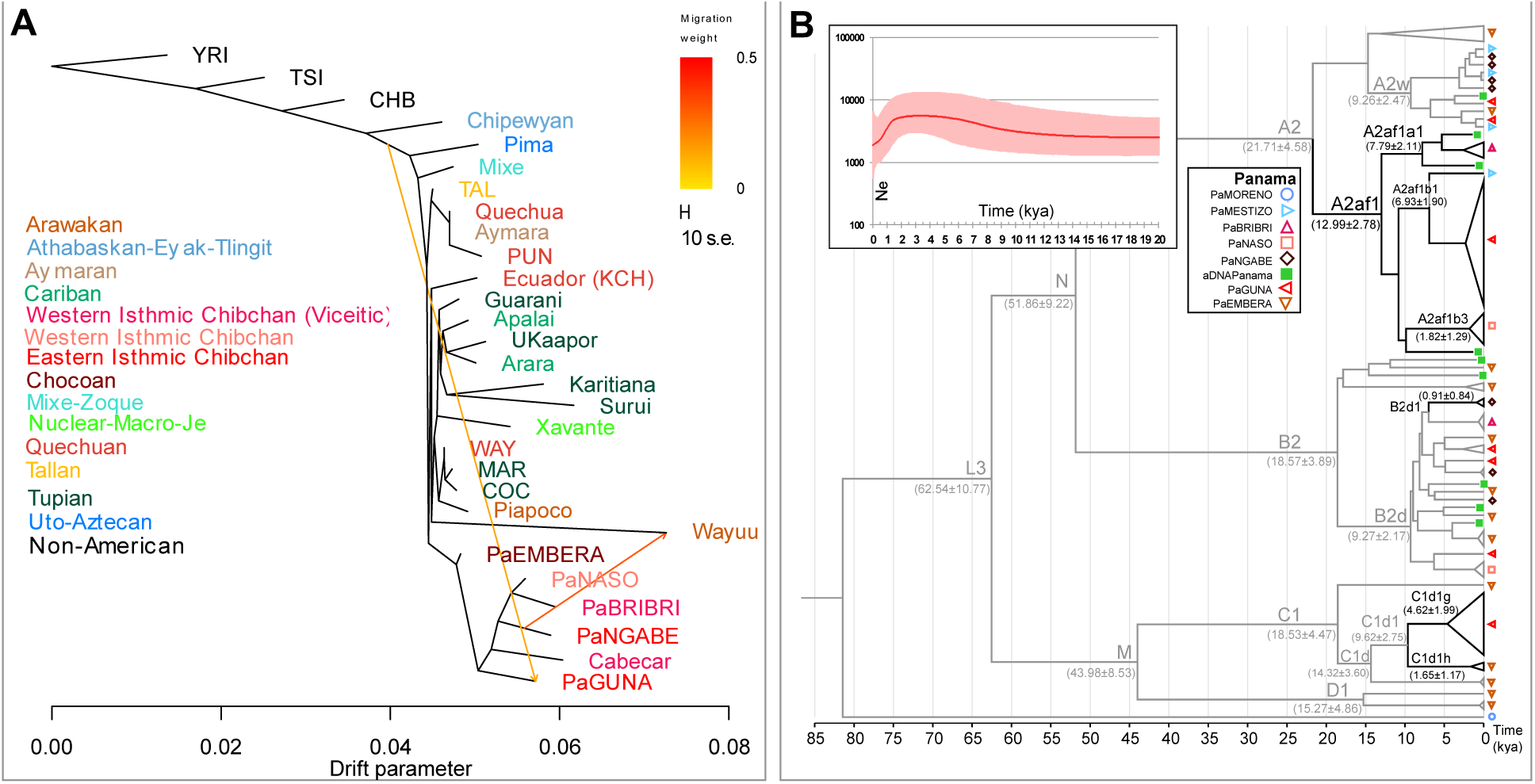
Schematic Phylogenetic Trees Based on Genome-wide and MtDNA Data. (**A**) Inferred Maximum Likelihood tree built with TreeMix on the unadmixed dataset uIA89 allowing two admixture edges (migration events). Population groups are colored according to linguistic/geographic affiliation. Horizontal branch lengths are proportional to the amount of genetic drift that has occurred on the branch. Migration arrows are colored according to their weight. (**B**) Bayesian phylogenetic tree of ancient and modern mitogenomes from Panama belonging to founding haplogroups. It was rooted on an L2c2 mitogenome from a “Moreno” individual. The Bayesian age (mean value with standard deviation) is shown for relevant branches. Black lines highlight Isthmo-Colombian-specific branches. The inset shows the Bayesian Skyline Plot (BSP) displaying changes in the effective population size (Ne) through time.

### A previously undescribed ancestry among ancient Indigenous peoples of the Americas?

To further understand the peculiarities of the Isthmo-Colombian populations within the context of the most updated archaeogenomic scenario of non-Arctic America (see Introduction), we employed *f4-statistics*, which control for possible biases deriving from population-specific drift, to compare ancient individuals and contemporary Indigenous groups to our individuals. As expected, the Isthmus shows an excess of allele sharing with modern and ancient Indigenous populations from Central and South America when compared to Ancestral Beringia (USR, Upward Sun River, Alaska, ∼11.5 kya) and Northern Native American (NNA, represented by ASO, Ancient Southwestern Ontario, ∼4.2 kya) genomes, but this picture is more intricate when dealing with the Southern Native American (SNA) related ancient genomes (Figures S24-S28). The affinities between the (Y/tested) Isthmian populations and other Indigenous (X) groups are significantly stronger in relation to Anzick-1 (Montana, ∼12.8 kya) than to Spirit Cave (Nevada, ∼10.9 kya) (W individuals; Figure 6A), suggesting that Isthmian populations are related to Spirit Cave as much as to other indigenous groups, while Anzick-1 is an outgroup to SNA indigenous groups. Moreover, the Isthmus seems more closely related to Spirit Cave than to Anzick-1 in comparison to Ancestral Beringia (Figure S28). We directly tested the relationships of Isthmian and other Central/South American populations with Anzick-1 and Spirit Cave highlighting a differential trend that becomes significant with a higher molecular resolution power, i.e. more SNPs (Figures 2C, S29).

**Figure 6.**
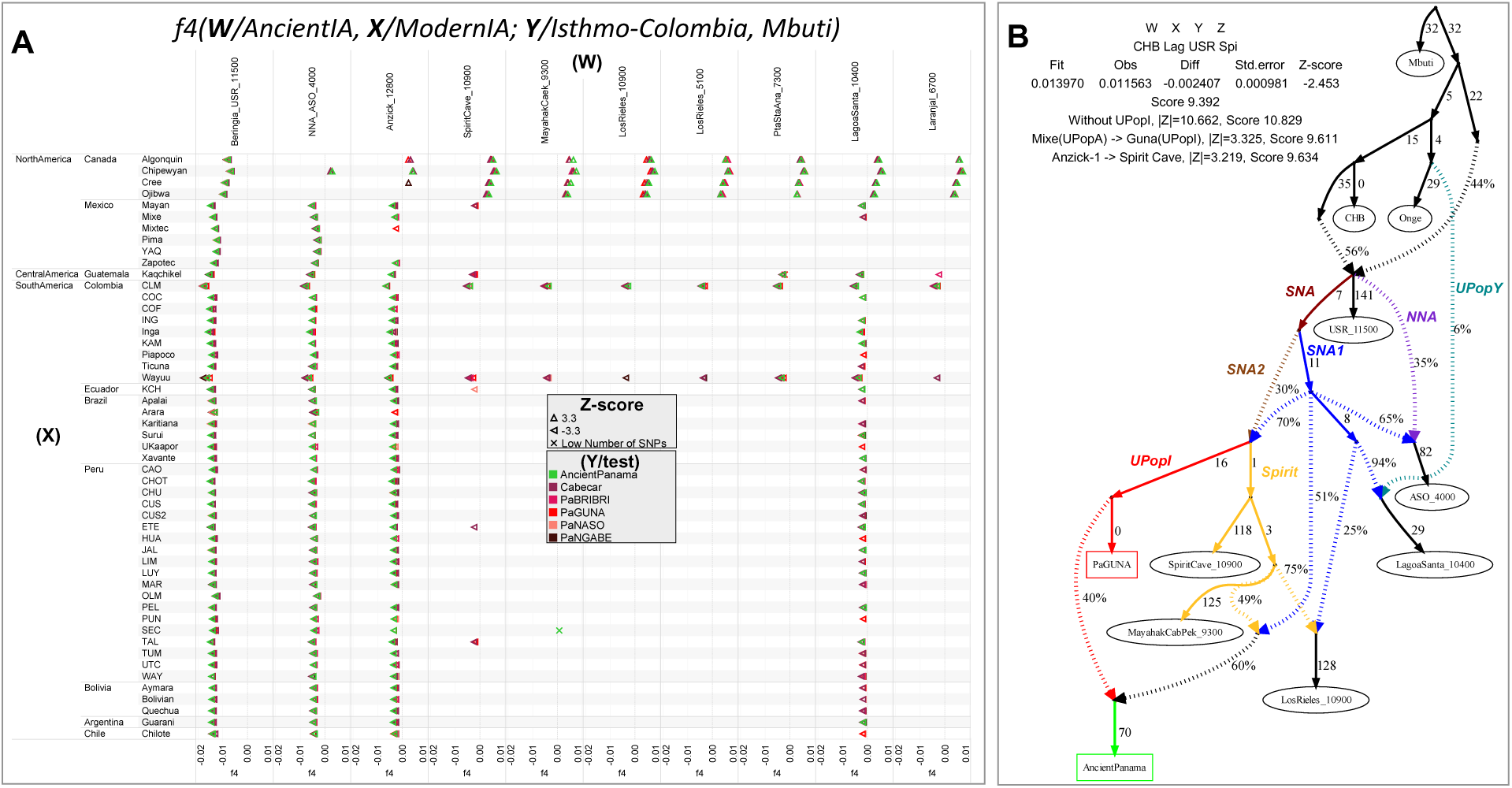
*f4-statistic* Tests and Admixture Graph Modelling Ancestries and Affinities of Isthmian Groups in America. (**A**) *f4-statistics* in the form *f4(**W**/modern NA, X/ancient NA; **Y**/Isthmus, Mbuti)* on uIA89, mIA417 and ancient genomes considering only transversions. Only sub-groups of meaningful ancient genomes were considered (see Figures S24-S28 for comparisons with the entire ancient dataset). The Emberá group was excluded due to its previously-demonstrated admixed origin. Each tested population (Y) is shown (with triangles pointing to X or W population) only when the initial conformation of the tree is rejected (*p∼0.001*, depending on the Z-score), thus visually pointing to the closer population (W or X) in each comparison. (**B**) Best fitting *f-statistics*-based admixture graph optimized using qpGraph. We modelled the genetic history of ancient Panamanians and the Guna directly linked to ancient Indigenous American genomes representative of the SNA ancestries. At the top, we show the four populations leading to the worst Z-score after optimizing the model. Numbers to the right of solid edges represent optimized drift parameters and percentages to the right of dashed edges represent admixture proportions. Different colors indicate the specific ancestries discussed in the text.

Such genomic differences are confirmed when moving southward in Central America (Figure S25) and particularly for the early ancient genomes excavated in the southern continent. The Pacific coast populations (Los Rieles, Chile, ∼10.9 kya, Figure S26) exhibit greater affinity to Spirit Cave, while the ancient genomes from the Atlantic side show the same pattern as Anzick-1 when considering individuals older than ∼7 kya (Lagoa Santa, Brazil, ∼10.4 kya, Figure S27). These distinctive signals persisted up to about 7 kya, when they were probably erased by a major population turnover in South America (Moreno-Mayar et al., 2018; Posth et al., 2018), facilitated by a widespread population decline due to mid-Holocene climate changes (Riris and Arroyo-Kalin, 2019).

The results of previous analyses revealed that Isthmian and non-Isthmian IA populations are differentially related to available Pleistocene individuals, suggesting the contribution of different sources. To test if the Isthmian and non-Isthmian groups derived from the same or distinct ancestral populations, we used *qpWave* (Patterson et al. 2012), which estimates the minimum number of sources necessary to explain the observed genetic composition of population groups. Significance values are consistent with pairs of Isthmian and non-Isthmian groups deriving from at least two separate streams of ancestry, as attested by Rank1 *p-value* <0.01 in most comparisons, especially for the Guna (Figure S30). This finding demonstrates that the distinctiveness of the Isthmo-Colombian area cannot be explained by genetic drift alone, as recently inferred in other population contexts (Nägele et al., 2020). The Guna also show lower values (mostly <25^th^ percentile) of shared genetic history with ancient genomes representative of well-known Indigenous ancestries than the average of the one shared by other IA populations (Figure S31). Therefore, we modelled admixture graphs looking for the most plausible origin of the ancestral source(s) of the Isthmian component (Figures S32-S38; see also the specific section of Supplementary Information for further details). The best supported topology successfully tested the hypothesis that the ancestral gene pool of the Isthmo-Colombian area, here represented by pre-Hispanic Panamanians, derives from a local admixture between different ancestries (Figure 6B). One derives from the differential mixture of two ancestries, SNA1 and SNA2, which in turn stem from an ancestral SNA source. This scenario is strongly suggestive of the first split between SNA and NNA occurring in Beringia, thus further north than generally proposed (Waters, 2019).

The NNA ancient individual in Figure 6B, ASO (Ancient Southwestern Ontario, ∼4.2 kya), results from an admixture between NNA and SNA1. We could not identify an unadmixed proxy for the NNA ancestry among the available ancient individuals (Figures S34, S36), but NNA does not seem to be involved in shaping the Isthmian genomic pool. Founding populations carrying the SNA1 ancestry probably took part in an early peopling of the double continent passing through the Isthmus and leaving signals on both sides of South America as attested by two of the most ancient genomes, Lagoa Santa in Brazil (∼10.4 kya) and Los Rieles in Chile (∼10.9 kya). The former also confirms a few traces of UPopY, the unknown population of Australasian ancestry, which was previously proposed to have contributed to the early peopling of South America (Moreno-Mayar et al., 2018; Skoglund et al., 2015). On the other hand, only Los Rieles shows significant inputs of the SNA2 ancestry, which moved later or slower than SNA1 along the Americas admixing multiple times with the first settlers along the way, as demonstrated by the ancient admixed genomes of Spirit Cave in North America, Mayahak Cab Pek (Belize, ∼9.3 kya) and our ancient Panamanians in Central America. Once SNA2 reached South America, it probably left a stronger contribution on the Pacific side, as suggested by Los Rieles and supported by the differential pattern depicted by the *f4-statistics* (Figure 6A). However, to fully explain the genetic variation of pre-Hispanic Panama, we need to consider an additional ancestry represented by a still Unsampled Population of the Isthmus (UPopI, Figure 6B), which parallels the Spirit Cave branch of SNA2 and left its strongest traces in the contemporary Guna. This best-fitting model was also checked without considering UPopI and replacing Spirit Cave with Anzick-1 and Guna with Mixe, previously used to identify UPopA (Moreno-Mayar et al., 2018), without finding any statistically supported graph (Figure 38C). Therefore, it is supported that Anzick-1 and Spirit Cave might represent different ancestries and UPopI is a still unsampled population distinct from UPopA. UPopI likely originated in the north during the late Pleistocene, as attested by the age (∼15 kya, Figure S7) of the entire mtDNA haplogroup A2af1 that probably represents its mitochondrial legacy, and expanded more than 10 kya in the Isthmo-Colombian area (according to mitogenome data; inset of Figure 5B). This scenario would explain the ancestral component, already seen in the ADMIXTURE analyses, which is geographically restricted to the Isthmo-Colombian area and prevalent in the Guna, where it was probably maintained by a high-level of isolation in antiquity.

## CONCLUSIONS

Our work enriches the Indigenous genomic database with autosomal data from cultural groups of Panama and ten low-coverage pre-Hispanic genomes obtained from human remains excavated in the tropical area of Panama City near the sea. The ancient genetic profiles from *Panamá Viejo* and *Coco del Mar* sites, radiocarbon dated from 603 to 1430 CE, confirm similarities in the gene pool of this pre-Hispanic population(s), suggesting common origins. The diachronic comparison with population groups presently living in Panama allowed us to identify genetic similarities with the Guna and Ngäbe and suggestive connections with some admixed individuals, implying that, in the wake of the conquest, there was extensive gene flow. A genetic substructure has been identified in the entire Isthmo-Colombian region, with a macro-group encompassing Cabécar, Bribri, Naso and Ngäbe, who currently live in the pre-colonial Greater Chiriquí cultural area. A wider genetic variation characterizes eastern Panama, here represented by pre-Hispanic individuals from *Panama Viejo* and *Coco del Mar* plus Guna, Emberá and northern Colombians. Our analyses suggest that pre-Hispanic demographic changes and isolation events, evident and more ancient in the Guna, contributed to create the genetic structure currently seen in the region. Moreover, through allele frequency analyses and haplotype-based reconstructions, we describe the presence of a novel axis of Indigenous genetic variation in the Americas, which is specific to the Isthmo-Colombian area and possibly extended to other Chibchan-speaking groups. This component was present not only among pre-Hispanic Panamanians, but also strongly characterizes current Isthmian groups, surviving both pre-colonial demographic fluctuations and the genetic bottleneck (and admixture) caused by colonialism.

The detection of this component has an impact that expands far beyond the Isthmo-Colombian area and the ancestry of its past and current inhabitants. The following clues point to the scenario that it arose in the late Pleistocene: (i) the pre-Clovis age of the Isthmian-specific mtDNA haplogroup A2af1, (ii) the internal structure that emerges when only the Indigenous genome-wide variation is analyzed, (iii) the longer shared genetic history among Isthmo-Colombian populations with respect to other Indigenous populations and (iv) the differential relationships with Pleistocene individuals from North America. Then, to identify its ancestral source(s), we built a statistically significant model that explains this Isthmo-Colombian component as a local admixture of different ancestries of northern North American origin. At least two SNA ancestries, SNA1 and SNA2, differentially associated to available Pleistocene genomes, should be considered, as well as an additional Isthmian-specific ancestry. The latter requires the contribution of UPopI, which stemmed from the same source (SNA2) that contributed to the pre-Clovis groups with Western Stemmed technologies associated with Spirit Cave (Figure 6B) and, according to mitogenome data, expanded within the Isthmo-Colombian area during the early Holocene.

The ancestral admixtures described here were probably bound to now-submerged archeological sites on the Pacific coast of the Isthmus. Nevertheless, the genomes of the pre-Hispanic individuals from *Panama Viejo* and *Coco del Mar* attest to these events, and the site of Vampiros-1 (initially named *Cueva de los Vampiros*), the only Pleistocene site on the lower Isthmian land-bridge that contains cultural but not human skeletal remains, provides further archeological support. Vampiros-1 shows evidence of both Clovis and Fluted Fishtail Point lithic traditions indicating that hunter-gatherers of extra-Isthmian origin were on the lower Isthmus 13.2-11.7 kya with specific composite weaponry and cutting/scraping tools (Ranere and Cooke, 2020). Our model also fits well with recent archeological records from both sides of the Isthmo-Colombian area. Archeological findings in southern North America report early occupations as far south as central-northern Mexico around the LGM (Ardelean et al., 2020) and more widespread settlements in warmer pre-Clovis times (14.7–12.9 kya) (Becerra-Valdivia and Higham, 2020). The cultural heterogeneity observed among the oldest reliable pre-Clovis archaeological sites of South America (dated 15.1-14.0 kya) along the Pacific coastal zone (Huaca Pietra in Central Andes; Monte Verde II in South Andes) (Dillehay et al., 2017) and in the Pampas (Arroyo Seco 2) (Politis et al., 2016) can be explained considering a deeper chronological time (between 16.6 and 15.1 kya) for the Isthmian crossing that led to the initial peopling of South America (Prates et al., 2020).

The preservation effect of an ancient legacy in “outlier populations”, such as the Guna, is not a novelty in population genetic studies. Such behavior recalls that of Sardinians and Basques in Europe (Achilli et al., 2004; Chiang et al., 2018; Novembre et al., 2008; Olivieri et al., 2017; Palencia-Madrid et al., 2017). In the European context Sardinians maintained the most evident traces of the early European Neolithic farmers (Lazaridis et al., 2016; Lazaridis et al., 2014; Raveane et al., 2019). Among Indigenous peoples, some Amazonian tribes, which match the very high internal similarities of the Guna, have preserved the specific ancestry of an unknown population linked to Australasia (Moreno-Mayar et al., 2018; Skoglund et al., 2015). Certainly, additional high-resolution genomic data (from early Holocene to colonial times) are needed to provide definitive answers to these and other questions concerning the Isthmo-Colombian crossroad by detailing the genetic patterns that we have identified in the Panamanian population(s) and by reconstructing variation in population sizes over time.

## ACKNOWLEDGMENTS

We thank Nicole Smith-Guzmán for her suggestions and comments and Sandra Kraus, Susanne Lindauer, Robin van Gyseghem, and Matthias Hänisch for processing and analyzing radiocarbon samples at the CEZA in Mannheim.

We are grateful to the Patronato Panamá Viejo, the Ministerio de Cultura de Panamá and all volunteers who generously participated in this survey and made this research possible.

This research received support from: the European Research Council (ERC), Horizon 2020 Consolidator Grant CoG-2014, No. 648535 (to B.A., J.R., I.H.-M., C.K., J.G.M., A.A.); the University of Pavia – INROAd program (to A.A.); the Fondazione Cariplo project no. 2018–2045 (to A.O., A.T., A.A.); the Italian Ministry of Education, University and Research (MIUR) for Progetti PRIN2017 20174BTC4R (to AA) and Dipartimenti di Eccellenza Program (2018–2022)

- Department of Biology and Biotechnology “L. Spallanzani,” University of Pavia (to A.O., A.T., O.S., A.A.) and Department of Biology, University of Florence (to M.L., D.C.); AIRC IG20109 (A.R., F.B). GH is supported by a Sir Henry Dale Fellowship jointly funded by the Wellcome Trust and the Royal Society (Grant Number 098386/Z/12/Z). The ancient DNA and modern population genetics lab at the University of Tartu is supported by the European Union through the European Regional Development Fund (2014–2020.4.01.16–0030) (C.L.S., L.P., L.O., F.M. and M.M.) and the Estonian Research Council (PRG243) (C.L.S. and M.M.).

## AUTHOR CONTRIBUTIONS

J.R ., T.M., I.H-M., J.G.M. excavated archaeological sites, provided or acquired ancient samples, and contextualized the archaeological findings; M.T., U.A.P performed modern sample collections supervised by J.M.P., J.M., O.S., A.A.; R.F., C.K. produced and analyzed radiocarbon data; M.R.C., H.L., A.M. performed ancient DNA laboratory work supervised by M.L., D.C., R.S.M., A.A., M.R.C., A.R.; A.G-C. produced modern genome-wide data supervised by A.S., A.A.; M.R.C., A.R., N.R.M., G.C., L.O., C.L.S., M.L. analyzed ancient and modern genomic data supervised by F.M., A.A.; M.R.C., N.R.M., G.L., G.G. produced and analyzed mitogenome sequences supervised by A.O., A.T., A.A.; M.R.C., A.R., G.C., V.G. classified Y-chromosome data supervised by O.S.; G.H., F.B., C.C., L.P., M.M. provided technical and computational resources; M.R.C., B.A., A.A. wrote the original draft with major inputs from R.C., R.S.M., A.T. and J.G.M.

All authors discussed the results and contributed to the final manuscript.

## DECLARATION OF INTERESTS

The authors declare no competing interests.

## SUPPLEMENTARY INFORMATION

### EXPERIMENTAL MODEL AND SUBJECT DETAILS

#### Ancient sample collection and archaeological information

Twenty ancient individuals (21 specimens in total) were collected from seven different archaeological excavations (six of these within today’s *Patronato Panamá Viejo*, and one at nearby *Coco del Mar*) (Table S1, Figure 1A). Based on the style of the ceramics recovered, archaeologists consider the area related to contiguous contemporary settlements as part of an extended pre-Hispanic occupation of Cueva-speakers within the Greater Darien cultural region (Martín, 2002b). This population coexisted and mixed with Spanish settlers from 1519 to 1541 Common Era (CE), as an Indigenous presence among Old Panama’s earliest Christian burials suggests. Six of the excavations sampled took place in Panamá Viejo, the site of the colonial city from 1519 through 1671 CE and an area of pre-Hispanic occupation before that.

##### 1. Plaza Mayor (N=4)

The burial site excavated in the Plaza Mayor, originally identified as *Tumba 1* (Figure 1B), contained the remains of a female individual with a spondylus necklace and surrounded by offerings that included nine male crania. Scholars have awaited genetic research in order to test the hypothesized relationships regarding the individuals buried within this tomb since its discovery in 1996 (Mendizabal, 2004). DNA was successfully extracted from the main individual (PAPV109) and three of the nine skulls (PAPV114, 117 and 118) around her. PAPV109 did not meet quality standards to be included in autosomal analyses (Table S1).

##### 2. Plaza Casas Oeste (N=1)

In the Plaza Mayor’s Casas Oeste, the remains of 35 pre-Hispanic individuals were found in different positions, including extended burials, as well as urns and packages of bones. DNA of one of these individuals (PAPV137), radiocarbon dated 898-1014 (2 sigma), a male individual of at least 15 years at death, extended yet leaning toward the right side, with his skull pointed northwest, was extracted in this work.

##### 3. Catedral (N=5)

Human remains excavated from the Old Panama’s Cathedral (PAPV52, 53, 57) and its courtyard (PAPV61, 93) in 2000 reflect the African and European presence, as well as a mixed Indigenous inheritance, in the city from 1542 to 1671 CE (Hernández Mora et al., 2020).

##### 4. Sur de la Plaza (N=2)

Two post-contact individuals analyzed here were excavated from the Sur de la Plaza (PAPV26, 27) and dated 1519-1541 CE, based on the historical and archaeological evidence.

##### 5. Parque Morelos (N=4)

Other pre-Hispanic burials were excavated from the Parque Morelos (roughly 1 km to the west of the Plaza Mayor) in 2001-2003. These excavations uncovered the remains of two pre-Columbian residential structures, including evidence of post-holes, pottery, grinding stones, seashell beads, fragments of three bone flutes, and a frog-shaped gold pendant. Within a few meters from the residential structures, urns and “packages” with cranial and long bones belonging to children and adults (PAPV146 and 156, dated 767-966 and 1264-1289 CE, respectively, and the first one with DNA extracted from two different bones) were also recovered (Martín, 2002a; Martín, 2006). An additional individual was excavated in an extended primary burial (PAPV167, 779-985 CE, 2 sigma) some 50 meters from the residential structures.

##### 6. Centro de Visitantes (N=1)

Another pre-Hispanic individual was extracted from an extended burial excavated by Juan Martín in 2001 at the Centro de Visitantes site, close to Parque Morelos, in the west part of Panamá Viejo (PAPV106), and dated 897-986 CE (2 sigma). Five ceramic offerings accompanied this female individual, of approximately 30-35 years at death.

##### 7. *Coco del Mar (N=4)* located outside Panamá Viejo

The additional site where pre-Hispanic human remains were uncovered in 2005 was in an adjacent residential area, *Coco del Mar*, one kilometer to the west of the *Morelos* statue. In this case, the burial is characterized by a particular conformation (Figure 1B), where a primary, female individual (PAPV172) was discovered with three male crania (PAPV173, 174, 175) and additional pottery as offerings (Martín et al., 2008), similar to that of Plaza Mayor’s *Tumba 1*.

#### Modern DNA sample collection

A total of 84 biological samples were collected in Panama from healthy individuals belonging to different Indigenous groups using NorgenBiotek kits (Table S2, Figure 1A). During sample collection, genealogical information (for at least two generations), cultural affiliation, and spoken language were also gathered from each subject.

#### Insights into pre-Hispanic Panama: details on archaeology, history, and linguistics

From c. 4.5 kya agriculture along Panama’s Pacific watershed was complemented by fishing in rivers and estuaries, and catching birds, large snakes and iguanas (*Iguanidae spp.*), and hunting mammals with body masses <55 kg on the offshore Pearl Island archipelago (Cooke et al., 2016). Some of these, e.g. white-tailed deer (*Odocoileus virginianus*), raccoons (*Procyon lotor*) and rodents including agouti (*Dasyprocta punctata*) and paca (*Cuniculus paca*), forage in human trash and eat crops in gardens and fields (Cooke et al., 2007, 2008; Martínez-Polanco et al., 2020). On *Pedro Gonzalez* (Pearl Islands), a very small deer (*Mazama sp*.) (7-10 kg) was extirpated by hunting. Dolphins (*Tursiops and Delphinus*) were consumed 6.2-5.6 kya and were possibly killed when beached (Cooke et al., 2016). Later, with the establishment of villages ca 4-2 kya, community activities diversified, especially regarding the exchange of goods. Hunting strategies now included communal drives in Pacific wooded savannas. Deer meat was stored salted and dried and then served at special feasts (Martínez-Polanco and Cooke, 2019).

Inhabitation of the Caribbean side of the Central American Isthmus (Costa Rica) began in the Late Pleistocene (13.4-12 kya) (Ranere and Cooke, 2020). In the very humid central Caribbean Panama, human activities date back to 5.9 kya when groups crossed the Central Cordillera to collect food and materials not available on the opposite side, e.g., embalming agents.

On the central Caribbean, maize is found in rock shelters with earth ovens from about 3.5 kya. Materials analysis demonstrates that Monagrillo pottery found here was manufactured in the central Pacific watershed (Griggs, 2005; Iizuka et al., 2014). One site in the coastal lowlands of Coclé province (Zapotal, PR-32), which used Monagrillo pottery, consisted of dwellings stratified within a shell-bearing midden accumulated between 4.3 and 3.2 kya. Zapotal has the characteristics of a small village. There are very large numbers of edge-ground cobbles here, which were used at many 8-3 kya sites in Panama for grinding several plant foods including maize and manioc. The abundance of small fish (<300g live weight) that were taken at Zapotal points to the use of tidal traps and weirs in the nearby estuary in order to maximize biomasses (Zohar and Cooke, 2019).

By 3 kya notable differences had arisen in the material culture of a western region (Greater Chiriquí; Chiriquí and Bocas del Toro provinces) and a central region (Greater Coclé; Coclé, Azuero Peninsula and Veraguas provinces). Greater Chiriquí shows material culture, art, genes and language that are broadly consistent (Ulloa, 2017), in distinction to the Greater Coclé or the even more diverse, but less studied, Greater Darién.

Unique ceremonial and religious precincts stand out in both the Chiriquí and Coclé cultural areas, although they are markedly different: Barriles in highland Chiriquí (Linares et al., 1975), and the twin sites of Sitio Conte and El Caño in the Pacific Coclé lowlands. The ceremonial site at Barriles consists of low platforms, boulder petroglyphs, urn burials, large statues depicting one man sitting on another’s shoulders -- in an apparent display of social dominance -- and an enormous maize-grinding stone (metate) showing explicit iconographic connections between maize and human fertility (Linares et al., 1975). Maize, beans (*Phaseolus* spp,), and palm and tree fruits characterize the samples of carbonized plant remains in the middens in domestic areas (Dickau, 2010; Smith, 1980).

The Barriles ceremonial precinct seems to have served many communities located between 1,000 and 2,300 meters above sea level in the shadow of the Barú volcano, which last erupted 0.5 kya (Holmberg, 2007). It is inferred that Barriles was the initial settlement of a cultural group that first entered this highland zone from elsewhere ca 2.8 kya. Friction led to fission. A sector of the colonizing population moved to Cerro Punta and henceforth maintained vacillating relations with its ancestors (“peace, trade, war”). Communion among the entire descent group was not severed. Periodically, perhaps annually, festivals were held at ancestral Barriles. The feats of the founders and supernatural helpers were celebrated. Large quantities of alcoholic beverages (e.g. maize and palm sap) were likely brewed.

The well-studied heritage of Greater Coclé, which reached its apogee at the great ceremonial and burial precincts of Sitio Conte and El Caño 1.5–0.95 kya (Lothrop, 1937, 1942; Mayo Torné, 2015), confirms a continuity of iconography and symbolism on decorated pottery from 2.5 kya until two decades after Spanish conquest in this central region. But whether this area was also linguistically united cannot yet be determined.

Cultural geography becomes even more complex in the Greater Darién area extending from the El Valle Pleistocene volcano to eastern Darién. Historians and most archaeologists agree that by 1,500 CE the eastern region, and possibly much of the central area, was inhabited by speakers of the Cueva language. Historian Kathleen Romoli and linguists Jacob Loewen and Adolfo Costenla have proposed for many years that the group of settlements that at the time of Spanish conquest spoke the “language of Cueva” were not an “ethnic group” but, rather, a collection of settlements that shared the Cueva language as a *lingua franca*. These three researchers also argue that some polities in fact spoke variants of languages in the Chocoan family, especially those on the Pacific side of the Isthmus (Costenla, 2012; Loewen, 1963; Romoli, 1987). Chibchan languages probably derive from a proto-language that coalesced about 10 kya (O’Connor and Muysken, 2014) in a “core area” on the lower Central American Isthmus (southern Costa Rica and western-central Panama). Ever since (Barrantes et al., 1990) conducted an isozyme-based study of modern Central American indigenous polities that spoke languages in the Nuclear Chibchan family (Costenla, 2012), it has been apparent that this population coalesced very early in the Holocene and at the onset of agriculture gradually experienced *in situ* fission and fusion (Linares et al., 1975).

Ceramics found in eastern Panama point to greater proximity to the Greater Coclé tradition than a putative Gran Darién cultural sphere among the peoples who inhabited this region from 200 BCE to 1200 CE. Recent findings on the Pearl Islands archipelago confirm the expansion of the ceramic style known as Cubitá, as well as molded and incised variations of the Conte style, both from the central region of Gran Coclé in the gulf of Panamá (Martín et al., 2016). Biese had already suggested that this expansion reached *Panamá Viejo*, where he reported examples of the Conte style excavated around Puente del Rey, toward the north of the site (Biese, 1964).

At the Miraflores site on the Banks of the Bayano River (Cooke, 1976; Oyuela-Calcedo and Raymond, 1998), 670–1015 CE and 700–1030 CE according to the latest 2 sigma calibrations (Martín et al., 2016), only one piece made in the region exhibited painted decorations with obvious influences from Greater Coclé: a plate with a tripod pedestal with the effigy of a monkey (Cooke, 1998). The same pattern of cultural replacement is documented on the Pearl Islands archipelago, where the islands’ Fifth Ceramic Horizon is identified from 750 through 1350 CE (Martín et al., 2016). This late ceramic configuration, which features incised decoration and molding procedures different from its immediate precedents, is that which archaeologists normally associate with Greater Darien, the region that sixteenth-century Spanish chroniclers described as populated by communities that spoke the “Cueva language” from Urabá to the eastern slope of the El Valle volcano (Martín, 2002b; Romoli, 1987).

Martín and colleagues (Martín et al., 2016) argue that the discrepancy between the Greater Darien and Greater Coclé cultural areas, whose geographical extension may have shifted over time, could derive from a change in the population inhabiting the Pearl Islands archipelago: the group using Cubitá ceramics and Conte variations may have ceded before the entry of a population with a very different ceramic tradition from the Darien region and related to northwest Colombia. Another hypothesis (Cooke, 1998; Martín and Sánchez, 2007; Sánchez-Herrera and Cooke, 2000) relates the changes observed to a reorganization of the commercial routes and exchanges from 500 CE, which intensified after 800-900 CE, with the introduction of metallurgy to the isthmus and the replacement of shell artifacts with those of gold to represent high social status. From at least the beginning of the common era through the end of its first millennia, the dispersion of Cubitá ceramics allude to the fact that the Pearl Islands archipelago, the Azuero Peninsula and the central coast of the bay of Panama participated in the same sphere of social interaction. This relationship changed completely in the subsequent period, and until European contact.

### METHODS DETAILS

#### Collagen extraction and ^14^C-dating

The individuals PAPV106, PAPV109, PAPV114, PAPV117, PAPV118, PAPV137, PAPV146, PAPV156, PAPV167, PAPV172, PAPV173, PAPV174 and PAPV175 were analyzed by accelerator mass spectrometry (AMS) at Mannheim (Table S1; Figure S1C), confirming the archaeological context. This time span can be used to assess the Indigenous population’s alleged continuity and cultural practices. extinction

Compact bone portions were cut and the surfaces removed. About 1 gram of sample was placed in glass tubes, demineralized in 10 ml of 0.5 N HCl at initially 4°C and later at room temperature for 14 days, rinsed to neutrality and reacted with 10 ml of 0.1 M NaOH for 24 h at 4°C, rinsed again to neutrality and gelatinized in 4 ml of acidified H_2_O (pH 2-3) for 48 h at 75°C. Insoluble particles were separated using EZEE filter separators. Ultrafiltration (molecular mass > 30 kD) removed the short-chained collagen and concentrated the long-chained collagen, which was frozen and lyophilized.

The collagen was combusted and the relative amounts of carbon (C) and nitrogen (N) determined using an elemental analyzer (Elementar Inc., MicroCube). The produced CO_2_ was reduced to graphite using either a custom-made, semi-automated graphitization unit or a fully automated and commercially available unit (IonPLus Inc., AGE3). The resulting graphite powder was compressed into aluminum targets and subsequently analyzed using a MICADAS-type AMS-system (IonPlus Inc.) (Kromer et al., 2013).

The isotopic ratios ^14^C/^12^C and ^13^C/^12^C of samples, calibration standard (Oxalic Acid-II), blanks and control standards were measured simultaneously in the AMS. ^14^C-ages were corrected for isotopic fractionation to δ^13^C=-25‰ (Stuiver and Polach, 1977) using the ^13^C/^12^C AMS-values and calibrated using the dataset INTCAL13 (Reimer et al., 2020; Reimer et al., 2013) and software SwissCal (L.Wacker, ETH-Zürich). Calibration graphs were generated using the software OxCal (Ramsey and Lee, 2013).

#### Ancient DNA processing

Different bones (femur, humerus and petrous bone) and teeth were available from ancient individuals for DNA extraction, which was carried out in the dedicated clean rooms at the Carl R. Woese Institute for Genomic Biology, University of Illinois, following published protocols (Dabney et al., 2013; Lindo et al., 2017; Pinhasi et al., 2015; Scheib et al., 2018). Bones or teeth were soaked in sodium hypochlorite (bleach 100%) for 3 minutes to remove surface contamination, then washed three times with DNA-free ddH_2_O and once with isopropanol. Dried samples were placed in a DNA Crosslinker under UV. About 0.1 grams of tooth or bone powder were drilled. The powder was incubated in 1 ml of 0.5 M EDTA with 60-100 µl of Proteinase K (33.3 mg/ml) and 50 µl of 10% N-lauryl sarcosine for 20-24 hours at 56°C. The digested samples were concentrated at ∼100 µl using Amicon centrifugal filter units. The DNA was purified using the MinElute Reaction Cleanup Kit (Qiagen). The extracted DNA was quantified with Qubit™ dsDNA HS Assay Kit and tested through a PCR amplification of mtDNA control-region fragments (<150 bps). Samples showing at least 5 ng of DNA and with at least one successful short mtDNA amplification were selected for shotgun sequencing.

Double-stranded DNA libraries were prepared starting from ∼55.5 µl of extracted DNA using the NEBNext® Ultra™ DNA Library Prep Kit for Illumina®. Adapters were used in a dilution of 1:20, which is recommended for low concentration DNA samples. Adaptor-ligated libraries were purified using the MinElute Reaction Cleanup Kit (Qiagen) and then prepared for amplification, which was carried out in thermocyclers in the modern DNA laboratory. The MinElute Reaction Cleanup Kit (Qiagen) was used to clean amplified libraries, whose quality was then checked on the E-Gel Precast Agarose Electrophoresis System and quantified on the Agilent 2100 Bioanalyzer Instrument with the High Sensitivity DNA kit.

Eventually, 21 DNA libraries (10 nM) were selected (two for the individual PAPV146) for Illumina sequencing on the HiSeq4000 (single-end 100 bp, for a total of 150-200M reads) at the Roy J. Carver Biotechnology Center of the University of Illinois.

Mitochondrial DNA capture was also performed at the University of Florence for four individuals (PAPV27, PAPV53, PAPV93, PAPV109), as in (Modi et al., 2020). Libraries were sequenced at the National Neurological Institute C. Mondino in Pavia using an Illumina MiSeq instrument (paired-end reads 150×2).

#### Modern DNA processing

After automated DNA extraction with Maxwell® RSC Instrument in the lab of Genomics of Human and Animal Populations at the University of Pavia, the samples were genotyped with the Affymetrix Human Origin 600K chip at the Institute of Healthcare Research in Santiago de Compostela (CEGEN).

As for the mitogenome sequencing, the entire mtDNA was amplified in two overlapping fragments (Modi et al., 2020). Libraries were prepared using the Illumina Nextera XT DNA preparation kit and sequenced at the National Neurological Institute C. Mondino in Pavia on an Illumina MiSeq instrument (paired-end reads 150×2).

### QUANTIFICATION AND STATISTICAL ANALYSIS

#### Ancient data preparation for analysis

Illumina adapters were removed from raw data using CutAdapt (Martin, 2011). Trimmed FASTQ files were then checked with FastQC (Andrews, 2010). Taking into account the high number of K-mers that were found, mostly poli-A probably generated during the blunt-end phase, CutAdapt was run twice to remove these artefacts. Trimmed reads were mapped against the human reference genome hg19 build 37.1, as well as versus the revised Cambridge Reference Sequence (rCRS) in separate runs, using the algorithm *aln* of bwa v0.6.1 (Li and Durbin, 2010). The numerous duplicates, generated during library amplification, were removed with the tool MarkDuplicates of the Picard package (http://broadinstitute.github.io/picard).

#### Validation tests

##### • Ancient DNA damage pattern

Damage patterns were analyzed with MapDamage2.0 (Jónsson et al., 2013). The molecular decay after death was calculated from the length of the reads, as in (Allentoft et al., 2012). Considering the debate about the possibility of dating the fragments with this algorithm, we only compared the decay of our individuals with three well-known ancient Indigenous genomes (Anzick-1, Kennewick, Taíno). Finally, we used the error rate estimate to confirm the antiquity of our reads. It calculates the excess of derived alleles observed in an ancient genome (compared to a high-quality genome) taking into account that all the anatomically modern humans should have the same percentage of derived alleles. The error rate analysis was performed comparing our trimmed FASTQ with the “Error estimation” tool of ANGSD (Analysis of Next Generation Sequencing Data) (Korneliussen et al., 2014), using the chimp genome as an outgroup and the genome NA12778 (from the 1000 genomes project), as an error free individual (Schroeder et al., 2018).

To double-check if an excess of damage and/or erroneous adapters’ trimming might affect the results of the subsequent analyses, we repeated two AncientPanama fastq trimming using more stringent criteria: removing 10 additional bases from both ends of each read; removing the adapters using a higher mismatch rate (up to 50%) than in the original pipeline (20%). The new trimmed fastqs were then mapped against the reference genome hg19 build 37.1. We repeated the specific Isthmo-Colombian PCA analysis (Figure 2B) including the Ancient Panamanian data trimmed with more stringent criteria (Figure S3A). The almost complete overlapping of the different Ancient Panamanian datasets confirms that neither excessive terminal damage nor missed partial adapter sequences have a detectable effect on SNP chip overlap results. The slight deviation of few samples that undertook aggressive trimming is due to an extensive reduction of the coverage.

#### Contamination tests

##### • Chromosome X contamination

For each library derived from ancient male individuals we estimated nuclear contamination using the approach described in (Moreno-Mayar et al., 2020), based on reads mapping to the X chromosome. This method relies on the fact that males are hemizygous for X-linked loci outside the pseudo-autosomal regions, making multiple alleles in these loci attributable to either errors or contaminations. These estimates can be used as proxy for nuclear contamination estimates in ancient male individuals.

We used ANGSD (Korneliussen et al., 2014) to estimate contamination on reads with mapping quality greater than 30 and base quality greater than 20. We considered at least 100 sites with depth greater than or equal to 2 (Nägele et al., 2020) matching the HapMap CEU allele frequencies (Altshuler, 2010) as potential contaminants, after excluding pseudo-autosomal regions on chromosome X. We then applied the tool presented in (Moreno-Mayar et al., 2020) to estimate contamination using low-depth X-chromosome data, setting to 1,000 the maximum number of jackknife samples used for estimating standard errors and considering the estimates from the Two-consensus method.

##### • Mitochondrial DNA contamination

###### Jones’s Method

Contamination in ancient mitochondrial sequences was first estimated by assessing the number of non-consensus base calls (with base quality greater than or equal to 20) at haplogroup diagnostic positions as a function of the total coverage for each of these sites (Jones et al., 2017).

###### contamMix

We also estimated mtDNA contamination in our data by using contamMix v. 1.0-10, which provides the maximum *a posteriori* probability of the consensus sequence being authentic (Fu et al., 2013). This method is based on the reconstructed mtDNA consensus to estimate contamination, which should not exceed 50% for contamMix to work. First, we built an mtDNA consensus sequence running ANGSD (Korneliussen et al., 2014) and using the parameters -doCounts 1 and -doFasta 2 (majority rule). We retained only reads with mapping quality higher than 30 and nucleotides with base quality greater than 20. Moreover, we filtered for sites with a minimum depth of 5X. Then, we remapped to the newly built consensus sequence only the reads that mapped uniquely to the mitochondrial reference sequence. We used mafft (Katoh et al., 2002; Katoh and Standley, 2013) to align our consensus sequence to a panel of 311 worldwide mtDNA sequences (Green et al., 2008), representing potential contaminant sequences. Finally, we used both the alignment and the remapped reads for contamination estimation with contamMix, running five independent chains for 50,000 iterations. The results were checked by monitoring the Gelman diagnostic (Gelman and Rubin, 1992) to confirm convergence.

###### Schmutzi

The third method to estimate mtDNA contamination was Schmutzi (Renaud et al., 2015), which jointly estimates present-day human contamination and reconstructs the endogenous mitochondrial sequence by considering both deamination patterns and fragment length distributions. Present-day human contamination was evaluated by an iterative likelihood method implemented in Schmutzi using a non-redundant database of 197 human mitogenomes available in the software package.

###### Molecular sex and kinship determination

The sex of each individual was determined using two computational approaches specific for low-coverage genomes, Ry (Skoglund et al., 2013) and Rx (Mittnik et al., 2016) (Figure S4A).

The relationships between the Panamanian ancient samples were verified using the tool READ with default parameters (Monroy Kuhn et al., 2018) (Figure S4B).

###### Checking for reference bias

The reference bias indicates an increased probability of detecting the alleles present in the reference sequence, especially when dealing with paleogenomic data affected by fragmentation and other post-mortem damage, often generating C to T and G to A transitions at 5′ and 3′ fragment ends (Type II damage) (Günther and Nettelblad, 2019). These characteristics might influence mapping scores in low-coverage ancient genomes, particularly for those reconstructed from very short reads, eventually leading to an artificial decrement of reads carrying alternative alleles, in comparison to high-coverage modern genomes. In order to verify that this issue did not affect our downstream analyses, we compared the alternative allele frequency distributions between modern and ancient individual data. We used the PLINK 2.0 toolset to calculate the average proportion of alternative alleles for each individual (Figure S3B). This analysis was repeated considering all SNPs, all SNPs without C to T and G to A and only transversions. As expected, the frequency values are higher for the diploid unmasked dataset due to the presence of heterozygous sites, while the pseudo-haploid datasets (derived from ancient and masked individuals) show overlapping distributions, thus excluding a significant incidence of the reference bias.

#### Modern data preparation for analysis

The raw data were initially checked with Affymetrix suite software, then the data from 81 individuals that passed the quality control (call rate >98.5) (Raveane et al., 2019) were converted into PLINK files and retained for the kinship analyses to exclude putatively related individuals using KING (Manichaikul et al., 2010). Considering that Indigenous populations have a higher degree of relatedness due to long periods of isolation and inbreeding, there was the need to evaluate a new threshold. For this reason, the 97.5-percentile of population IBD for each population was used as the threshold for exclusion (Busby et al., 2015). For each pair of related individuals, the exclusion was based on the number of missing SNPs. A total of 74 individuals (out of the initial 84) were retained after quality and kinship filters.

#### Ancient comparative datasets

The 545,942 SNPs retained in the rWD1,560-dataset (see “Reduced worldwide (rWD1,560) dataset” section) were called on the ancient dataset that contained our 12 ancient Panamanian individuals merged with 241 ancient Siberian and American individuals with a minimum coverage of 0.01X (Table S4). The calling was performed for all individuals in one run using ANGSD (Korneliussen et al., 2014) with the *haplocall 1* option, which picks a random read starting from an input set. In addition, to avoid possible biases due to low coverage data, we down-sampled all ancient genomes to 1X and 0.5X coverage using ANGSD with the *-downSample* option.

The ancient dataset was merged using PLINK 1.9 (Purcell et al., 2007) with our modern datasets and then filtered using --geno and --mind options set respectively to 0.60 and 0.98 (Scheib et al., 2018), keeping only individuals with at least 10,000 SNPs (Table S4) (Posth et al., 2018).

#### Modern comparative datasets

##### Reduced worldwide (rWD1,560) dataset

A first dataset of 4,939 modern individuals was built encompassing worldwide Affymetrix Human-Origins genotyped individuals and American whole-genome sequences from the literature considering a minimum threshold of 500K overlapping SNPs. This dataset was merged using PLINK 1.9 with our modern individuals and then filtered using --geno and --mind options set to 0.2. After excluding related individuals (see “Modern data preparation for analysis” section), the dataset was geographically restricted to 1,560 individuals (rWD1,560, Table S3) including our modern Panamanians (74), all individuals from America (1,084) and Siberia (203), and the western Eurasian (61), African (73) and Australasian (65) populations that left a greater genomic impact on Indigenous Americans during colonial times (Chacón-Duque et al., 2018; Homburger et al., 2015; Montinaro et al., 2015; Ongaro et al., 2019).

##### “Nearly unadmixed Indigenous American” (uIA217 and uIA89) datasets

European colonialism and the African slave trade left a strong impact on the Indigenous American populations. Therefore, the analyses of pre-colonization genetic history might be strongly influenced by these components and it is very difficult to find non-admixed modern individuals, even among Indigenous groups. Therefore, three different approaches (ADMIXTURE, Local Ancestry and *f*4) have been used to create a sub-set of individuals with the maximum possible amount of Indigenous genetic component, to avoid signals altered by recent admixture events. Considering the inconsistency of the preliminary results obtained by using the three methods independently, a stepwise merging approach was preferred retaining individuals classified as Indigenous (with the lowest content of non-Indigenous component) after each step.

*1. ADMIXTURE* (Alexander et al., 2009)

We extend the analyses on our rWD1560 dataset until K20. However, considering that K14 has the lowest cross-validation (CV) error (Figure S10), we used K14 to identify the individuals that have more than 95% of Indigenous components (290 in total).

2. f4 statistics

In the second approach, we used *f4-statistics* in the following form:

*(ancientIndigenous, X; Europe/Africa, Mbuti)*

*Ancient Indigenous* was composed by five high-coverage ancient genomes selected on the basis of country of origin and age (Table S4). The selected individuals (N=305) were those with a Z score < |3| for both Europe and Africa.

*3. Local Ancestry (LA)*

Combining the positive results of ADMIXTURE and *f*4 statistics, we could retain 230 individuals to be selected for the Local Ancestry (LA) using the software RFMix (Maples et al., 2013). Among them, we selected 58 individuals to be used as ancestral source in LA analysis. The overall criteria used to select these 58 ancestral individuals were as follows: i) successfully passing the *f4* filter (see “*f4* statistics” section); ii) 100% NA in K14 (see “ADMIXTURE” section); iii) belonging to a population that best represents a specific NA component in the ADMIXTURE analysis (Figure S10) for each K (until K20):

*o K1 6 Puno individuals (selected at K20 among the population with more K1)*
*o K6 25 Guna individuals (selected at K14)*
*o K10 2 Chipewyan individuals (selected at K14)*
*o K11 10 Karitiana individuals (selected at K14)*
*o K17 8 Surui individuals (selected at K18)*
*o K20 7 Kichwa Orellana individuals (selected at K20)*

We also used all African (73) and European (51) individuals (representing the respective ancestries). The Finns were excluded due to their known admixture with a central Asian population (Saag et al., 2019; Sikora et al., 2019). The entire dataset was screened for the LA of these selected individuals allowing us to identify 210 individuals showing less than 5% of non-NA ancestry (plus the 58 used as NA sources).

Merging all the positive results of these three independent analyses, we identified a restricted dataset with 217 almost unadmixed individuals (uIA217, Tables S2 and S3). Moreover, more stringent criteria, i.e. <1% African, <2% European (Gnecchi-Ruscone et al., 2019) and Z>|2|, were used to select a second restricted dataset with only 89 almost unadmixed individuals (uIA89, Tables S2 and S3).

##### Indigenous American dataset with masked haplotypes (mIA417)

The non-Indigenous component (>5%) identified using (LA) was removed from each haplotype of the 417 individuals, not included in the almost unadmixed Indigenous dataset. This masked dataset mIA417 combined with uIA217 (or uIA89) creates an overall Indigenous dataset encompassing 634 (506) individuals.

#### Uniparental analyses

##### Ancient mitogenomes

Processed reads from shotgun (single-end) sequencing were aligned to the rCRS sequence (Andrews et al., 1999) with BWA v0.7.17 aln/samse algorithm (Li et al., 2009) and realigned with CircularMapper (Peltzer et al., 2016). Duplicate reads were removed with Picard MarkDuplicates (https://github.com/broadinstitute/picard) and BAM files were further processed with SAMtools (Li et al., 2009).

Raw paired-end reads derived from captured mitogenomes that overlapped for at least 11 bases were merged using ClipAndMerge v1.7.7 (Peltzer et al., 2016) and then processed as above (see “Ancient data preparation for analyses” section) with an additional step to remove new indexes used for multiplexing on the MiSeq sequencer. Clean reads were mapped to rCRS (Andrews et al., 1999) with BWA v0.7.17 mem algorithm and BAM files were filtered with Picard MarkDuplicates (https://github.com/broadinstitute/picard) to remove duplicates and with SAMtools (Li et al., 2009). The final mtDNA BAM of the four captured individuals were merged with the shotgun mtDNA BAM using SAMtools merge.

For all mitochondrial BAM files, only reads with minimum mapping and base quality of 30 and positions with a minimum depth of 1 were retained for downstream analyses. Eventually, we obtained samples with genome coverage ≥ 0.99 for a total of 11 individuals.

Two strategies were used to determine mitochondrial haplotypes for these 11 individuals. First, we performed variant calling with BCFtools (Li et al., 2009) and filtering the VCF files with VCFtools (Danecek et al., 2011). Haplotypes were refined by manually checking BAM files.

Then, to better define indels in our dataset, as there are diagnostic deletions for some Indigenous lineages, we realigned cleaned reads of our ancient individuals to modern Panamanian mitogenomes belonging to the same haplogroup. The alignment was performed with BWA 0.7.17 aln/samse algorithm (Li et al., 2009) and reads were realigned with CircularMapper (Peltzer et al., 2016). Consensus sequences were generated using the same filters as before and then compared to the rCRS to obtain final haplotypes.

We also reconstructed the consensus sequence for the four contaminated individuals. We used ANGSD (Korneliussen et al., 2014) applying the same filters as in (Sánchez-Quinto et al., 2019). Haplogroups classification, based on phylotree.org (mtDNA tree Build 17) (Van Oven, 2015), was assessed using the online tool HaploGrep2 (Weissensteiner et al., 2016) (Table S1).

##### Modern mitogenomes

Raw paired-end reads were processed as follows: read adaptors were trimmed and reads were aligned to rCRS (Andrews et al., 1999) with BWA mem algorithm (Li et al., 2009). BAM files were filtered with Picard MarkDuplicates (https://github.com/broadinstitute/picard) and SAMtools (Li et al., 2009). Variant calling was performed with GATK HaplotypeCaller (McKenna et al., 2010) and mitochondrial haplotypes were also checked by manually inspecting BAM files. HaploGrep2 was used for haplogroup assignment (Table S2). Four mitogenomes (PaGUN9659, PaGUN9671, PaNAS16050, and PaNGA1193) were obtained with Sanger sequencing and analyzed using Sequencher v5.0 (http://www.genecodes.com/).

##### MtDNA tree, dating and demography

Phylogenetic tree and Bayesian Skyline Plot (BSP) were generated using BEAST v2.6.2 (Bouckaert et al., 2019). BEAST was also employed to calculate Bayesian age estimates. Radiocarbon dates of ancient individuals were used as priors. The L2c2 mitogenome from a Moreno individual (PaMOR16007) was included in the analyses as an outgroup. BEAST runs were performed with complete mtDNA sequences under the HKY substitution model (gamma-distributed rates plus invariant sites) with a fixed molecular clock as in (Brandini et al., 2018). We set the clock rate considering the ones published in (Posth et al., 2016; Soares et al., 2009). The chain length was established at 10,000,000 iterations with samples drawn every 1,000 Markov chain Monte Carlo (MCMC) steps, after a discarded burn-in of 10% steps (default value 0). Panama-specific haplogroups were set as monophyletic in the analyses. The same BEAST settings were used to: i) estimate the ages of haplogroups and (ii) evaluate population expansions in Panama through BSPs (Drummond et al., 2005) by including all Panamanian Indigenous mitogenomes analyzed in this study. BSPs were visualized in a plot using Tracer v1.7 (Rambaut et al., 2018) and converted into an excel graph by assuming a generation time of 25 years as in (Brandini et al., 2018). The maximum clade credibility tree was determined using TreeAnnotator and visualized with FigTree (http://tree.bio.ed.ac.uk/software/figtree/).

##### Ancient Y chromosomes

The Y-chromosome haplogroup classification of the nine ancient males, previously identified (see Ancient data preparation for analysis section), was deducted from the aDNA aligned sequences by: i) extracting with bcftools all the positions belonging to the Y chromosome; ii) considering only the positions that matched the list of the SNPs belonging to the main branches of the phylogenetic tree present in (Poznik, 2016) after taking into account any possible aDNA damage (C/T --> T/C; G/A -->A/G) as in (Fregel et al., 2018). To further sub-classify the ancient Y chromosomes the same workflow was performed by considering the list of 1,104 specific haplogroup Q SNPs reported by (Grugni et al., 2019). All codes and pipeline for this part can be found at the link: https://github.com/raveancic/aDNAYchromosome.

##### Modern Y chromosomes

The Y-chromosome haplogroup classification of the 43 modern male individuals was first inferred from genotyped files by using the script called Haplogroups.py in Yhaplo with Python3 (Poznik et al., 2016) (https://github.com/23andMe/yhaplo), using default parameters. Then, the obtained classification was confirmed by hierarchical analysis, as previously described (Battaglia et al., 2013), of the following Y-chromosome haplogroup markers: M9, M242, M3, M89, YAP, M96, M304, M172, M241, M269, L23, S116. In addition, the M242 positive samples (Hg Q) were further sub-classified by typing the signature markers (M848, Z780, M925, Z5908, Y780, CTS2731) of the main Indigenous sub-haplogroups recently identified (Grugni et al., 2019; Pinotti et al., 2019). Haplogroup nomenclature is according to (Grugni et al., 2019).

#### Population genetics analysis based on allele frequencies

##### Principal Component Analysis (PCA)

PCAs were performed using ‘smartpca’ program from the package EIGENSOFT v7.2.0 (Patterson et al., 2006). Ancient data, characterized by a large amount of missing data, were projected onto the modern variation with the lsqproject and autoshrink options. The same approach was used for the masked dataset (*mIA417)* that also shows a variable amount of missing data. Several PCAs were performed considering ancient and modern world-wide datasets and different sub-datasets. Those individuals showing peculiar outlier positions in the PCA plots were excluded from the downstream analyses (Tables S1-4).

##### ADMIXTURE clustering analysis

Different datasets (only modern and modern plus ancient individuals) were pruned with using PLINK 1.9 (-- indep-pairwise 200 25 0.4) and used to perform a biogeographical ancestry analysis with ADMIXTURE v.1.23 (Alexander et al., 2009). We performed ten independent unsupervised ADMIXTURE runs for each K, from K1 to K20, adding the –cv flag to identify the 5-fold cross-validation (CV) error for each K. The average cross-validation (cv) value for each K were plotted to select the most likelihood model (Figures S10 and S12). The software CLUMPAK (Kopelman et al., 2015) was used to combine different runs and to find the best alignment of the results across a range of K values with the tool DISTRUCT (Rosenberg, 2004). An additional ADMIXTURE analyses was performed projecting the ancient individuals on to the population structure from only modern individuals using the option *-P* and the *.P file* from K14 (Figure S11).

##### Admixture tests (f-statistics)

The *f* statistics was performed using EIGENSOFT v7.2 and AdmixTools v4.1 (Patterson et al., 2012). Outgroup *f3* was used to highlight only the shared genetic history between individuals or populations relative to an outgroup (Peter, 2016). High value of *f3* means more genetic history shared between the pair population analyzed. This method is less sensitive to lineage-specific genetic drift over the use of pairwise distance measures, such as Fst (Skoglund et al., 2015). A graphical explanation of the outgroup *f3* is reported in Figure S18.

We analyzed the shared genetic history of modern IA populations (included in the mIA417 and uIA89 datasets) against some ancient reference genomes (aRG) from Siberia, Beringia, North America (representative of the NNA ancestry) and South America (representative of SNA). The Guna always show a shared genetic drift with the ancient reference individuals lower than the average *f3* value (dotted line) of all modern IA groups. It could be also noticed that the average *f3* value is higher in the comparison with Spirit Cave than with Anzick-1.

We also built a distance matrix using the inverse values derived from the outgroup *f3* statistics on all Central and South American populations pairs plus Anzick-1, Early San Nicolas (ESN), Spirit Cave and USR (as an outgroup). We retained only populations with more than 30K overlapping SNPs and significant Z-scores (>3.3) in all comparisons. This distance matrix was used to generate a neighbor joining tree (Figure S19) with the program PHYLIP 3.6 (Nägele et al., 2020). The tree was visualized with FigTree 1.4.4 (https://github.com/rambaut/figtree).

The *f4* statistics was eventually used to identify gene flows among different populations. The comparison was performed in the form *f4(W, X; Y/test, Outgroup),* as reported in the software documentation. A graphical explanation of the *f4* statistics analyses is reported in Figure S16. In each figure showing a *f4* statistics test the form is reported above the plot(s). Results from *f4* analyses presented in Figures 2C, S24-S28 display only tests in which the initial conformation of the tree is rejected (*p∼0.001*, Z-score >3.3 or <-3.3) (Moreno-Mayar et al., 2018), meaning that the investigated population (Y) has a significantly higher genetic affinity with W rather than with X if the *f4* results are positive, the opposite when values are negative. If the results (triangle spaces) are not displayed, the proposed tree cannot be rejected and there is no significant preferential relationships between the test population (Y) and W or X. In Figures S16 and S29 the Z-score is displayed in abscissa and the region where the tested tree cannot be rejected is highlighted in grey (see above).

In all *f4* statistics, we considered a minimum threshold of 30K SNPs, the comparisons with less SNPs are highlighted with a specific symbol (X). Due to the low number of SNPs retained in multiple analyses, the following individuals were excluded: Baja_100, CuevadelPerico_2700, Enoque_3500, Kaillachuro_4000, LosIndios_600, Moraes_5800, SanFranciscoBay_25, ShukaKaa_10300, SoroMikayaPatjxa_6800 and Tibes_1200.

In particular, to specifically check for the relationships between Anzick-1 and Spirit Cave, with the Isthmian populations as well as with other ancient and modern IA individuals from Central and South America, we run two *f4* statistics in the following forms *(Anzick-1, Spirit Cave; Isthmo, Mbuti)* and *(Anzick-1, Spirit Cave; Central and South IA, Mbuti)*. The datasets uIA89, mIA417 and ancient individuals were used considering different sets of variants: all SNPs, all SNPs without C to T and G to A variants, only transversions and only transversion with ancient individuals coverage downsampled to a maximum of 1X (to avoid coverage biases).

We observed a clear pattern that becomes significant when increasing the number of SNPs. We further verified this pattern using a *f4* statistics in the form *(USR, Anzick-1/Spirit Cave; Central and South IA, Mbuti)* that confirmed a higher proximity to Spirit Cave in comparison to Anzick-1. The same pattern has been also observed in the outgroup *f3* analyses (see above).

##### TreeMix

In order to obtain a maximum likelihood tree, we run TreeMix (Pickrell and Pritchard, 2012) on the pruned dataset uIA89 using TSI, CHB and YRI (Tuscans, Chinese Han and Yoruba) as outgroups. The -noss and -global parameters were added considering zero to five admixture edges. The trees with the highest likelihood were selected after 1,000 runs (Gnecchi-Ruscone et al., 2019; Moreno-Mayar et al., 2018).

##### Ancestry modelling with qpWave

In order to verify that the Isthmo-Colombian ancestry (UpopI) is independent from other IA ancestries, we compared in pairs the Isthmian populations with all other modern and ancient IA populations using *qpWave* (Patterson et al., 2012) to test whether they were homogeneously related to a set of external outgroups. The outgroups were kept to the minimum and chosen to represent different IA ancestries identified here and in other papers:

- Mbuti, Papuan, CHB, Malta_24000 and USR_11500 as non-IA sources (Nägele et al., 2020; Posth et al., 2018)
- ASO_4000 and Chipewyan for NNA
- LagoaSanta_10400 for SNA1
- SpiritCave_10900 for SNA2
- Mixe for UpopA
- GuayaboBlanco_1700 for Archaic Caribbean
- Ayayema_4500 for Patagonia
- Aymara and KCH (K1 and K19 respectively in FigureS10) representing modern South American populations.

We took in consideration the *p-value* of “taildiff” for Rank1, a statistically significant *p-value* (< 0.01) means that each compared pair could be explained by two sources. We observed that one ancestry is usually needed (*p-value* >1) to define pairs of Isthmo-Colombian populations, while pairs of Isthmian and non-Isthmian populations require two ancestries. This pattern is more evident in the Guna, the best representative of the Isthmo-Colombian component. This pattern confirms that a different ancestry, instead of only genetic drift by isolation, is needed to explain the distinctiveness of the Isthmo-Colombian populations.

##### Demographic modelling with qpGraph

We used qpGraph (Patterson et al., 2012), on a merged dataset of the uIA89, mIA417 and ancient individuals (considering only transversions), to reconstruct the best tree modelling the relationships between Isthmian populations and ancient Indigenous genomes. In our *f4*-statistics we noted a differential relationship between Isthmian groups and modern/ancient Indigenous Americans in comparison to the individuals older than 10 kya (Figures 6A, S28). Therefore, we modelled a basal tree with three of the most ancient available genomes of the SNA ancestry, an ancestry that certainly went through the Isthmus to reach South America. Our best tree revealed that the SNA dispersal involved a complex demographic pattern, with three possible ancestries (Figure S32A). To resolve the inferred length-zero internal branch, we tested all three possible split orders obtaining similar scores. Therefore, it might represent a very short branch that we cannot resolve with this dataset power (Lipson, 2020) and the three lineages, Anzick-1 (Montana, ∼12.6 kya), Lagoa Santa (Brazil, ∼10.4 kya; SNA1) and Spirit Cave (Nevada, ∼10.9 kya; SNA2) are statistically consistent with forming a trifurcation. To increase the resolution power and considering the results of previous analyses (*f3-* and *f4*-statistics), we added to this graph the captured and genotyped data from Lapa do Santos (Brazil, ∼9.6 kya) as SNA1 and the whole-genome sequences of ESN (California; ∼4-5 kya) as SNA2. This allowed us also to check for any bias due to sequencing methods. Even in this case the best fit tree confirms the trifurcation (Figure S32B). After this step, we added Los Rieles (Chile, ∼10.9 kya), the most ancient Pacific coast genome, which turned out to be better modelled as an admixture of SNA1 and SNA2 (|Z|=2.835) than as non-admixed and considering only geographic origins (|Z|=3.930) (Figure S32C). This finding confirms that both SNA1 and SNA2 reached South America and seems to indicate that the latter had a lower impact on the Atlantic side of South America. When Los Rieles is replaced by Lapa do Santos, the tree does not fit (|Z|=3.930) (Figure S32D).

We then took into account that, in the *f4*-statistics (Figures 6A, S26), there is a significant allele sharing of the Isthmian populations with Spirit Cave and two Central American ancient genomes, Mayahak Cab Pek (Belize, ∼9.3 kya) and Saki Tzul (Belize, ∼7.4 kya), as well as a higher genetic proximity to Los Rieles relative to Lagoa Santa (Figures S26, S27). Therefore, we attempted to model ancient Panama in relation to these ancient genomes (Figure S33). Ancient Panama fits better when considering an admixture between the Central-South American branch of SNA2 and another ancestry parallel to SNA2, and shows the best score with Mayahak Cab Pek (Figure S33B). The additions of NNA ancient individuals (Figure S34), ASO (Ancient Southwestern Ontario, ∼4.2 kya), 939 (Lucy Island, British Columbia, ∼6.1 kya) and Kennewick (Washington State, ∼8.8 kya), hold better when assuming some admixture events involving these genomes. These findings confirm that we cannot identify an unadmixed proxy for the NNA ancestry among ancient individuals. The best graph was obtained when including ASO as NNA and Mayahak Cab Pek as the central American ancient genome (Figure S34B).

The next step was to add the Guna modern group, which was the best representative of the Isthmian-specific component in our previous analyses (Figure 2). In this setting only the admixed model was supported, with the Guna group representing a still unsampled population of the Isthmus, UPopI (Figures S35, S36). To assess if UPopI might correspond to the previously identified UPopA, we replaced Guna with Mixe, previously used to identify UPopA (Moreno-Mayar et al., 2018). The resulting admixture graphs were not statistically supported (|Z|>3 in all attempts (Figures S35, S36).

At this point, we further evaluated the relationship of Panama with SNA1 that we initially linked to Lagoa Santa. As before, we started from basal admixture graphs without Guna (Figure S37A) and considered the conformation with ASO and Mayahak Cab Pek, based on previous results (Figure S34). We first tested the tree without admixture in Los Rieles, placing Lagoa Santa as a parallel branch of SNA2, but the tree would fit only when considering Los Rieles as an admixture between SNA1 and SNA2 (|Z|=2.896), the two early South American ancestries that we identified above (Figure S37A). It is worth mentioning, however, that the graph without admixture in Los Rieles became statistically significant (|Z|<3) when Lagoa Santa was replaced with the younger Laranjal sample (Brazil, ∼6.8 kya) (Figure S37B). This confirms our *f4*-statistics (Figure 6A) and the scenario of a widespread population turnover in South American during mid-Holocene, as previously suggested (Moreno-Mayar et al., 2018; Posth et al., 2018), a finding that also correlates with climate changes in the southern continent (Riris and Arroyo-Kalin, 2019). As for the Ancient Panamanians, they are a mix between the source of SNA2 (prior to Spirit Cave) and the admixture between SNA1 and SNA2 (Figure S37A).

Lastly, we assessed different admixture graphs including also the Guna. We obtained the best Z-score in the graph when UPopI was placed as a parallel ancestry to SNA1 and SNA2, all radiating from the same early SNA source (Figure S37C). The latter derives from an initial split of the early Indigenous group into SNA and NNA. In this scenario the ASO group is the result of an admixture between NNA and SNA1. It is therefore likely that the first NA split occurred further north and earlier than the diversification of SNA into SNA1 e SNA2. Finally, the same graph shows that when UPopI reached the Isthmian area admixed locally with population groups derived from both SNA1 and SNA2.

Taking into account the presence of a zero-length branch in the final tree with UPopI (Figure S37C) and the results obtained on the basal graph (Figure S32A), we modeled a tree without zero-branches when considering Anzick-1 and Spirit Cave admixed with the same ancestry that leads to Lagoa Santa (Figure 38A). Therefore, we rebuilt the final graphs (with and without NNA) by considering the early SNA2 source admixed with SNA1 (Figure S38B). This admixture gave rise to two ancestries, one, related to Spirit Cave, that reached South America, leaving evident footprints on the Pacific coast, and another restricted to the Isthmo-Colombian area (UPopI) that is well represented in Guna. The best topologies were also checked replacing Guna (UPopI) with Mixe (UPopA) and Spirit Cave with Anzick-1 (Figures 38B, 38C), but no statistically supported graphs were found.

#### Population genetics analysis based on reconstructed haplotypes

##### Phasing

Phased haplotypes were generated from the rWD1560 dataset using the Segmented Haplotype Estimation and Imputation tool SHAPEITv2 (Delaneau et al., 2011) and the HapMap37 human genome build 37 recombination map.

##### Local Ancestry and Masking

The local ancestry for genomic fragments in the American individuals was estimated using RFMix (Maples et al., 2013). As source populations, we used Bantu, Esan (ESN), Gambia (GWDwg), Mandenka, Mbuti and Yoruba (YRI) for Africa, Spanish (IBS), British (GBR), French, Icelandic and Tuscany (TSI) for Europe and Chipewyan, Kichwa Orellana, PaGUNA, Puno, Surui and Karitiana for Indigenous ancestry. We used “PopPhased”, “-n 5” and “--forward-backward” options as recommended in RFMix manual. Then, starting from RFMix output files, we built a PLINK file set in which the non-Indigenous SNPs were masked. The masking process was done with this rationale: if in the “Viterbi” output a particular SNP was not assigned to the Indigenous ancestry and if the probability of belonging to the Indigenous ancestry (reported in the “forwardbackward” output) was less than a threshold (< 0.9) that allele was set as missing. In this analysis we kept individuals as separated into the two phased haplotypes.

##### ChromoPainter

To obtain the painting profile of all the 217 individuals in the uIA217 dataset consisting in a matrix of ‘recipient’ individuals (rows) that appear as a mosaic of the ‘donors’ (columns), we processed the genomic information contained in phased data (haplotypes) through the use of inferential algorithms implemented in CHROMOPAINTERv2 (Lawson et al., 2012). Technically for this analysis the recombination (-n) and mutation (-m) parameters used were respectively 233.1352 and 0.00084 estimated on five randomly selected chromosomes (3, 7, 10, 18 and 22). Since the genetic variability among NA populations is low, we run CHROMOPAINTER in two runs, one with standard parameters, the other adding the flag -k 50 (Gnecchi-Ruscone et al., 2019). No significant differences were observed between the two runs.

##### fineSTRUCTURE

The CHROMOPAINTER square (217 x 217 individuals) *chunkcounts.out* matrix was used as input file for fineSTRUCTURE in order to identify similar genetic clusters. We ran the software with three millions MCMC iterations thinned every 10,000 and preceded by one million burn in iterations: -x 1000000 ; -y 3000000 ; -z 10000 ; -t 1000000. The MCMC file (.xml) was used to build the tree structure using both the options –T1 and –T3, without major changes between the two methods.

Initially, we obtained 50 clusters in the final tree (data not shown). However, to obtain more robust genetic inferences the number of clusters was reduced to 19 considering the number of individuals in each cluster (less than five) and the Total Variation Distance (TVD <0.03) as elimination criteria. TVD is an index that measures the similarity between copying vectors of the CHROMOPAINTER matrix (calculated on the chunklengths) (Leslie et al., 2015); lower values of TVD mean similarity, while higher values indicate heterogeneity.

##### Haplotype Analyses on ‘Masked’ Individuals

The masked haplotypes (mIA417) were initially filtered for the individuals that had a maximum of 50% of missing SNPs (considered as the mean of the summed missingness of the two haplotypes) and, among these individuals, we selected only those with at least 25% of SNPs retained in each haplotype. Eventually, we obtained a restricted dataset of 311 masked individuals (rmIA311). This dataset (rmIA311) was then converted in PLINK1.9 format and subsequently in a CHROMOPAINTERv2 input. To enable missing data in CHROMOPAINTERv2, we slightly modified CHROMOPAINTERv2 such that a recipient/target’s emission probability is set to 0 at missing (i.e. masked) SNPs when tabulating the expected number of segments matched to each “Donor” individual that the recipient is compared to. Therefore, in regions of high missingness the expected number of segments matched to each “Donor” will tend towards the prior, which assumes equal matching to all “Donor” individuals. However, in our application here we found that inference seems to be dominated by data at non-missing SNPs, where the usual CHROMOPAINTERv2 machinery is employed. In particular, we did not identify any correlation between the percentage of missing data (even when reaching 50%) and bad placements/outlier behaviours in the PCA created from the CHROMOPAINTER output, both projecting and not projecting the ‘masked’ individuals (Figure S14).

##### Identical by Descent (IBD) Analysis

The pattern of IBD sharing within each population of the uIA217 phased dataset was analyzed using Refined-IBD (Browning and Browning, 2013), which allows to improve the accuracy and efficiency of identity by descent detection in population data, using default parameters. The average IBD-sharing was calculated for nine different bin categories reported in (Gnecchi-Ruscone et al., 2019).

In order to reconstruct the population dynamics, we applied IBDne on the uIA217 dataset, using both IBDseq, which did not require phased data, and Refined-IBD that uses phased data, analysing different IBD segment length (2-4 centimorgan, cM). Lower confidence intervals were obtained when using windows of 2 cM (Figure S20). Moreover, the Ne obtained with Refined-IBD is more compatible with historical estimates on the Panamanian pre-colonial population size (see introduction). Therefore, Refined-IBD was applied using 2 cM windows for the subsequent analyses. We observed the same trend of the ancestry specific effective population size (asIBDne) on the 74 Panamanian individuals, masked for the NA component, following the pipeline presented by (Browning et al., 2018), as reported in (Ongaro et al., 2019) (Figure S20). To retain more information, we used the three macro-cluster from the fineSTRUCTURE tree: Guna (30 individuals), Emberá (18 individuals) and Western Panama (20 individuals) encompassing Bribri, Naso and Ngäbe (Figure S21A). The Guna show a peculiar trend that can be related to the demographic history of this population (Figure S21B). Removing the Guna from the comparison, it is clear that no one of the other clusters could describe the decrease in population size observed in Panama, due to a strong bottleneck in the Guna before the arrival of Europeans. The same approach was applied including all macro-clusters obtained with fineSTRUCTURE (Figure S22C). This analysis confirmed the same trend of the Panama plot.

##### Dating Admixture Events with IBD Sharing

In order to date the admixture events between our Panamanian groups and other IA populations included in the uIA217 dataset, we calculated the IBD sharing segments using RefineIBD as performed by (Liu et al., 2020). The IBD blocks were divided into three categories, based on their length (1-5 cM, 5-10 cM and over 10 cM), each roughly representing different time periods: 1,500-2,500 ya, 500-1,500 ya and <500 ya (Liu et al., 2020; Ralph and Coop, 2013). We have calculated the mean of summed IBD lengths shared between population pairs for each length category. To reduce noise and false positives only the pairs that shared at least two blocks >5 cM and four < 5 cM were considered.

#### Additional Notes

The text has been revised in order to minimize the use of “colonial language” and to avoid the connotation of people as mere samples or data. The adjective Indigenous has been preferred to Native American with only notable exceptions of Northern Native American (NNA) and Southern Native American (SNA) ancestries, which were used for consistency with previous papers.

Many figures in the paper have been created using various versions of the software Tableau (https://www.tableau.com/) or with different R packages.

---

**Figure S1.**
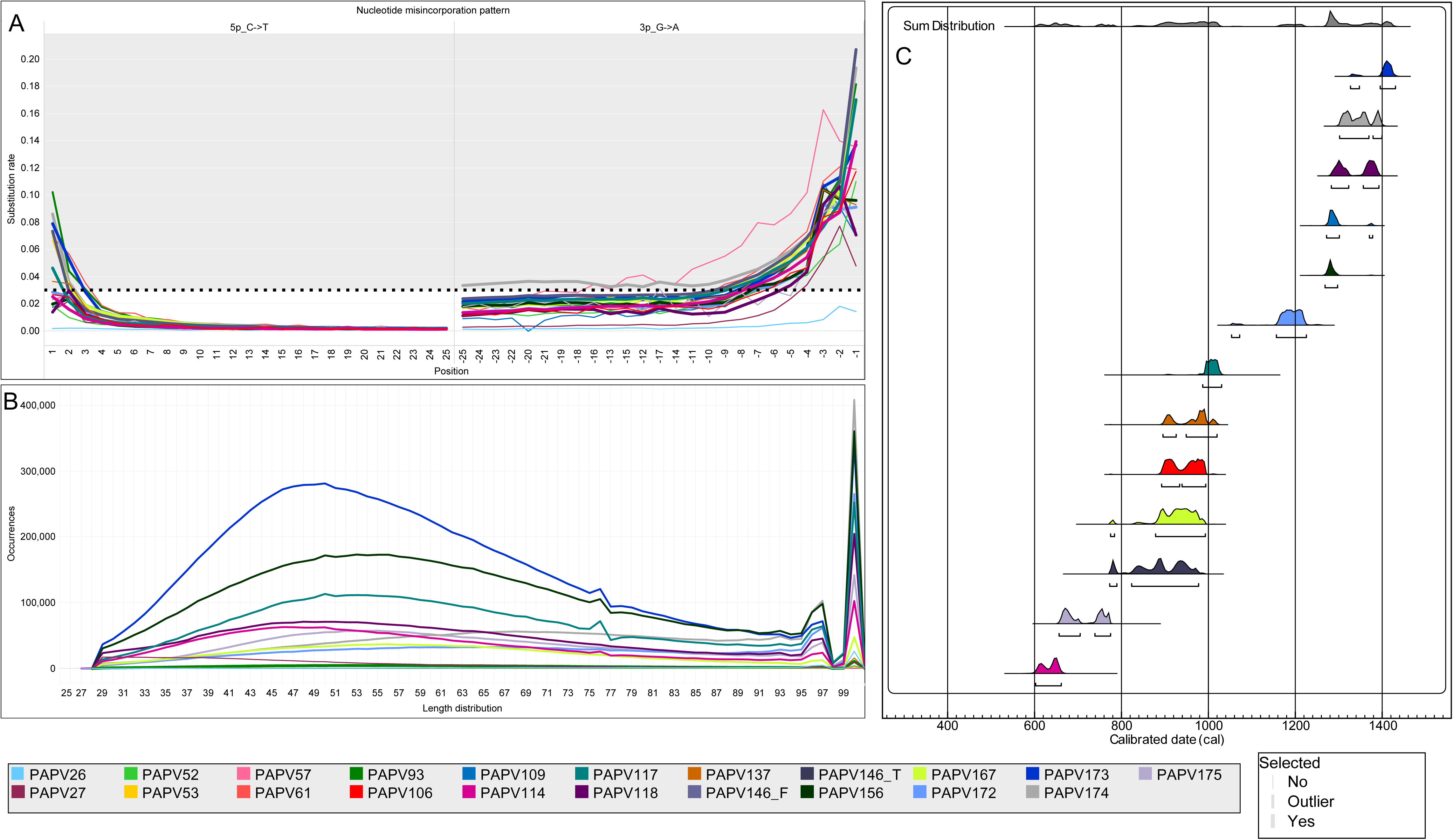
Post-Mortem Damage Patterns and Radiocarbon Dates. Related to Table S1. A) Rates of C to T and G to A substitutions. Thicker lines indicate data used in autosomal analyses (Table S1). B) Length distribution of reads. C) Radiocarbon dates.

**Figure S2.**
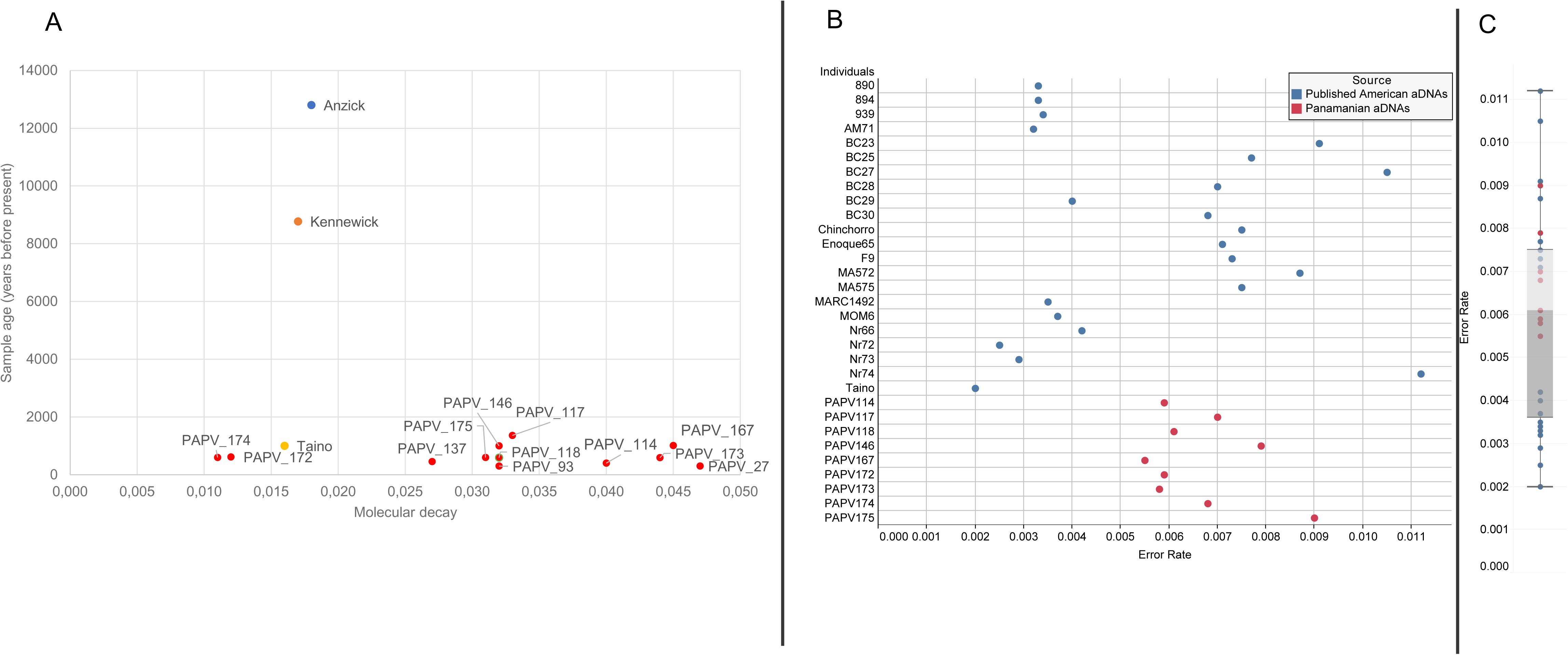
Molecular Decay and Error Rate. Related to Table S1. A) Molecular decay and age estimate of Panamanians and other three ancient Indigenous American genomes. B) Error rate comparison between Panamanians and other ancient Indigenous American individuals. C) Boxplot distribution of error rates, the light grey encloses the 75^th^ percentile while the dark grey the 25^th^ percentile.

**Figure S3.**
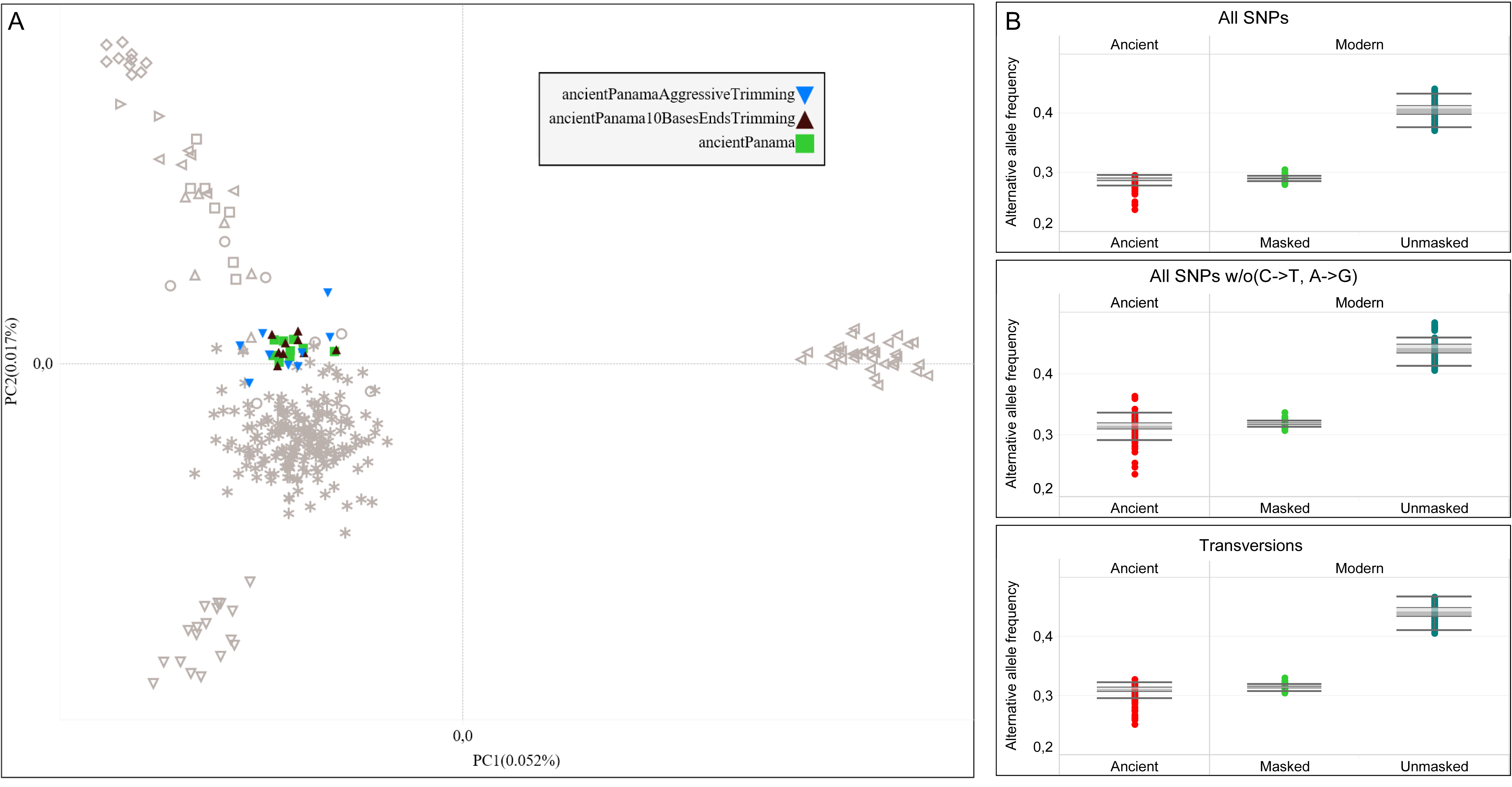
Additional Validation Tests on Ancient DNA Data. Related to Figure 2B. A) Isthmo-Colombian PCA analysis including three Ancient Panamanian datasets trimmed with different criteria: with original pipeline (green); trimming 10 bases from both the 5’ and 3’ ends of all reads (seal brown); removing adapters using a higher mismatch rate (up to 50%) (blue). B) Comparison of alternative allele frequency distributions between modern and ancient individuals considering different sets of variants: all SNPs, all SNPs without C to T and G to A and only transversions.

**Figure S4.**
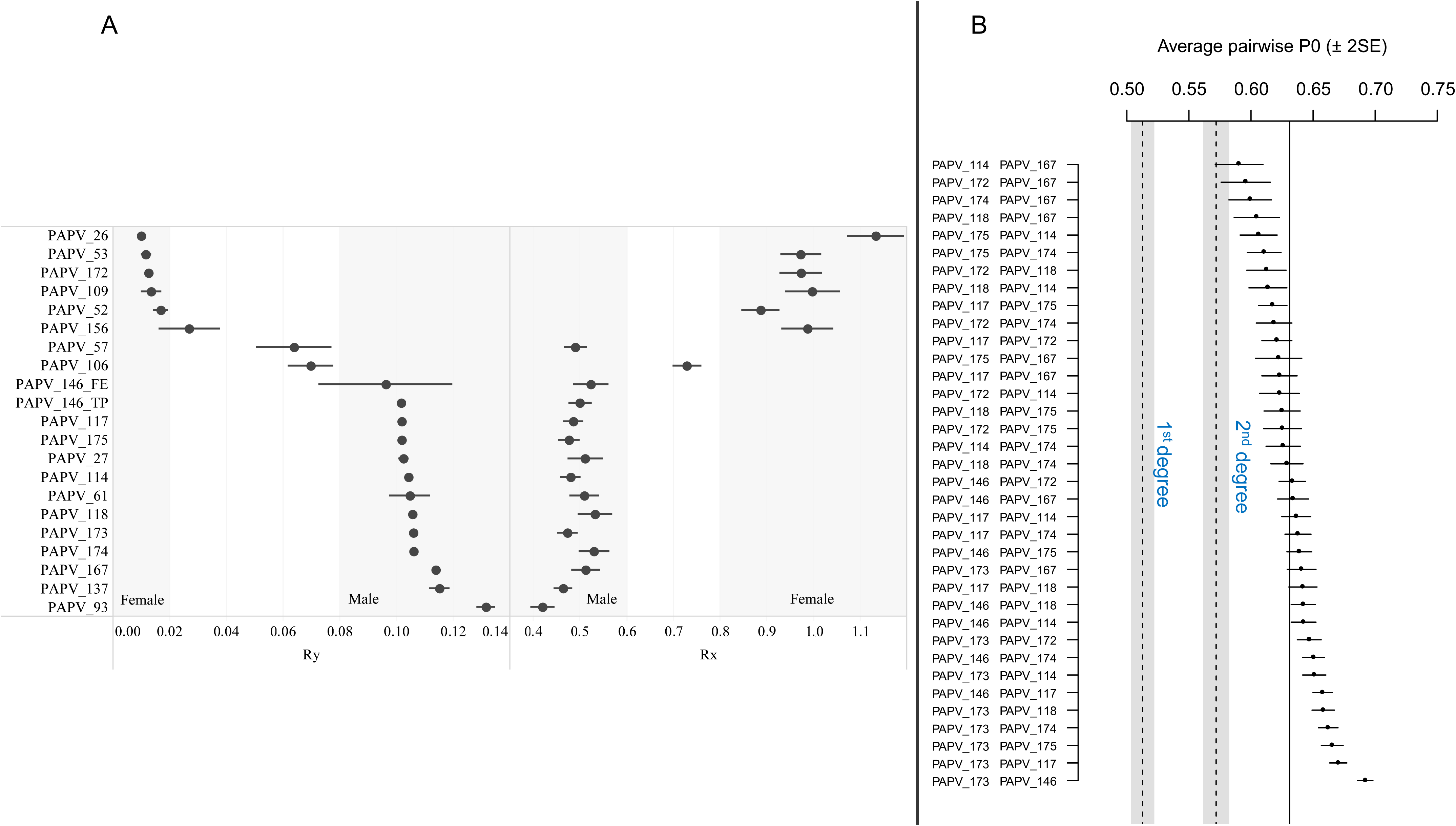
Molecular Sex Determination and Family Relationships. Related to Table S1. A) The sex of each individual was inferred using two computational approaches specific for low-coverage genomes, Rx (Mittnik et al., 2016) and Ry (Skoglund et al., 2013). B) Kinship analysis using the software READ (Relationship Estimation from Ancient DNA) (Monroy Kuhn et al., 2018).

**Figure S5.**
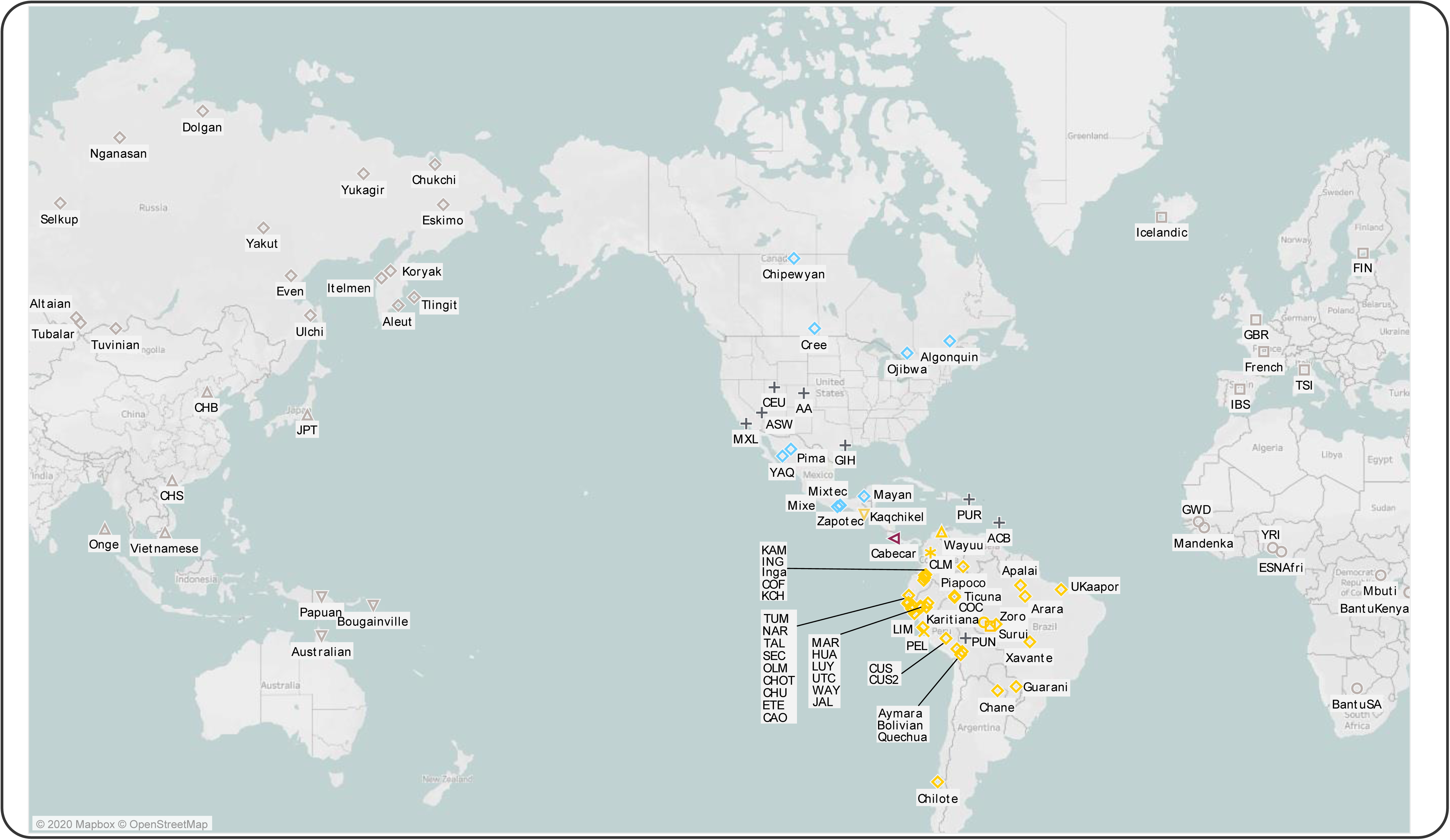
Geographic Origin of the Individuals in the rWD1560 Dataset. Related to STAR Methods. Symbols as in Figure 2.

**Figure S6.**
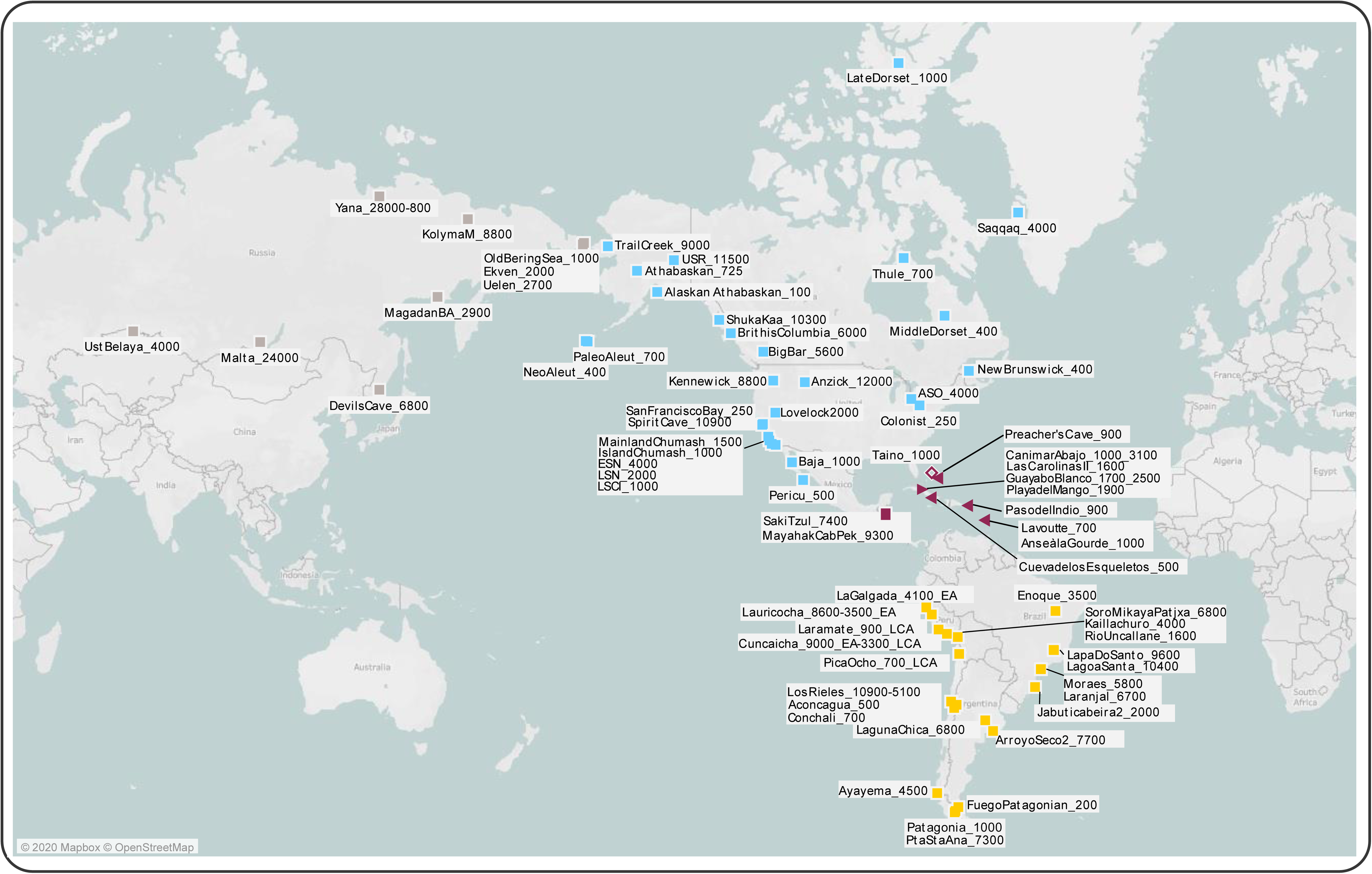
Geographic Origin of the Ancient American and Siberian Individuals Used for Comparison. Related to STAR Methods. Symbols as in Figure 2.

**Figure S7.**
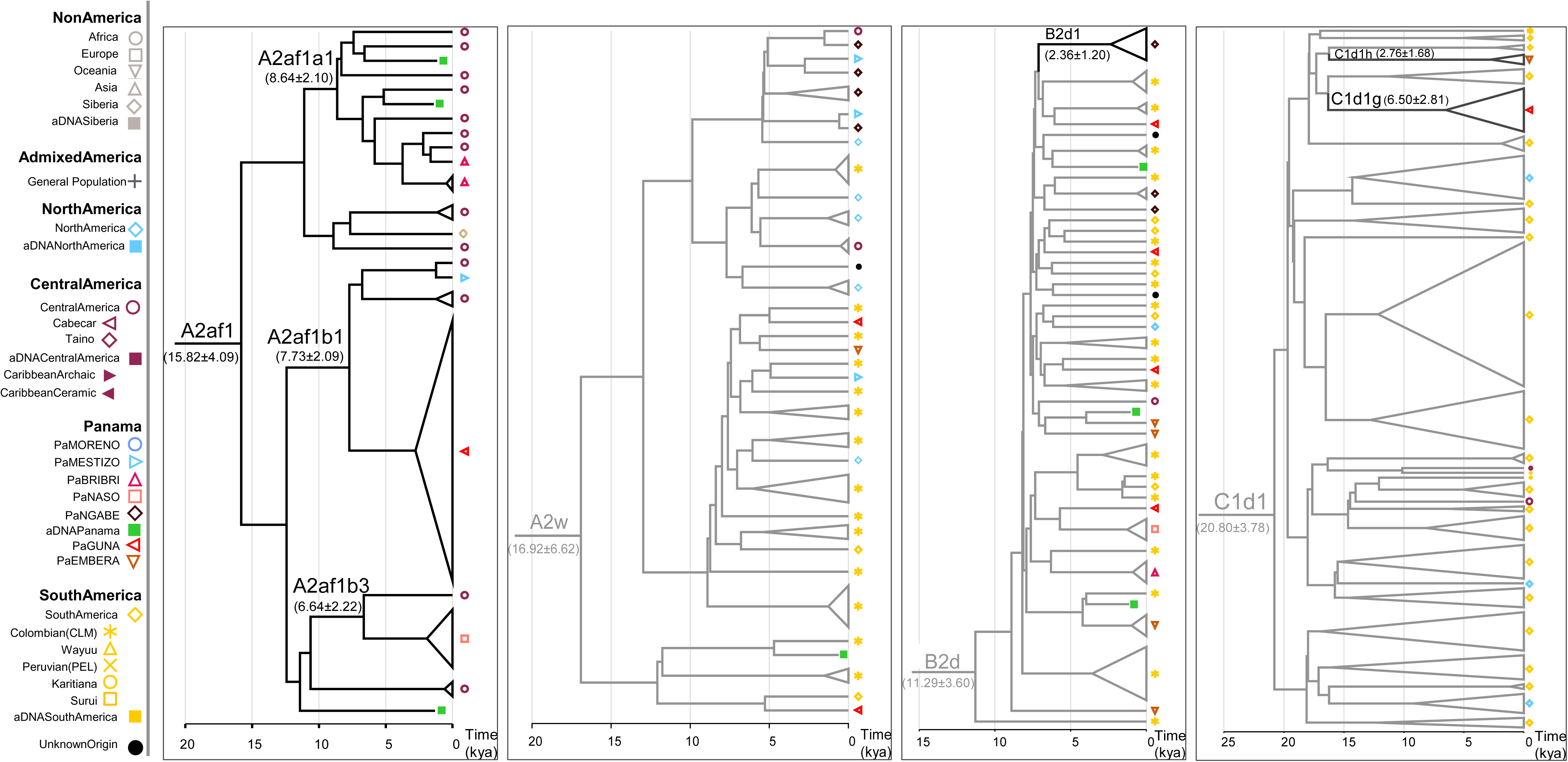
Schematic Phylogenetic Trees of the Four Major MtDNA Haplogroups Identified in the Isthmus. Related to Figure 5B. The Bayesian phylogenetic trees rooted on an L2c2 mitogenome from a Moreno individual include all modern mitogenomes belonging to the four major haplogroups of modern and ancient Panama (A2af1, A2w, B2d, C1d1). Black lines highlight branches specific to IA from the Isthmo-Colombian area. The Bayesian age (mean value with standard deviation) is shown for relevant branches.

**Figure S8.**
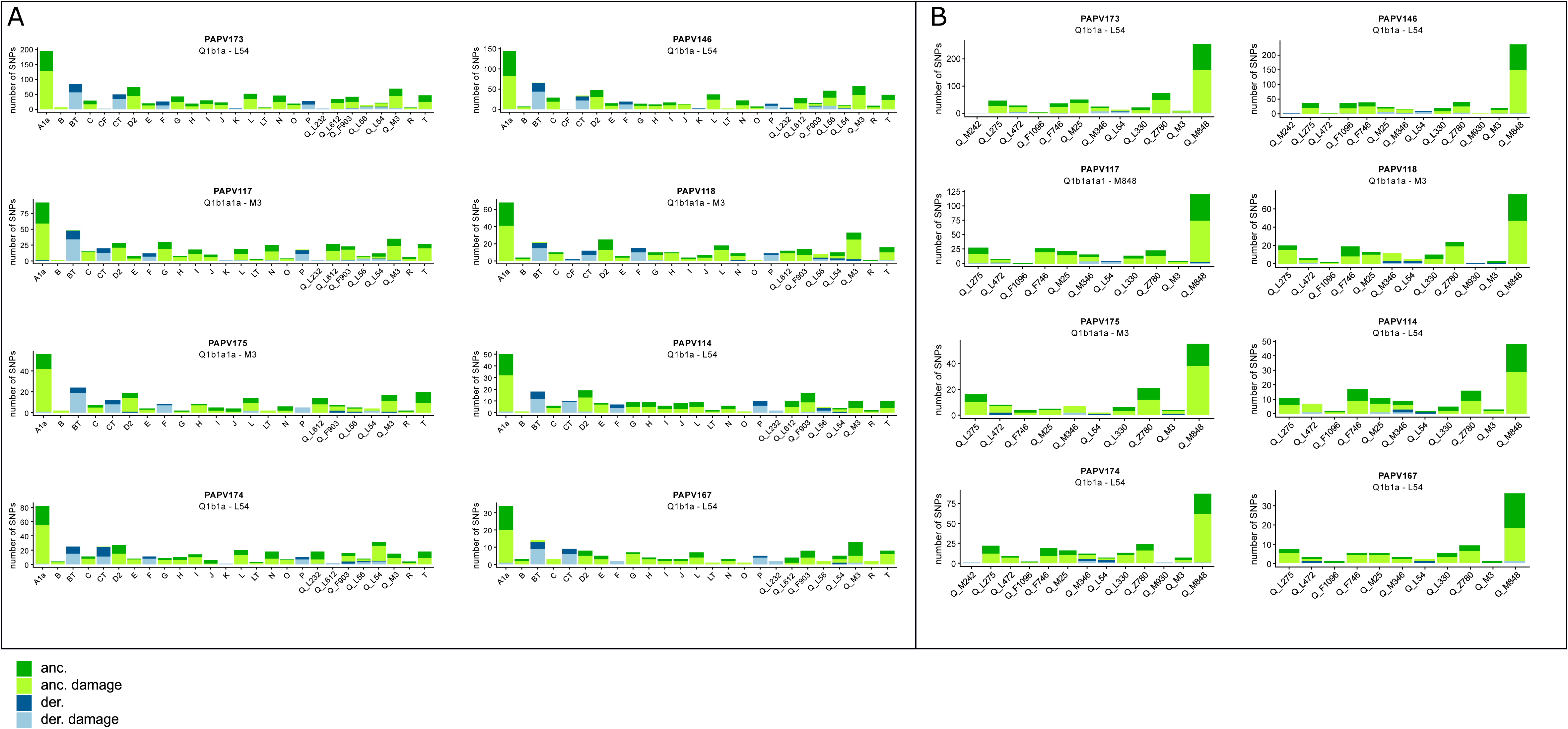
Ancient Y-Chromosome Classification. Related to Table S1. A) SNPs for each macro-haplogroup present in (Poznik et al., 2016). B) SNPs for each sub-haplogroup Q in (Grugni et al., 2019) and (Pinotti et al., 2019). Different colors refer to the allele status (green: ancestral; blue: derived), while different shades indicate the aDNA possible damage. Haplogroup nomenclature as in ISOGG 2019.

**Figure S9.**
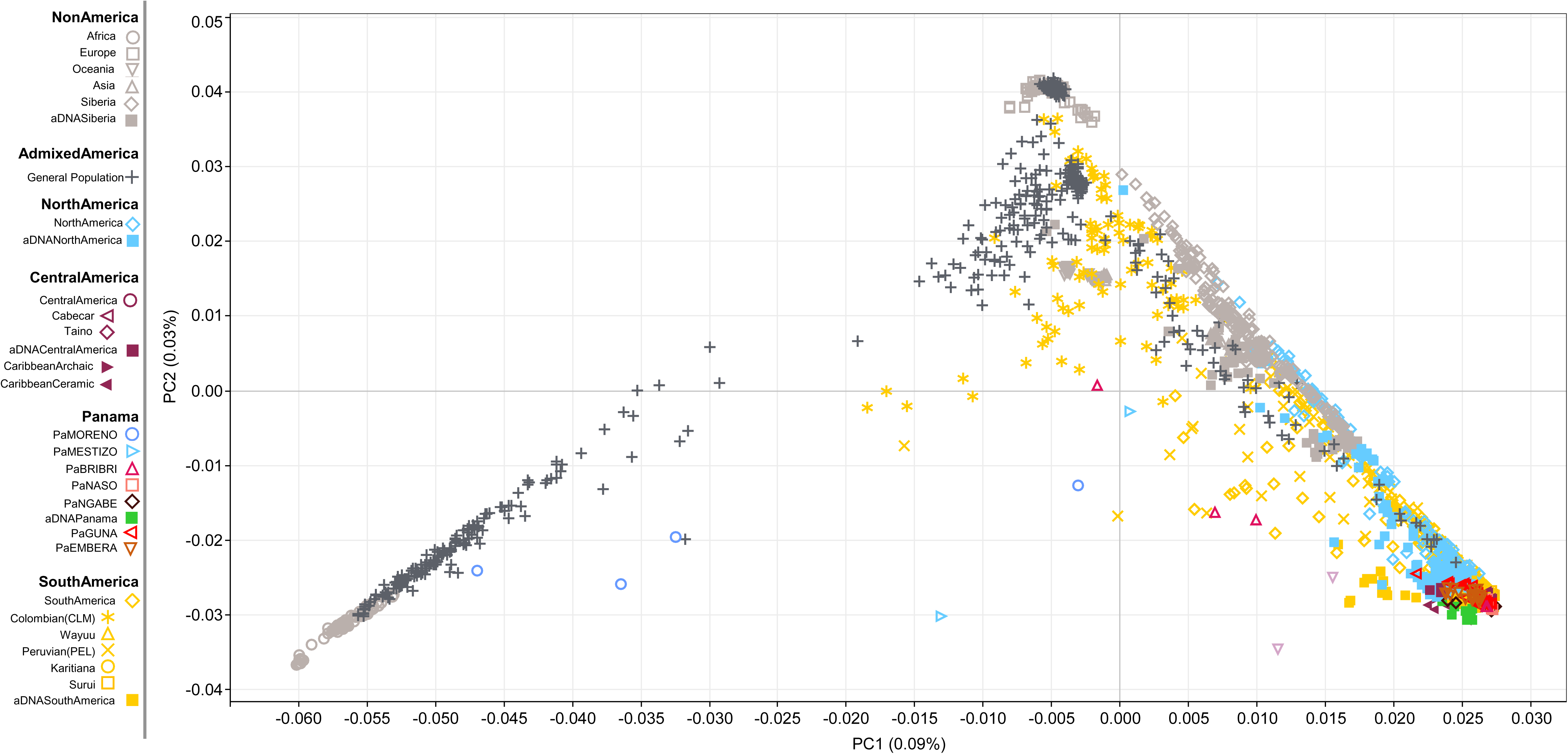
Worldwide PCA Plot on the rWD1560 Dataset. Related to Figure 2B. Ancient individuals from Siberia and the Americas were projected on PCs obtained from modern individuals.

**Figure S10.**
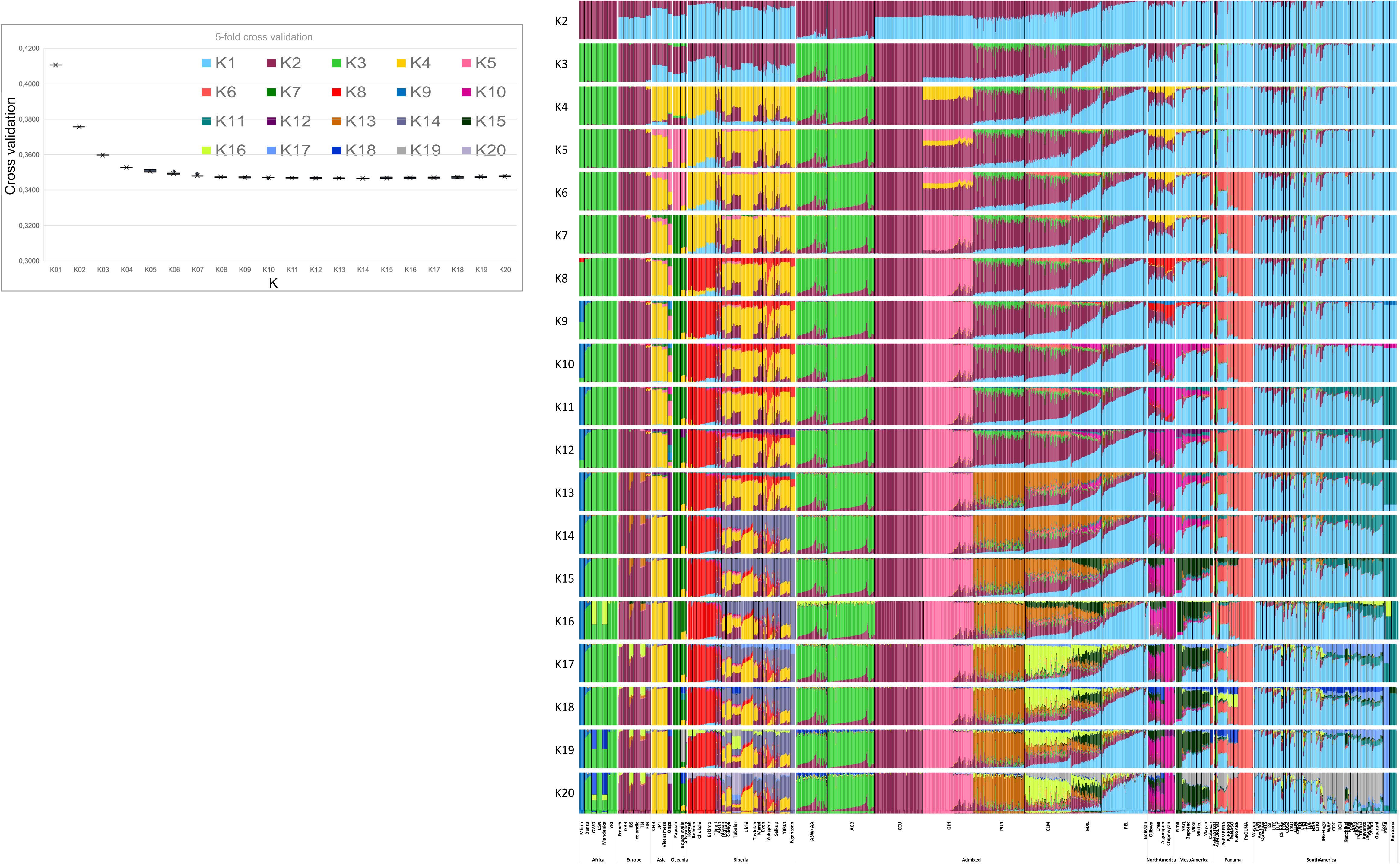
Worldwide ADMIXTURE Plot on Modern Individuals. Related to Figure 2A. ADMIXTURE analysis on the rWD1560 dataset from K2 to K20. The inset shows the boxplot of 5-fold cross validation values for each K after 10 runs. A highly structural variability, associated with low CV, is still present at K20 suggesting that the observed pattern is not an artifact. In the ADMIXTURE plot it is possible to observe seven specific IA clusters, some mirroring the PCA results (Figure 2B): • K1 that is widely distributed in all IA populations with high percentage in Puno, Aymara, Quechua and Paran-Cusco individuals, all speaking Andean languages. • K6 is modal in PaGUNA and highly represented in all the populations from Costa Rica to Panama, speaking Chibchan languages. • K10 is typical of the Chipewyan (speaking a Na-Dene language) with lower percentage in all populations from northern North America. • K11 is modal in Karitiana and Surui speaking Tupi (Equatorial-Tucanoan) languages. • K17 separates the Surui from Karitiana. • K15 is represented by Pima and mostly present in Mexico, but also widely distributed in Central and Northern IA speaking groups. • K19, likewise K1, is another component present in the entire double continent that reaches the highest level in South America, particularly in the Andes Mountains (i.e. KCH). Additional interesting components are K13 and K16 that are present in high percentage in Puerto Rico (PUR) and Colombia (CLM), respectively, but could also be observed in admixed American and European populations. The above-mentioned components are present in different proportions and differentially distributed among IA populations including the Panamanians, with the only notable exception of the Guna, showing their specific component.

**Figure S11.**
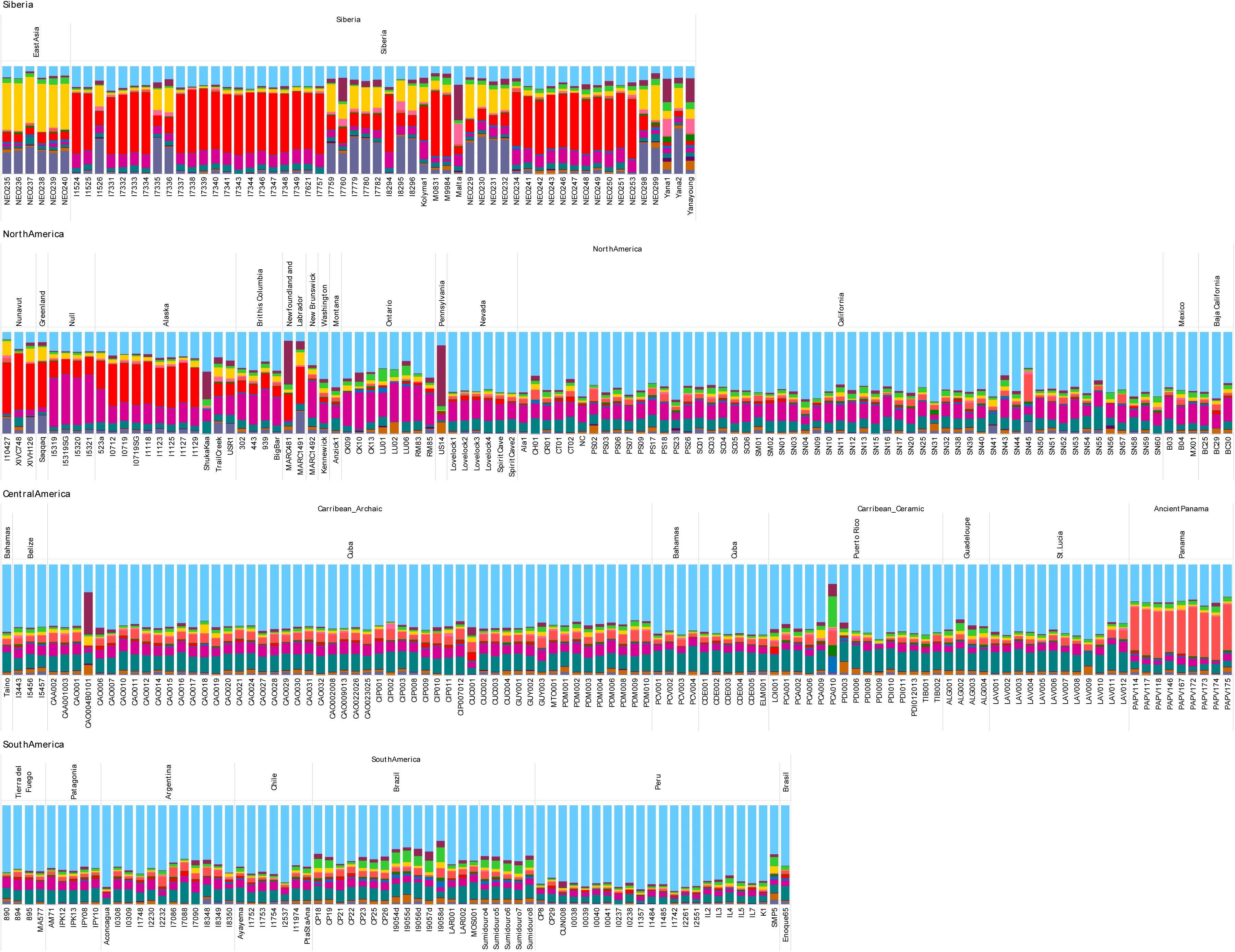
ADMIXTURE Plot Projecting Ancient Individuals. Related to Figure 2A. ADMIXTURE analysis projecting ancient Siberian and American individuals (Table S4) on the modern worldwide variability (Figure S10). The .P file of K14 was used.

**Figure S12.**
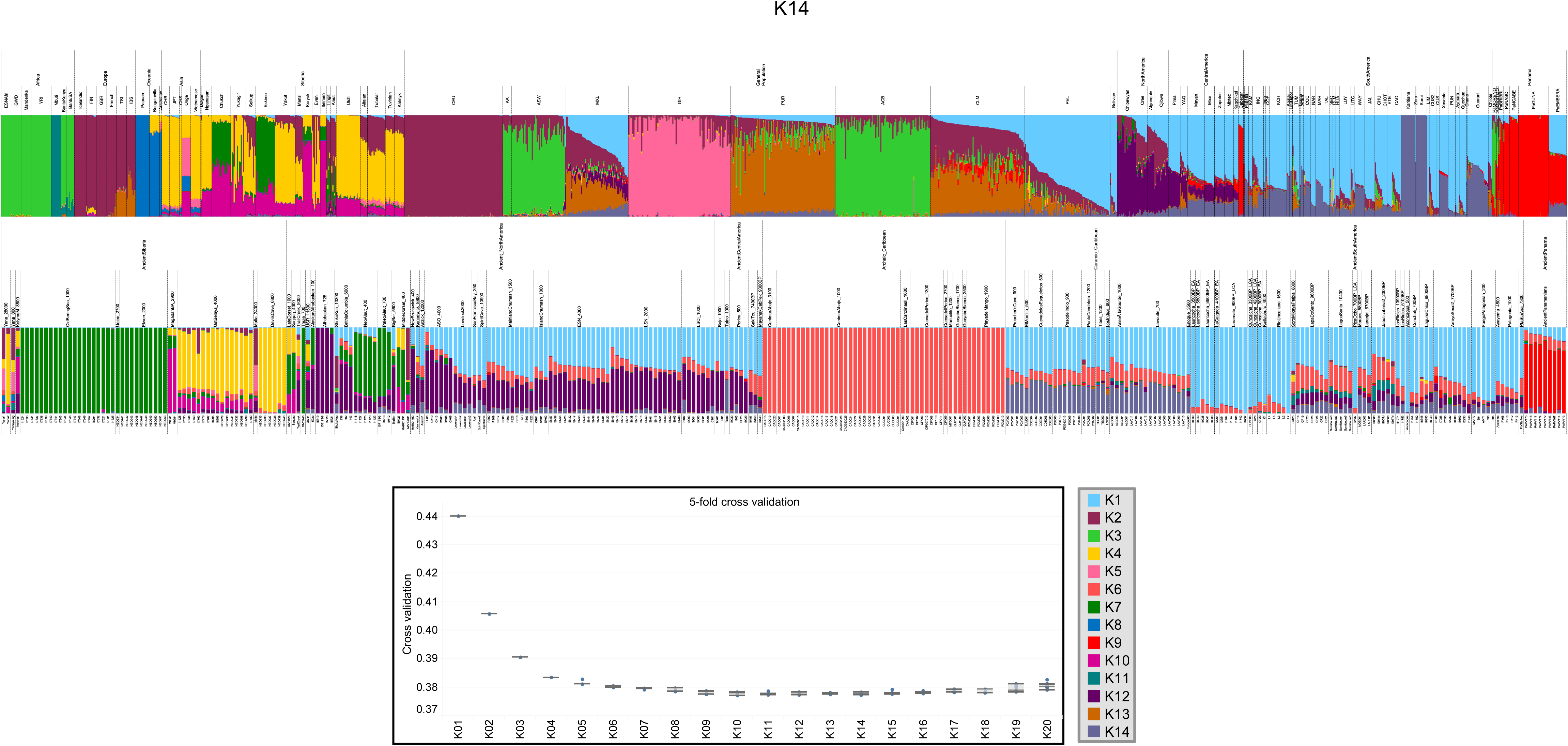
ADMIXTURE Plot including Ancient Individuals. Related to Figure 2A. ADMIXTURE analysis including the rWD1560 dataset plus the 329 ancient Siberian and American individuals. Only K14, which has the lowest error in the 5-fold cross validation boxplot (inset), is shown. This analysis confirms the results already discussed in Figure S10, adding a new K specific of Archaic Caribbean individuals .

**Figure S13.**
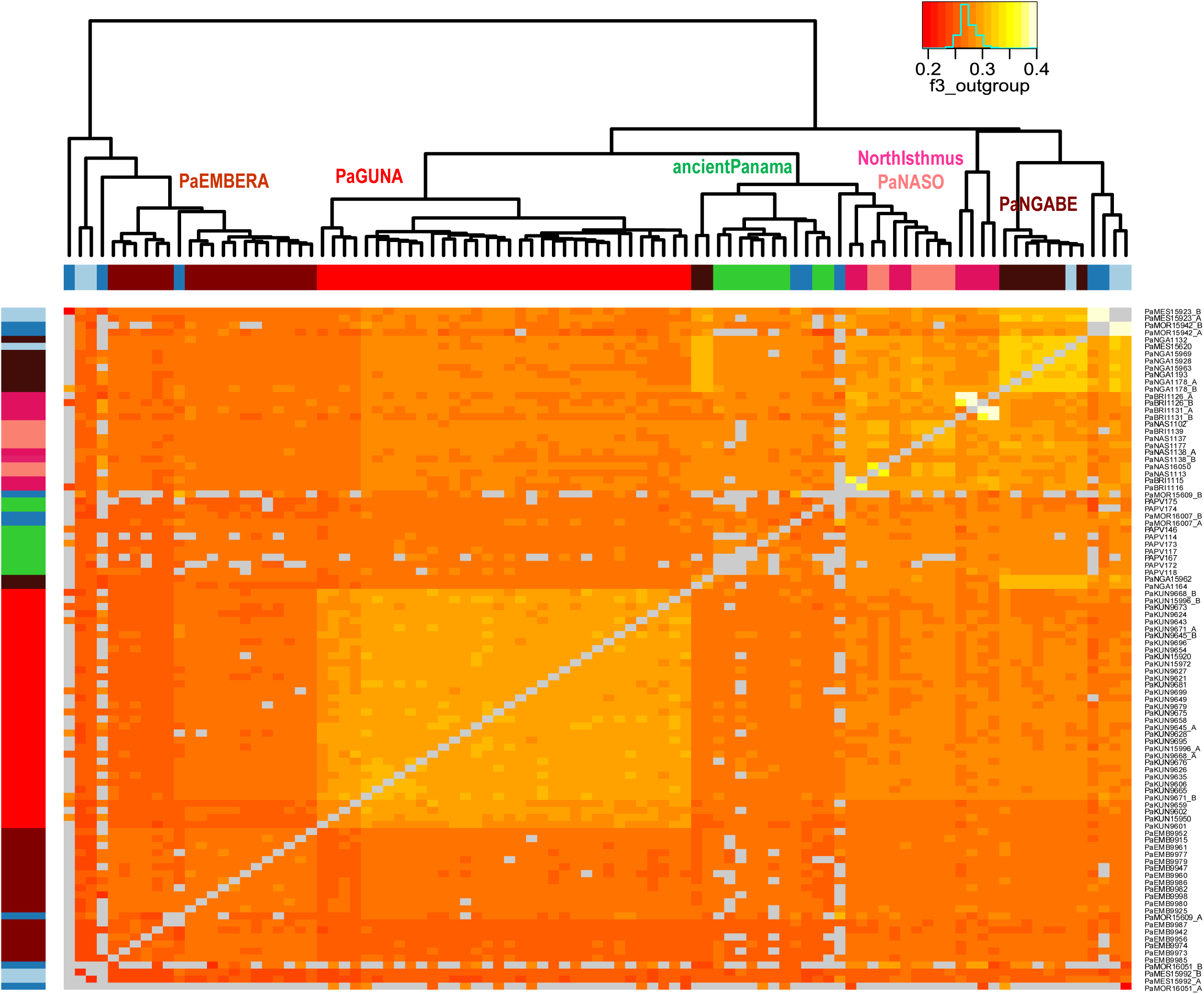
Heatmap Based on Outgroup *f3-statistics* of Panama. Related to Figure 2B. The shared drift among the Panamanian individuals was analyzed considering those included in uIA217 plus the masked data in mIA417. Color intensity is inversely proportional to the shared ancestry among individuals, which was used to build the dendrogram.

**Figure S14.**
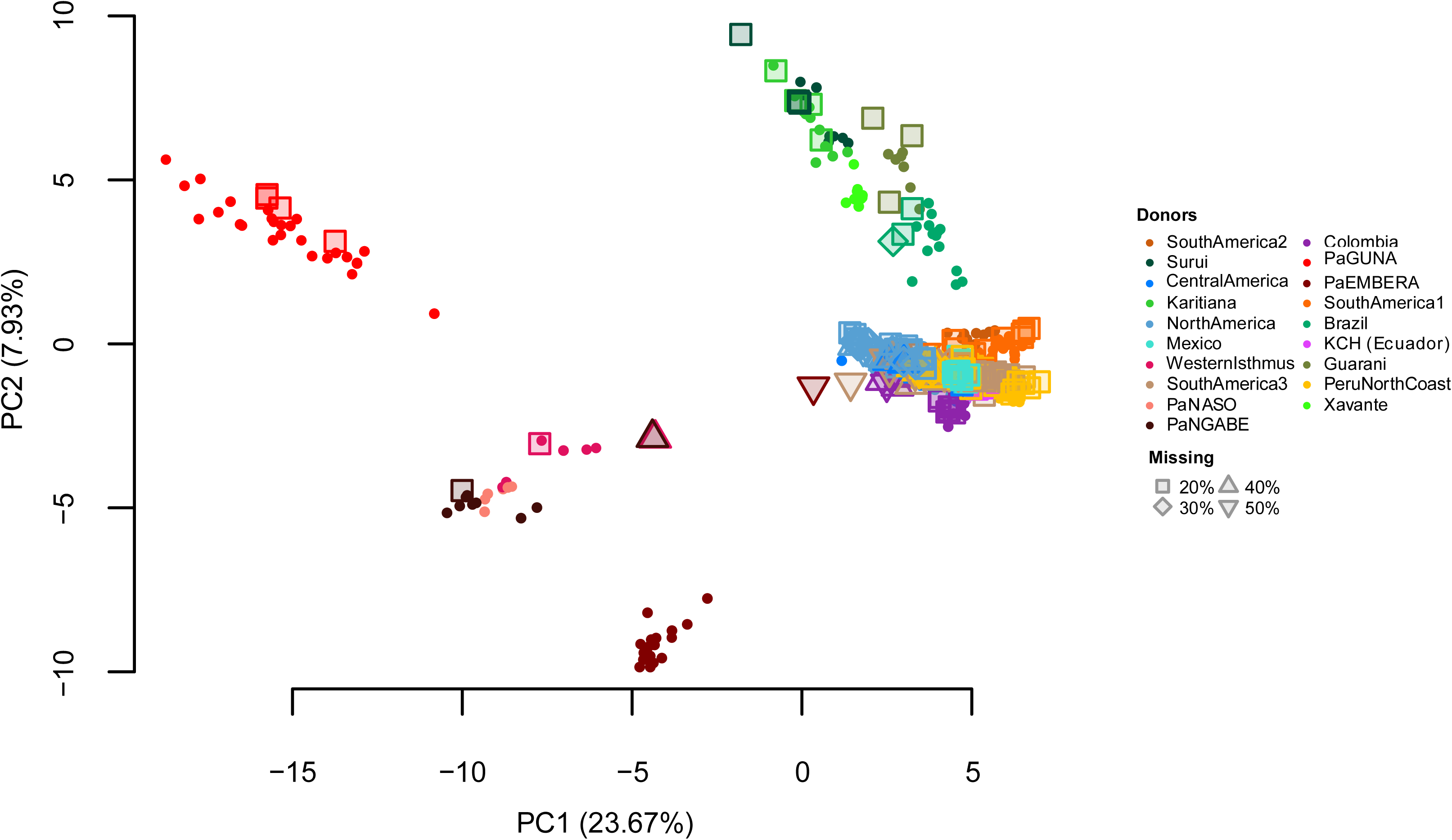
Haplotype-based Indigenous American PCA. Related to Figures 2B and 3A. The PCA was built using copying vectors inferred using a modified version of ChromoPainter allowing for the presence of missing data. The masked individuals (rmIA311) have been projected on the variability of the nearly unadmixed individuals (uIA217) regardless of the level of missing data. The unadmixed individuals are indicated with full filled dots while the masked ones are represented by different shapes, according to the percentage of missing SNPs. The colors refer to the clusters (donors) of Figure 3A.

**Figure S15.**
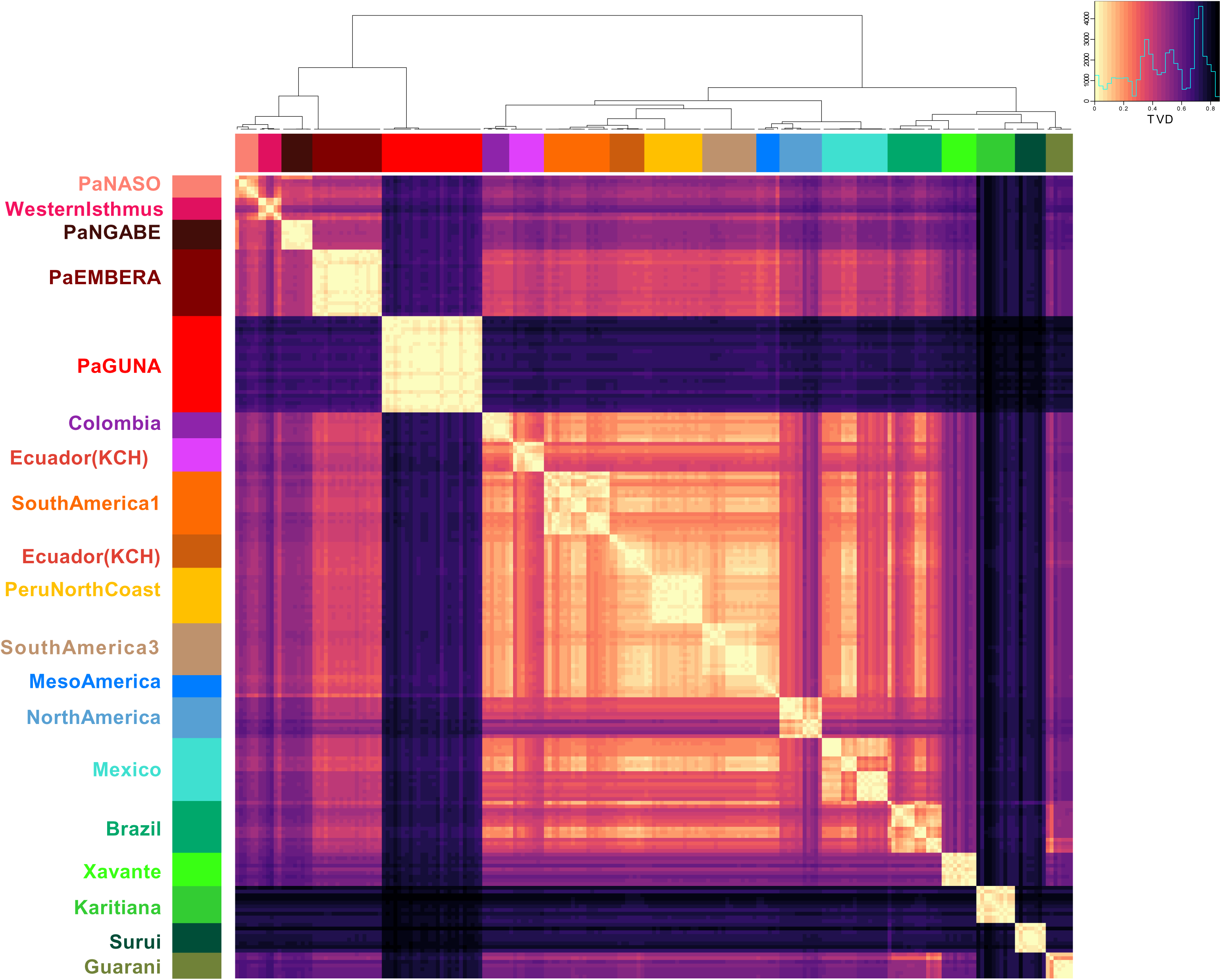
Heatmap Based on Individual TVD Values. Related to Figure 3A. Dendrogram branches are colored according to 19 clusters (Figure 3A). The Total Variation Distance (TVD) was compared both among and within clusters. Lighter colors (lower TVD values) in the matrix mean similarity, while darker colors (higher TVD values) indicate heterogeneity.

**Figure S16.**
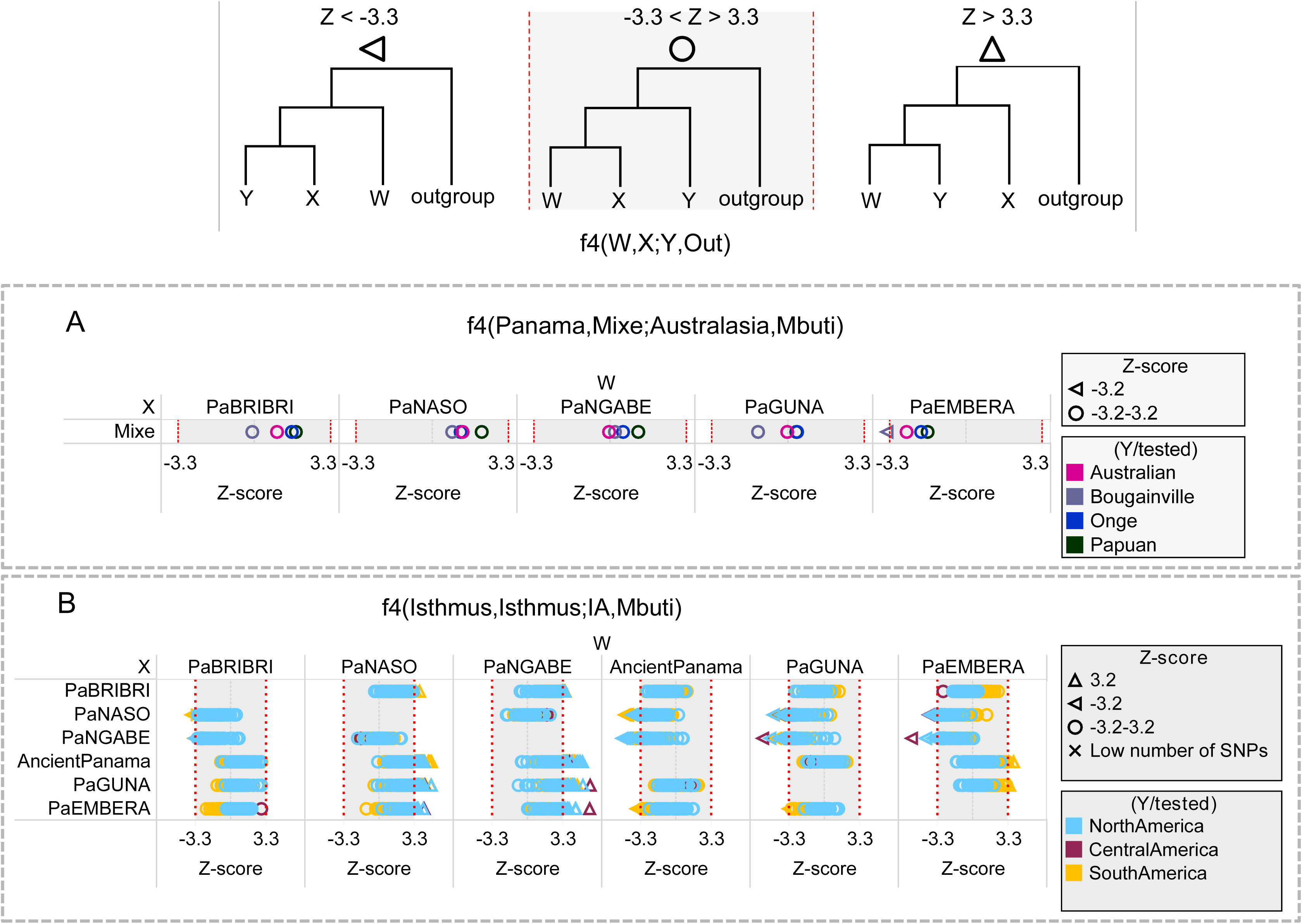
*f4-statistics* Involving Panama Individuals. Related to Figure 2C. The grey region represents the *Z-score* (between -3.3 and 3.3.) where the tested conformation of the tree is confirmed with *p ∼0.001*. The analyses were performed on uIA89, mIA417 plus ancient Panamanians considering all SNPs. A) The Panamanian populations (X) were compared to Mixe (W), typically used to reveal pop-Y among IA, and to three Australasian populations (Y). Then, B) each Panamanian Indigenous group (W) was compared to the others (X) considering the IA populations (coloured according to their geographic location) to test through the *Z-score* whether a given Panamanian group carries excess of a specific IA ancestry compared with others.

**Figure S17.**
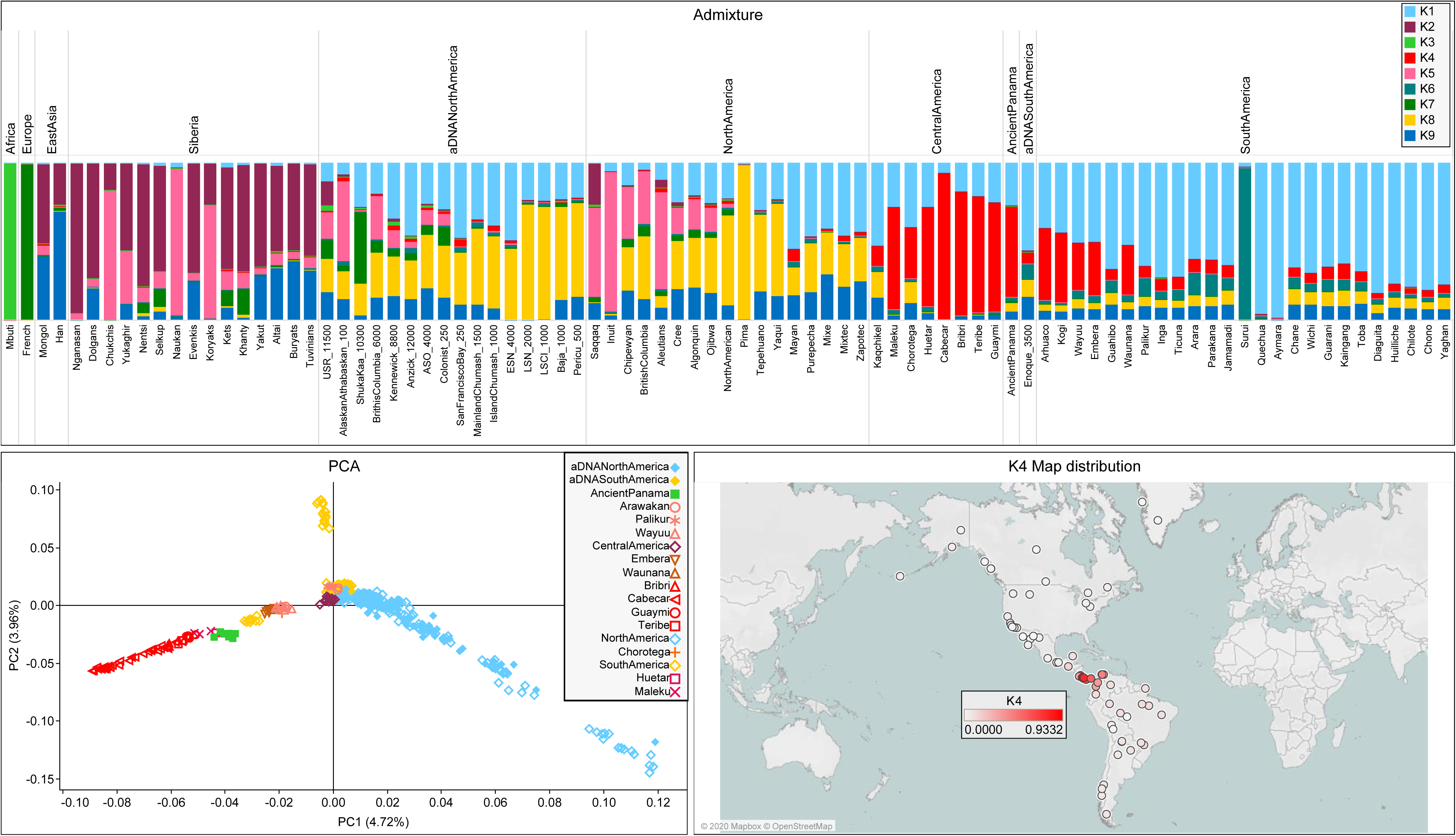
Admixture and PCA on a Comparative Dataset Including Additional Chibchan-speaking Populations. Related to Figure 2. The analyses were performed on a comparative dataset from Scheib et al. 2018 (Table S5) that includes the following Chibchan-speaking populations: Arhuaco and Kogi from Colombia; Guaymi, Cabecar, Teribe, Bribri, Huetar, and Maleku from Costa Rica.

**Figure S18.**
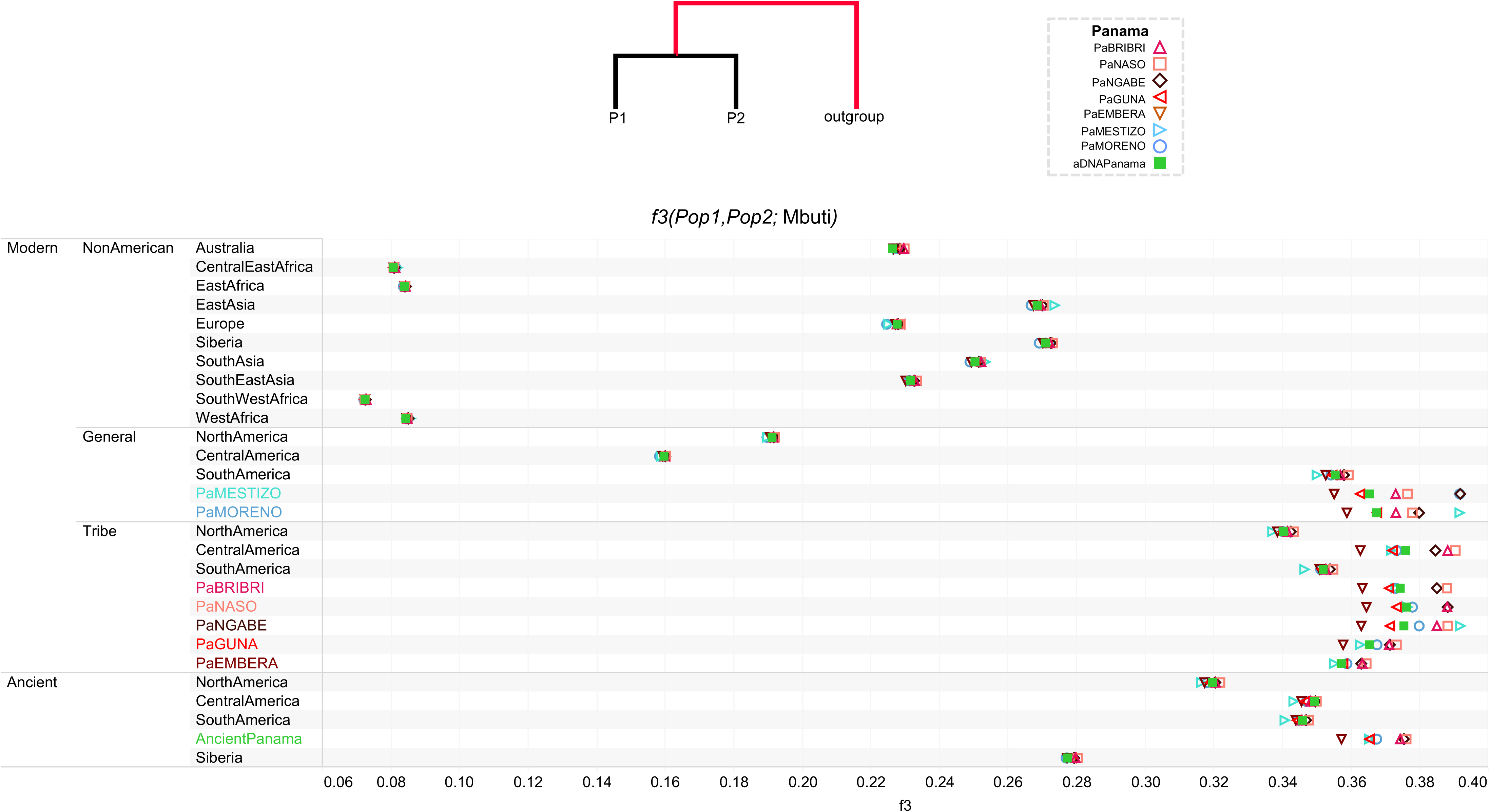
Worldwide Outgroup *f3-statistics* of Panama Individuals. Related to Figure 4A. The Panamanian groups (W) where compared to worldwide population (X) including non-American populations in the rWD1560 dataset, all populations in the mIA417 and uIA217 datasets and all ancient individuals. All comparisons have a *Z-score* > 32.912. The average *f3* value for each population is reported in abscissa.

**Figure S19.**
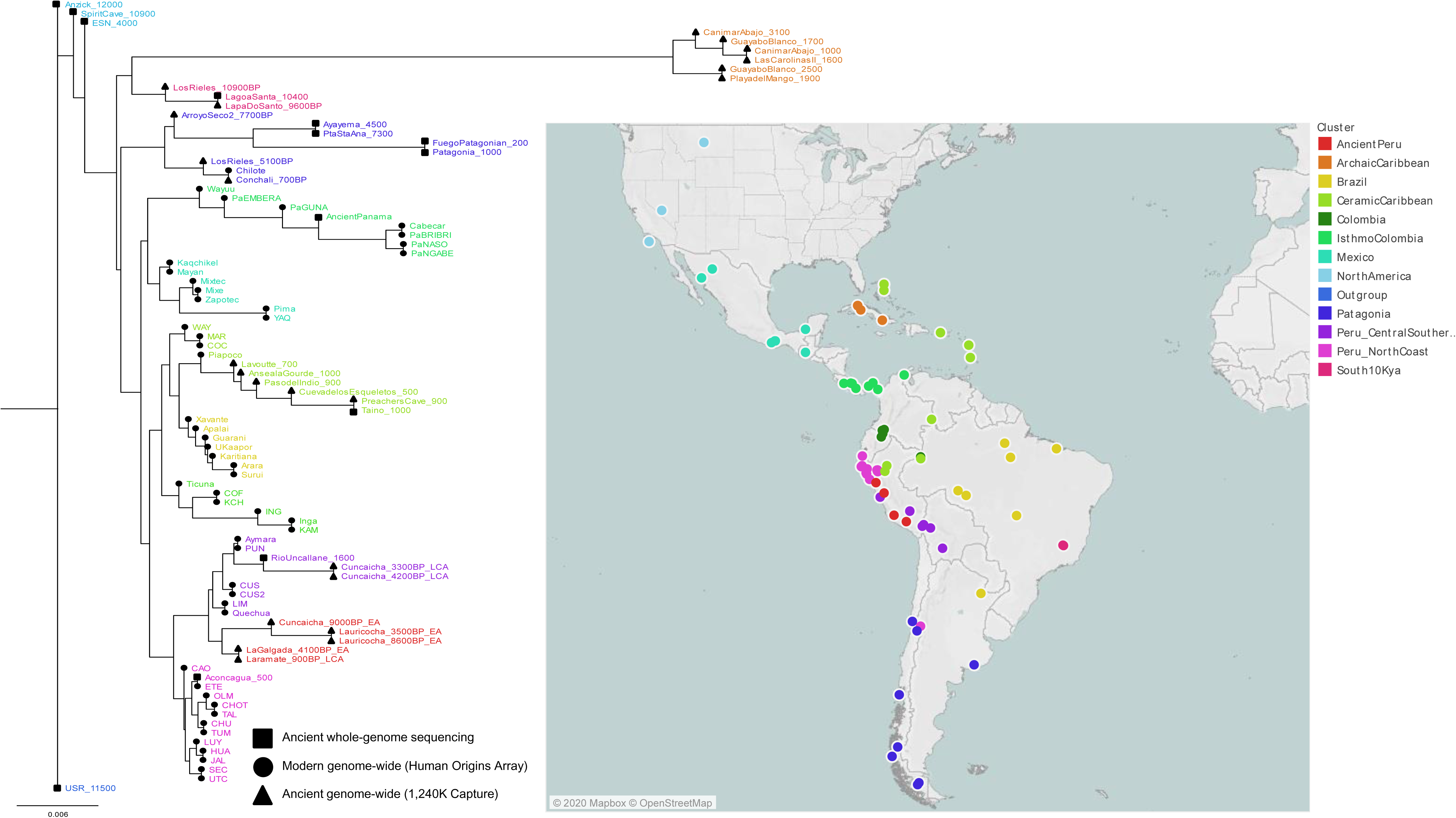
Neighbor-joining Tree Based on Inverted Outgroup *f3-*Statistics. Related to Figures 3A, 4A, and 5A. The tree is built using the inverse values derived from the outgroup *f3*-statistics on all Central and South American populations pairs plus Anzick-1, Early San Nicolas (ESN), Spirit Cave and USR. The latter is considered as an outgroup in the tree. We retained only populations with more than 30K overlapping SNPs and significant Z-scores (>3.3) in all comparisons. The map shows the geographic distribution of the populations, which are colored according to their genetic proximity in the tree.

**Figure S20.**
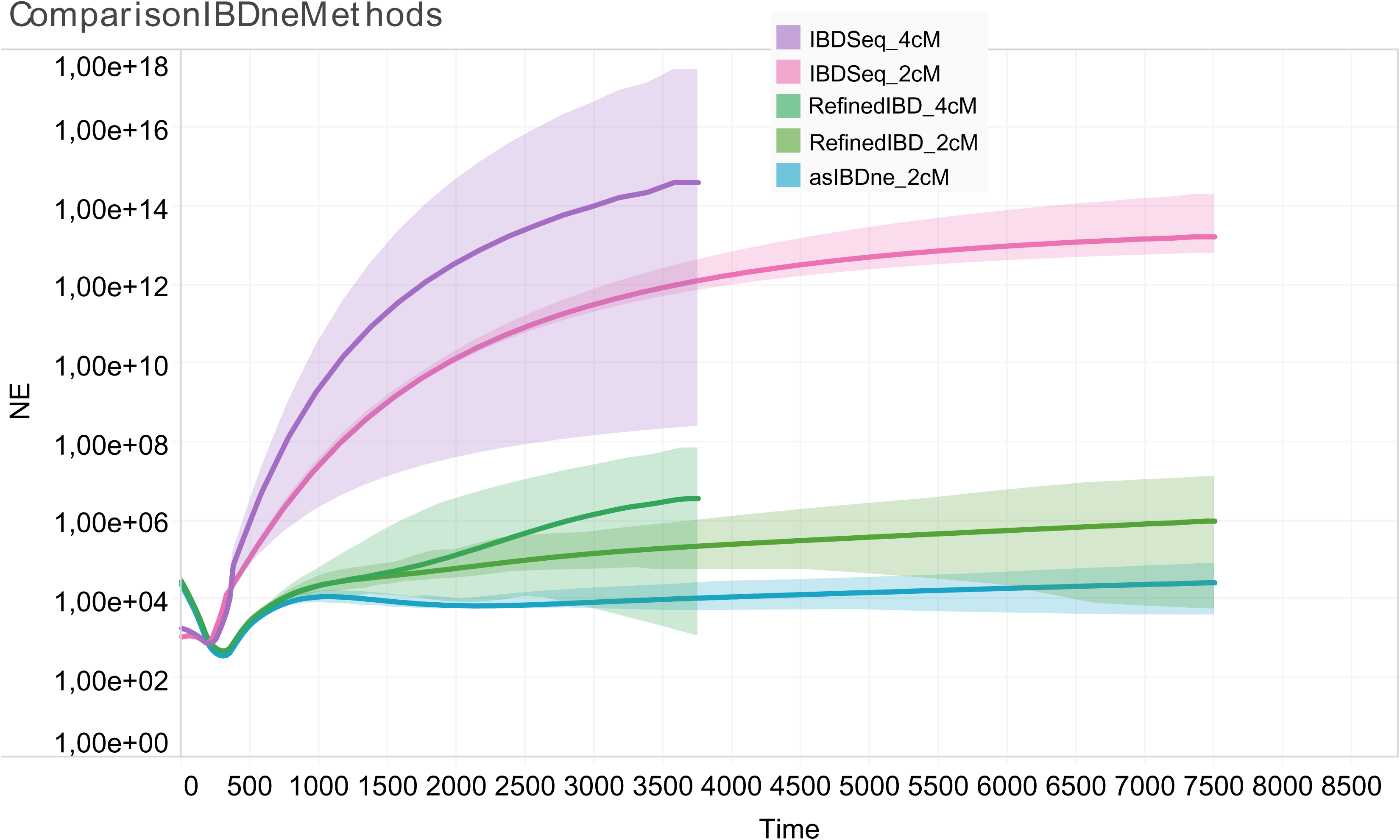
Estimating Effective Population Size by Using IBD Segments Inferred from Different Tools. Related to Figure 3C. We compared IBDseq, which does not require phased data, with the haplotype-based method RefinedIBD because switch errors in estimated haplotypes can erroneously break long IBD segments into shorter sub-segments. The colored regions show 95% confidence intervals. The results for the uIA217 dataset are shown considering different IBD-segment lengths (2cM and 4cM). Ancestry-specific effective population size was also considered for IBD segments larger than 2cM (asIBDne).

**Figure S21.**
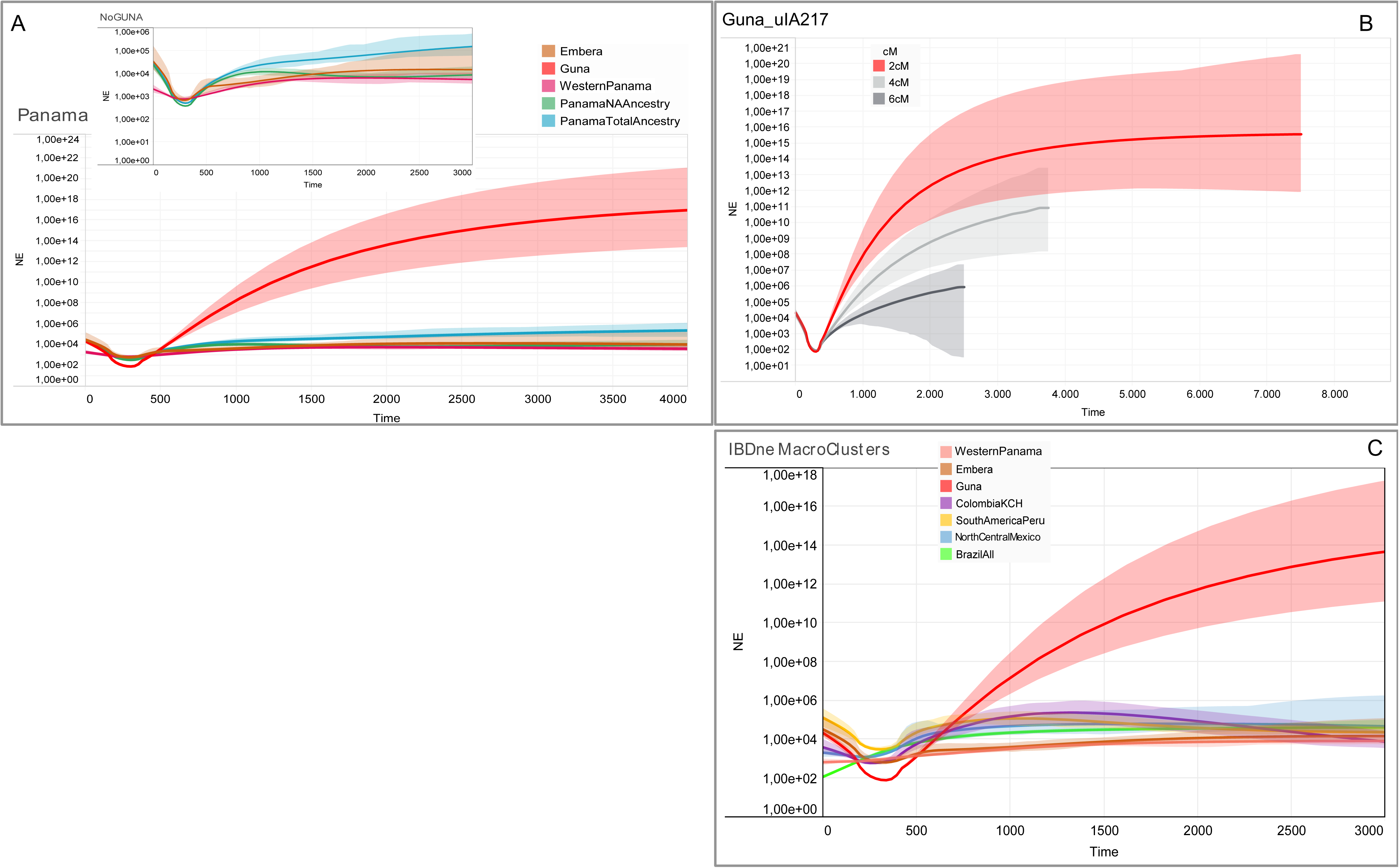
Estimating Effective Population Size of Panama Over Time. Related to Figure 3C. We used RefinedIBD with a threshold of 2 cM on inferred IBD length. The y-axes show effective population size (Ne), plotted on a log scale and the colored regions show 95% bootstrap confidence intervals (CI). The x-axes show the time before the present as years ago (ya) considering a generation time of 25 years. A) We evaluated the entire Panamanian dataset (included in uIA217), considering: i) all ancestries, ii) only the Indigenous ancestries and iii) the three macro-clusters identified in Panama (Emberà, Guna, and North Panama, made of North Isthmo, Naso, and Ngäbe (Figure 3A)). The inset magnifies the same data without Guna, that shows a large CI due to the lack of short IBD fragments as shown in B), where Ne of Guna is evaluated considering different IBD thresholds (2 cM, 4 cM and 6 cM). C) We also compared the IBDne of the Panamanian macro-clusters with the others identified in Figure 3A (WesternPanama = WesternIsthmo, PaNASO, PaNGABE; Emberà = PaEMBERA; Guna = PaGUNA; ColombiaKCH = Colombia, Ecuador(KCH); SouthAmericaPeru = SouthAmerica1, SouthAmerica2, PeruNorthCoast, SouthAmerica3; NorthCentralMexico = CentralAmerica, NorthAmerica, Mexico; BrazilAll = Brazil, Xavante, Karitiana, Surui, Guarani).

**Figure S22.**
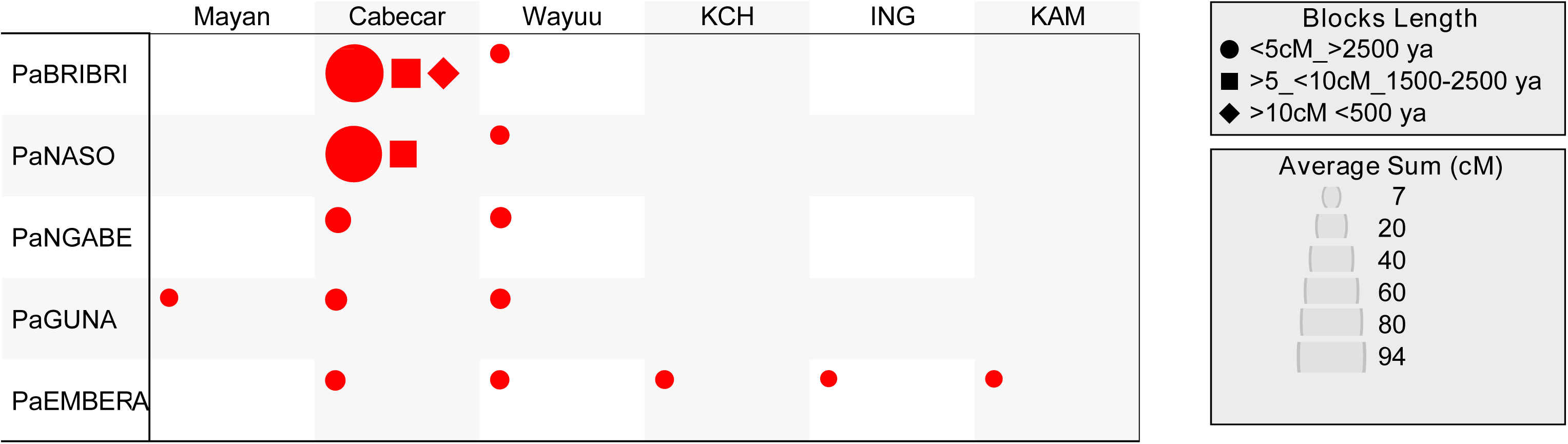
IBD Sharing Between Populations. Related to Figure 3C. Visualization of the average of summed IBD lengths shared between modern Panamanians and other IA populations in each paired comparison, with identified IBD blocks in the range of 1–5 cM (oldest), 5–10 cM, and over 10 cM (youngest). Shape sizes are proportional to mean values; only those pairs sharing at least two blocks >5 cM and four <5 cM are plotted.

**Figure S23.**
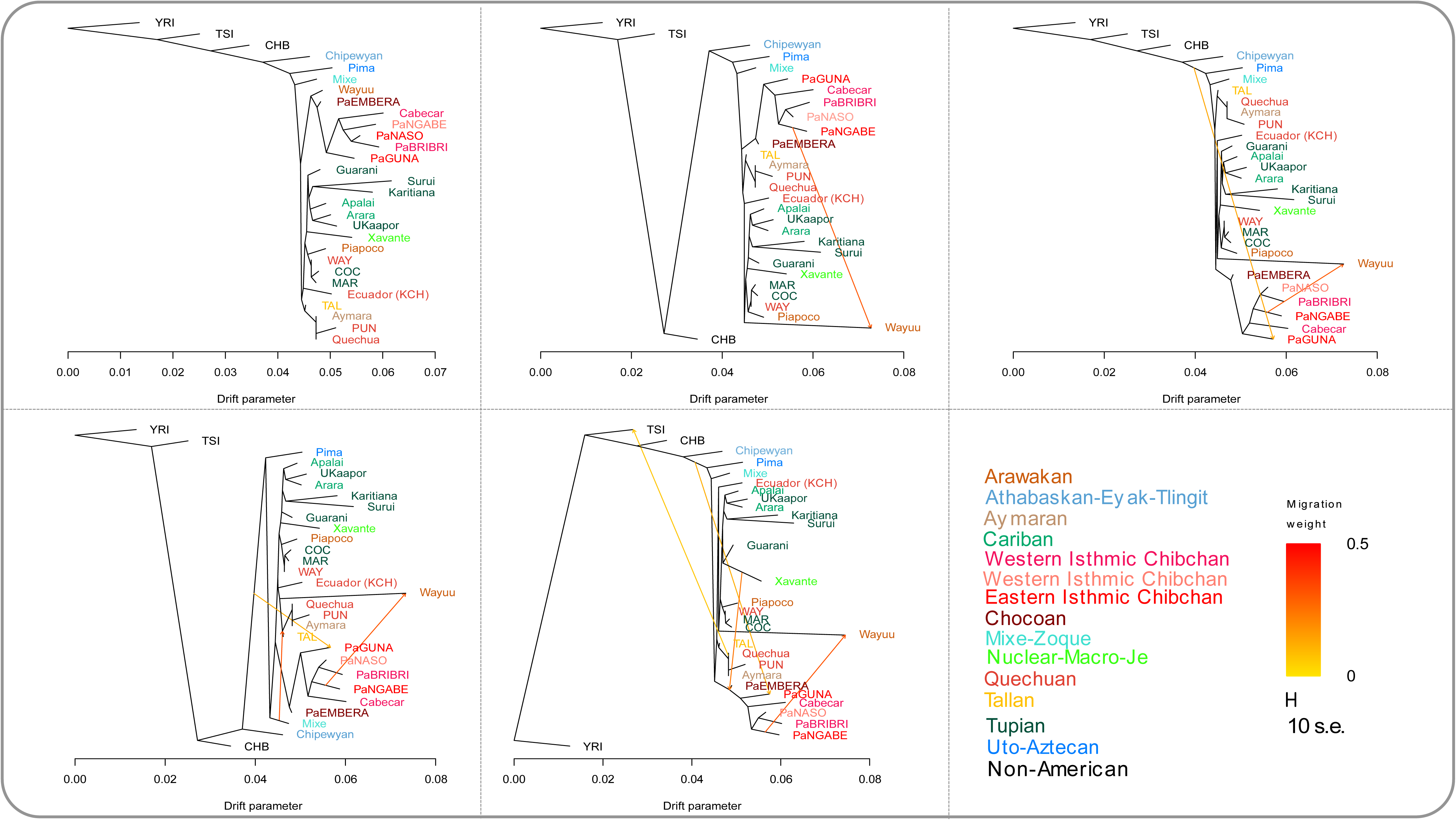
Inferred Maximum Likelihood Tree with Admixture Events. Related to Figure 5A. Plotted is the structure of the graph inferred by TreeMix Trees for the dataset uIA89, using Yoruba (YRI), Tuscan (TSI) and Han (CHB) as non-Indigenous American outgroups, and allowing from zero to five admixture edges (migration events). Population groups are colored according to linguistic/geographic affiliation. Horizontal branch lengths are proportional to the amount of genetic drift that has occurred on the branch. Migration arrows are colored according to their weight. The scale bar shows ten times the average standard error of the entries in the sample covariance matrix (Pickrell and Pritchard, 2012).

**Figure S24.**
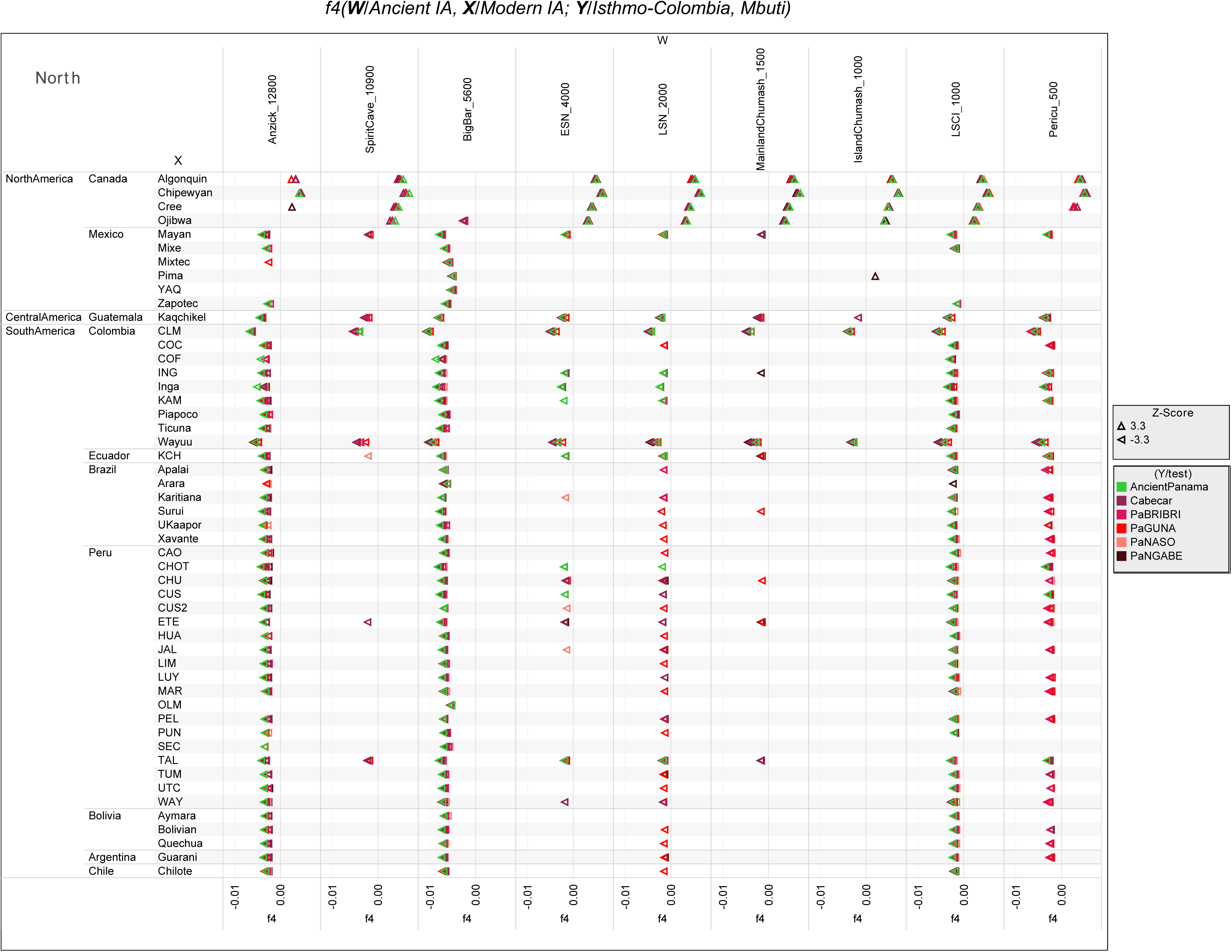
*f4-statistics* Testing the Genetic Affinity Between Isthmian Groups, Modern IA and Ancient IA Genomes from North America. Related to Figure 6A.

**Figure S25.**
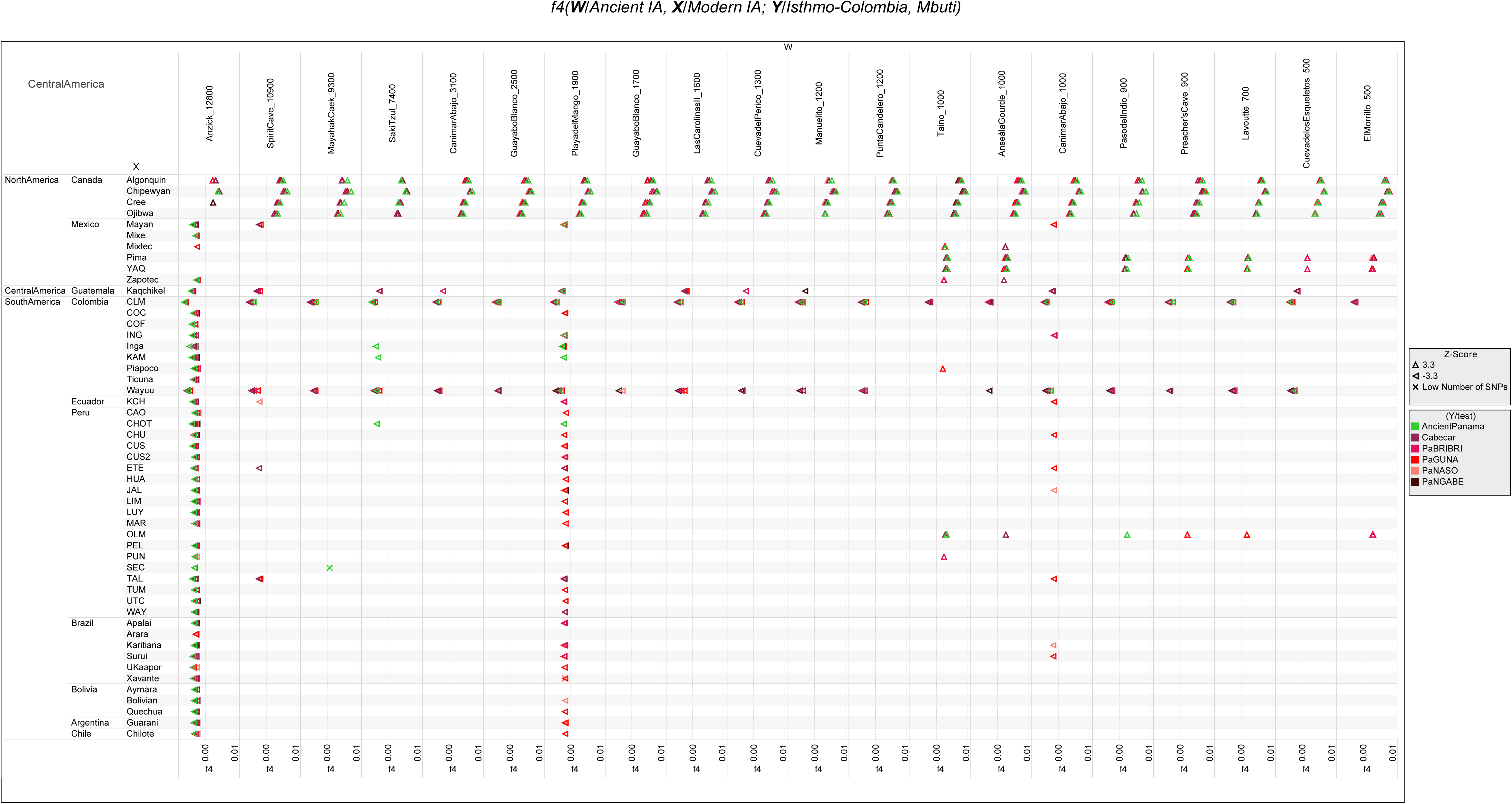
*f4-statistics* Testing the Genetic Affinity Between Isthmian Groups, Modern IA and Ancient IA Genomes from Central America. Related to Figure 6A.

**Figure S26.**
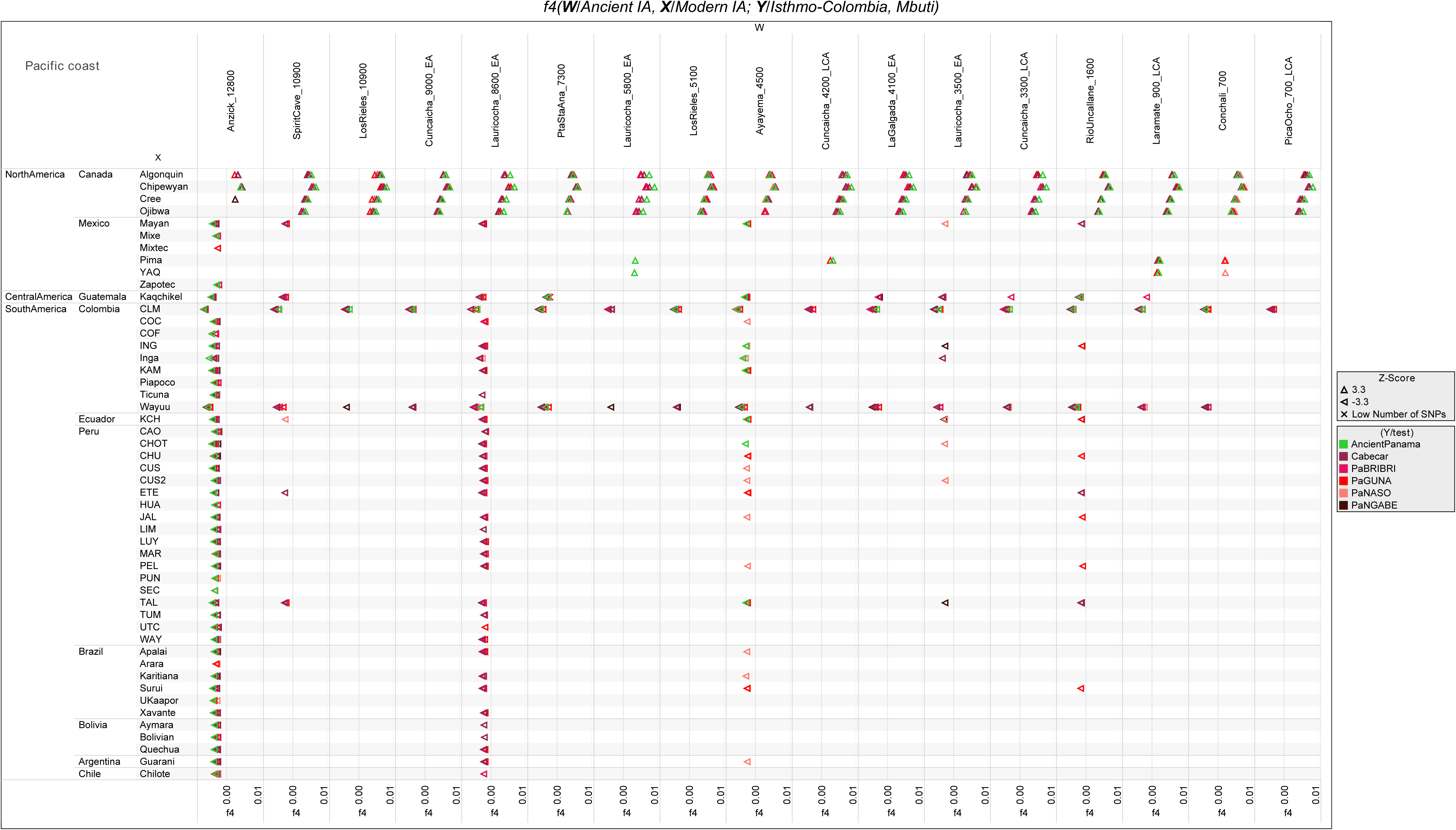
*f4-statistics* Testing the Genetic Affinity Between Isthmian Groups, Modern IA and Ancient IA Genomes from South American Atlantic Coast. Related to Figure 6A. We tested the form *f4*(modern IA, ancient IA; Isthmo, Mbuti) on uIA89, mIA417 and ancient genomes considering only transversions. See Figure 6A for further details.

**Figure S27.**
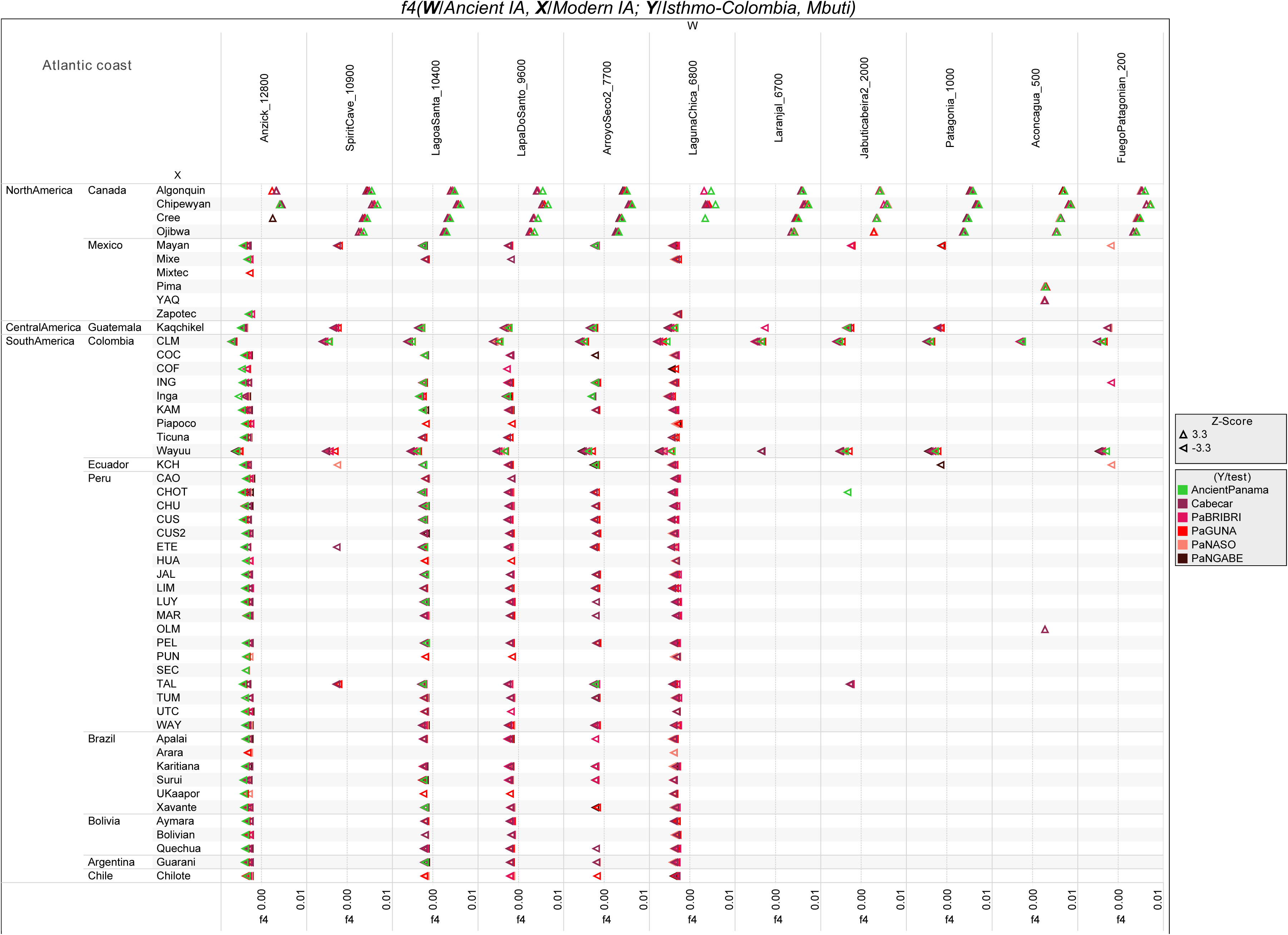
*f4-statistics* Testing the Genetic Affinity Between Isthmian Groups, Modern IA and Ancient IA Genomes from South American Pacific Coast. Related to Figure 6A. We tested the form *f4*(modern IA, ancient IA; Isthmo, Mbuti) on uIA89, mIA417 and ancient genomes considering only transversions. See Figure 6A for further details.

**Figure S28.**
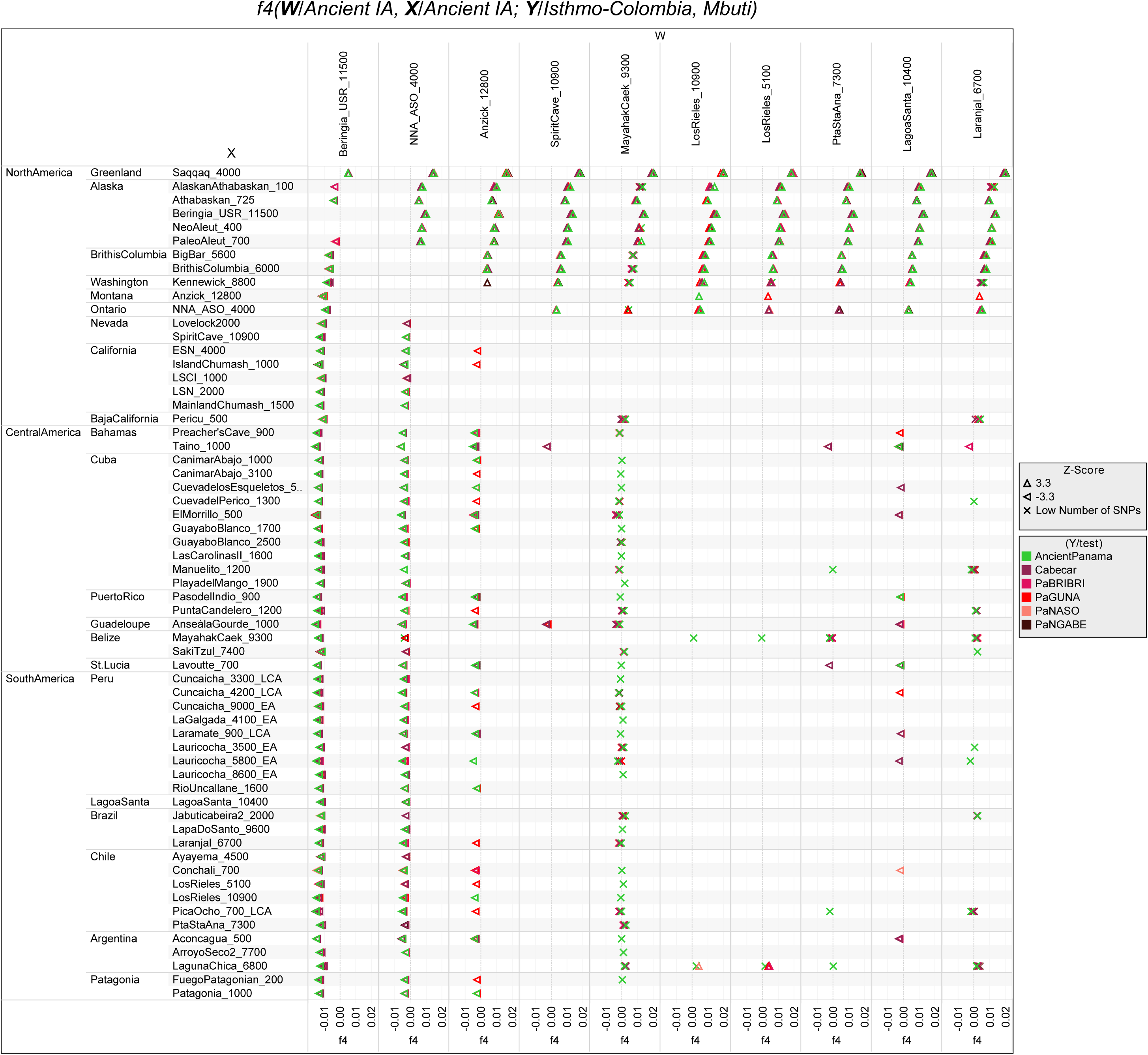
*f4-statistics* Testing the Genetic Affinity Between Isthmian Groups and Different Ancient IA Genome Pairs. Related to Figure 6A. We tested the form *f4*(ancient IA, ancient IA; Isthmo, Mbuti) on uIA89, mIA417 and ancient genomes considering only transversions. See Figure 6A for further details.

**Figure S29.**
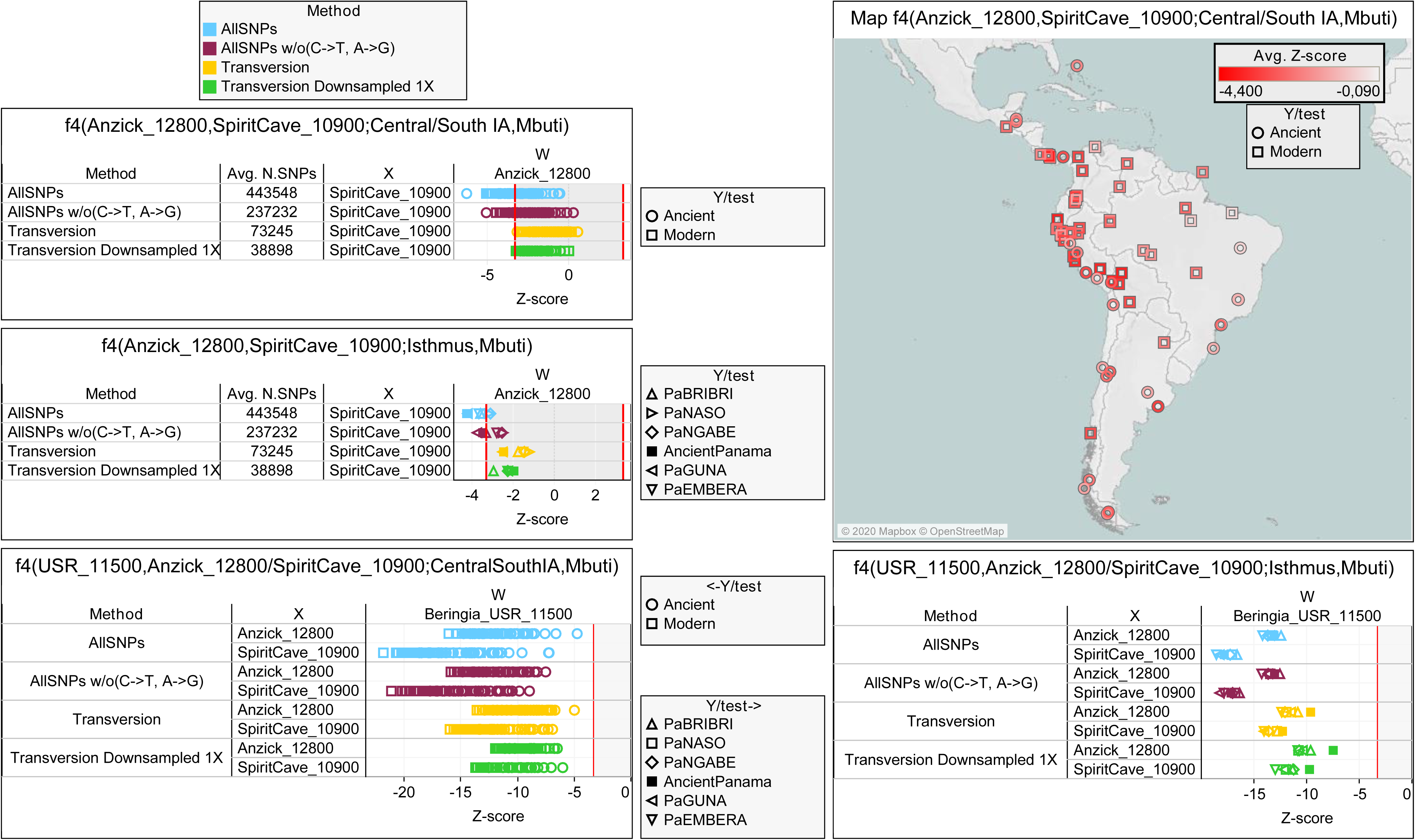
*f4-statistics* Testing the Relationships of Isthmian and Other Central/South American Populations with Anzick-1 and Spirit Cave. Related to Figure 2C. We run the f4-statistics in the following forms (Anzick-1, Spirit Cave; Isthmo, Mbuti) and (Anzick-1, Spirit Cave; Central and South IA, Mbuti). The datasets uIA89, mIA417 and ancient individuals were used considering different sets of variants.

**Figure S30.**
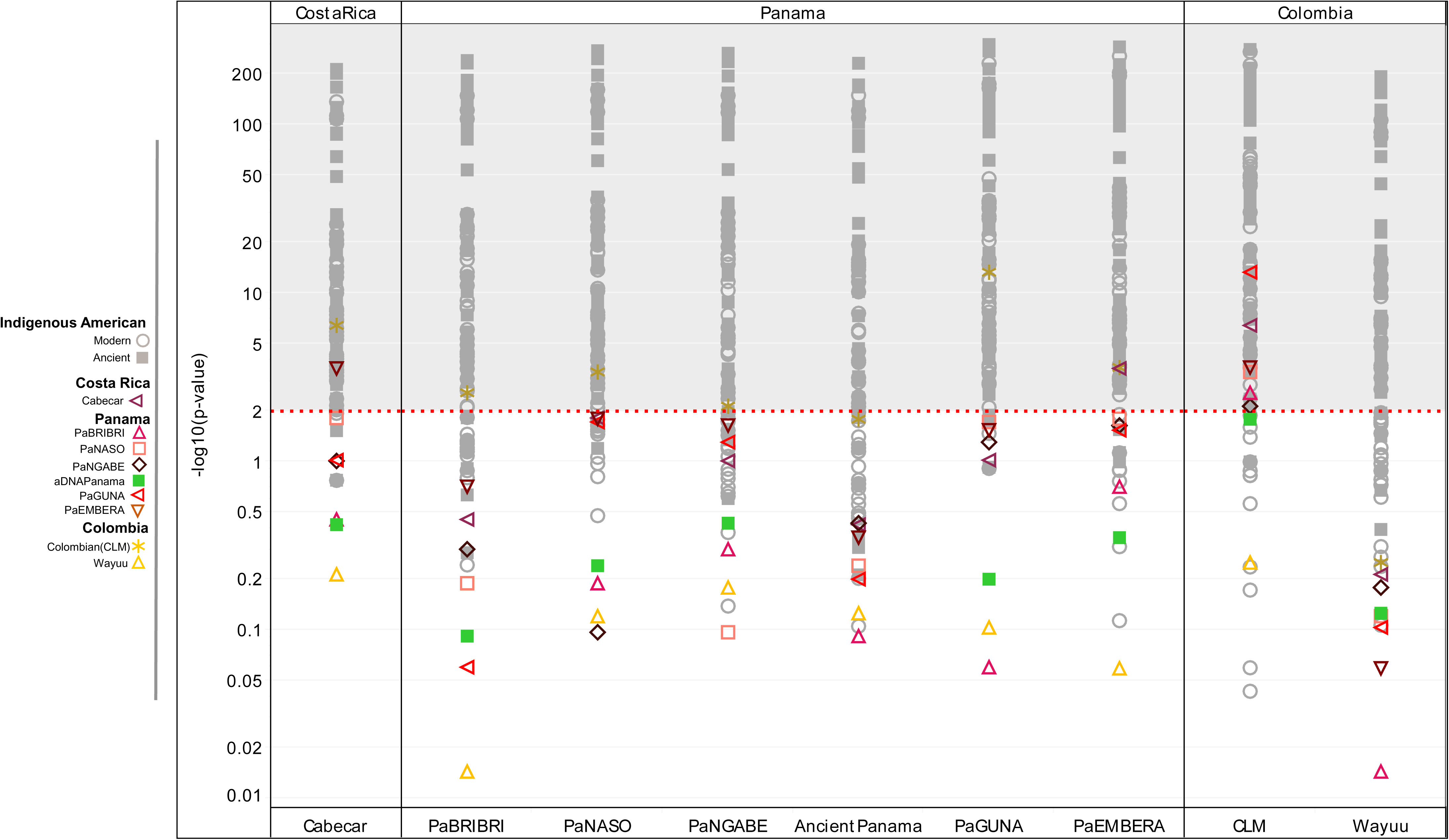
Minimum Number of Ancestral Sources for the Isthmian Populations (Considering Rank 1). Related to Figure 6. qpWave analyses where we compared in pairs Ancient Panama and present-day Isthmian groups with all IA populations. The outgroups were kept to the minimum and chosen to represent different IA ancestries identified here (e.g. in the Neighbor-joining Tree of Figure S20) and in other papers. Rank 1 refers to a model in which all paired-populations fit as derived from two ancestral sources, relative to the outgroups. A *p-value*>0.01 (2 in –log10 scale, dotted red line) means pairs that could be explained by a single independent source.

**Figure S31.**
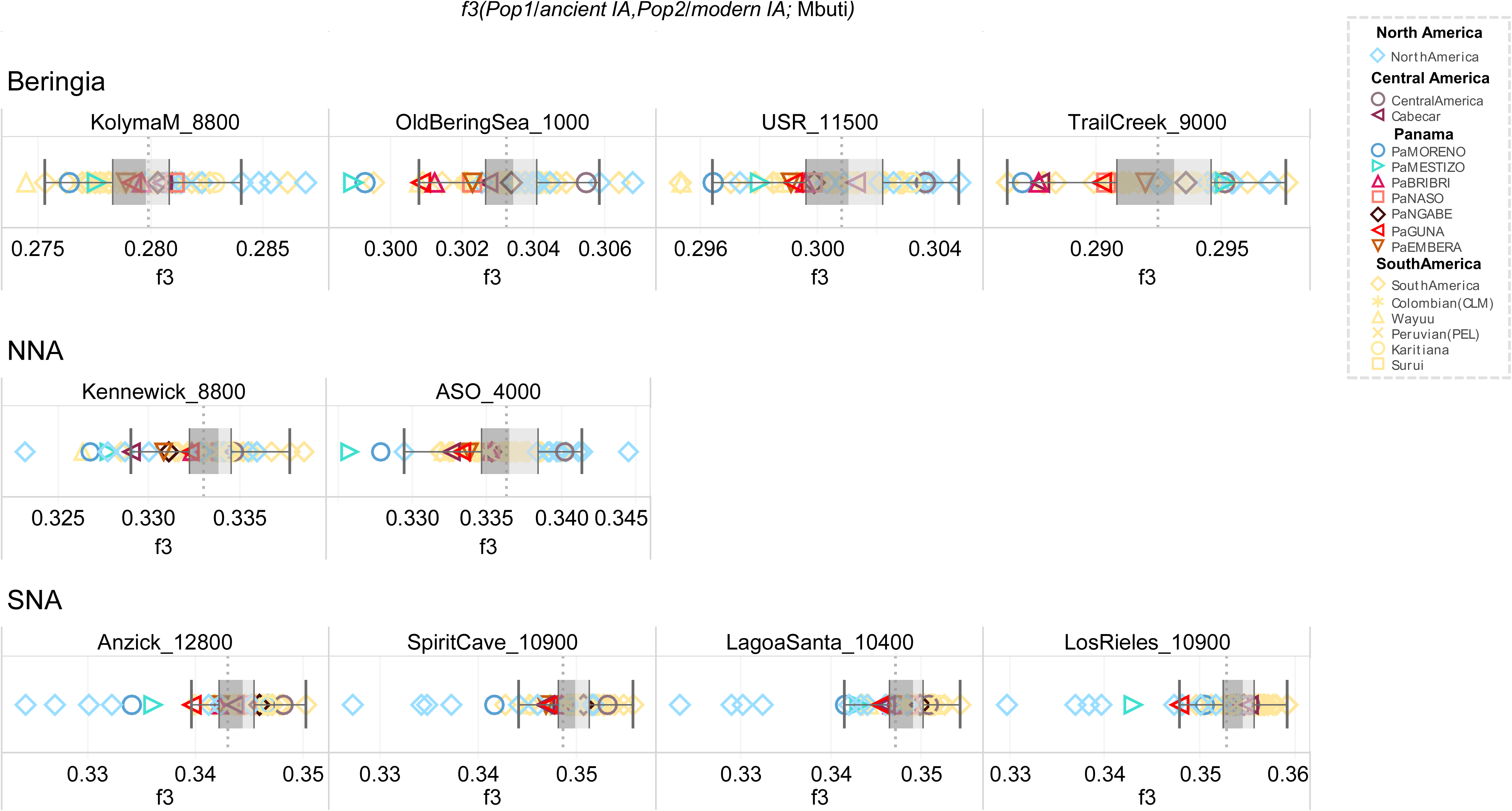
Outgroup *f3-statistics* of Modern Indigenous Americans. Related to Figure 4A. We analyzed the shared genetic history of modern IA populations (included in mIA417 and uIA89) against ancient reference genomes from Beringia and the Americas (representative of the NNA and SNA ancestries).. Box Plots in grey help to visualize the distribution of *f3* values in each comparison, indicating the median (most typical) value, 25^th^ and 75^th^ percentiles (dark and light grey, respectively), and arms extending 1.5 times the IQR (interquartile range). The dotted line indicates the *f3* average value.

**Figure S32.**
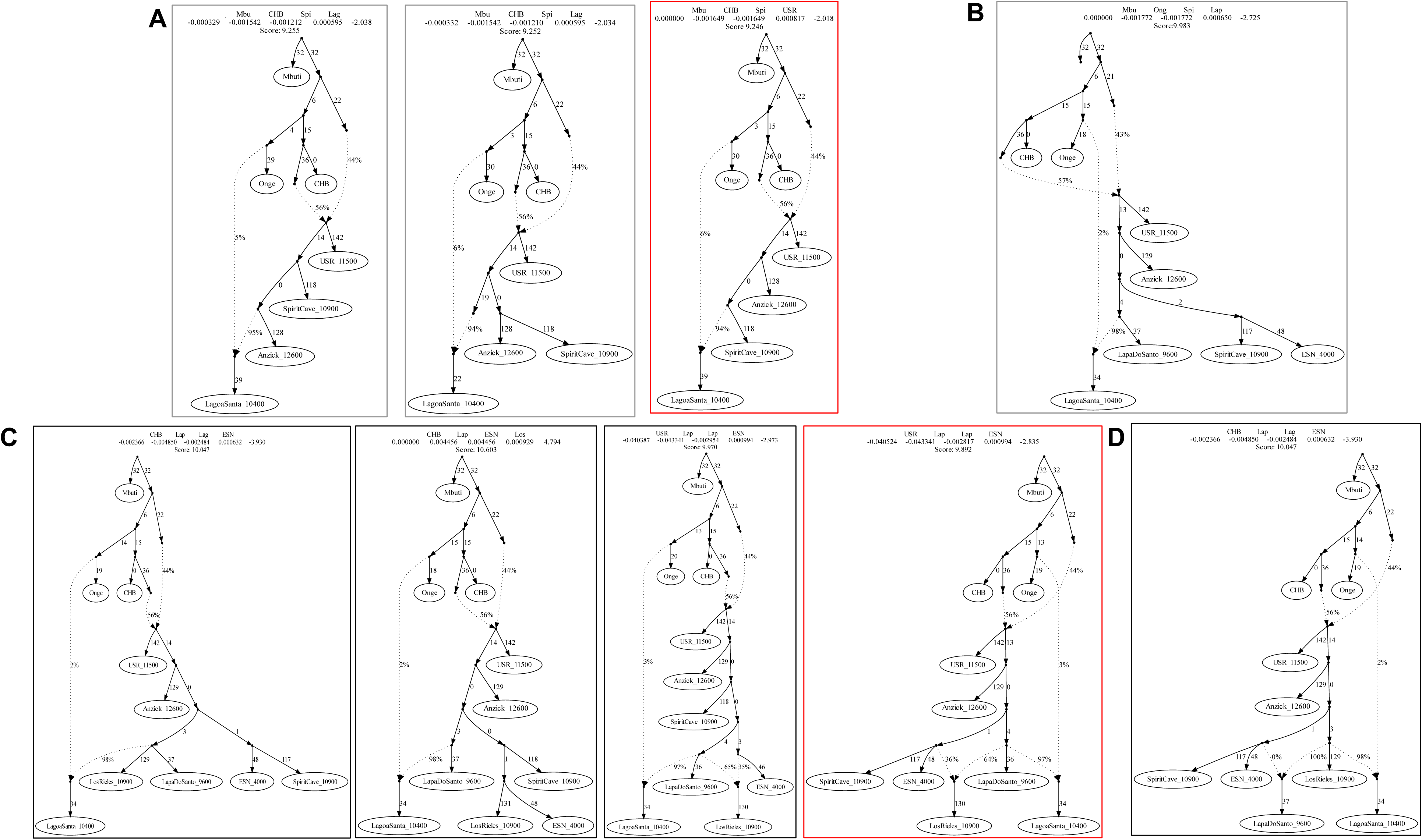
Admixture Graphs Modelling Ancient Individuals Representative of the SNA ancestry. Related to Figure 6B. Basal tree with three of the most ancient available SNA genomes (**A**). The best fitting topology, highlighted in red, was extended: adding Lapa do Santos as SNA1 and ESN as SNA2 (**B**); testing Los Rieles as either unadmixed or admixed (**C**); checking Lapa do Santos as admixed (**D**).

**Figure S33.**
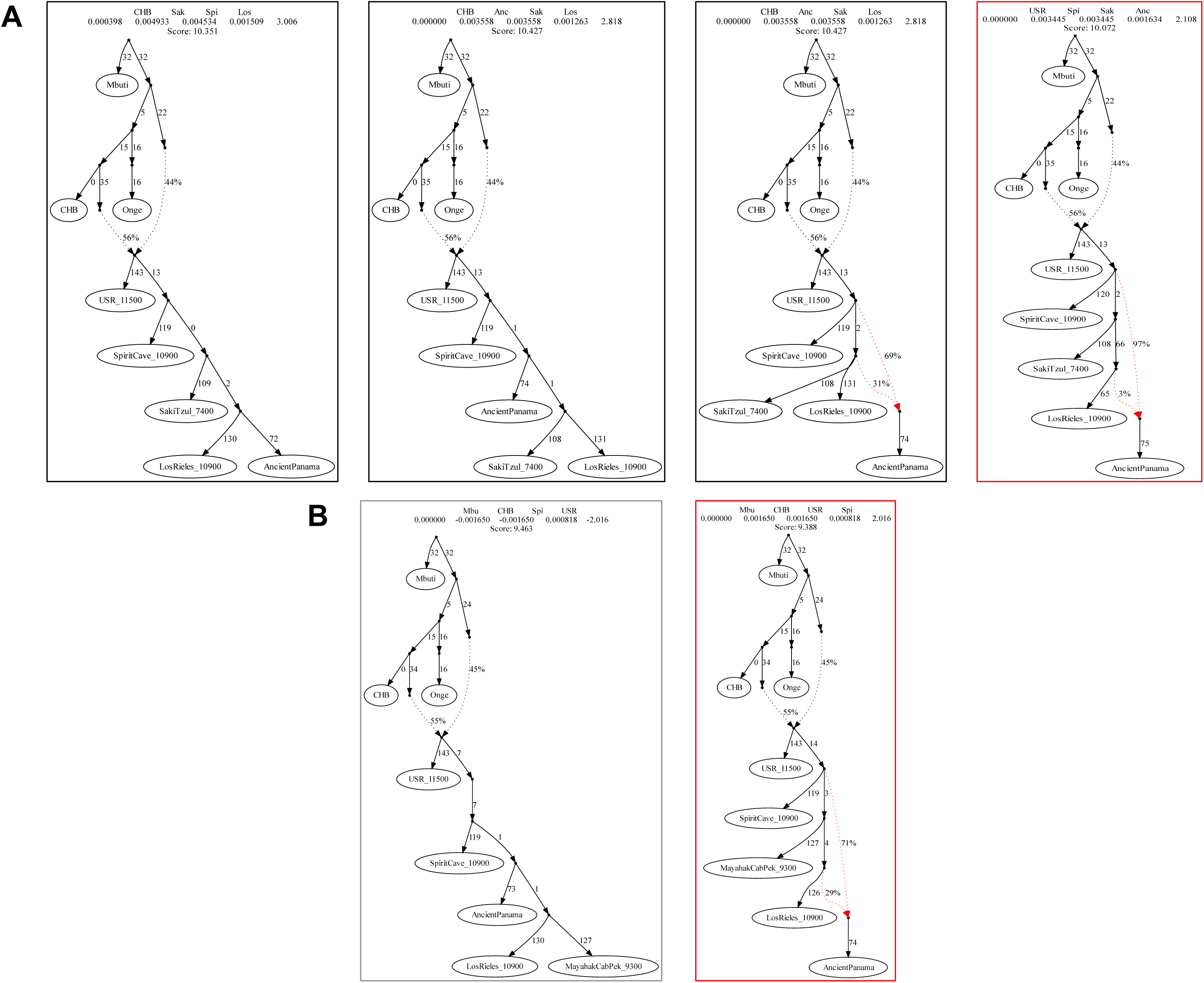
Admixture Graphs Modelling Ancient Panamanians Linked to SNA2 Ancient Genomes. Related to Figure 6B. Possible extensions of the basal SNA2 tree adding ancient Panamanians and other ancient Central American genomes, i.e. Saki Tzul (**A)** and Mayahak Cab Pek (**B**). The best fitting topologies are highlighted in red. See the legend of Figures 6B for further details.

**Figure S34.**
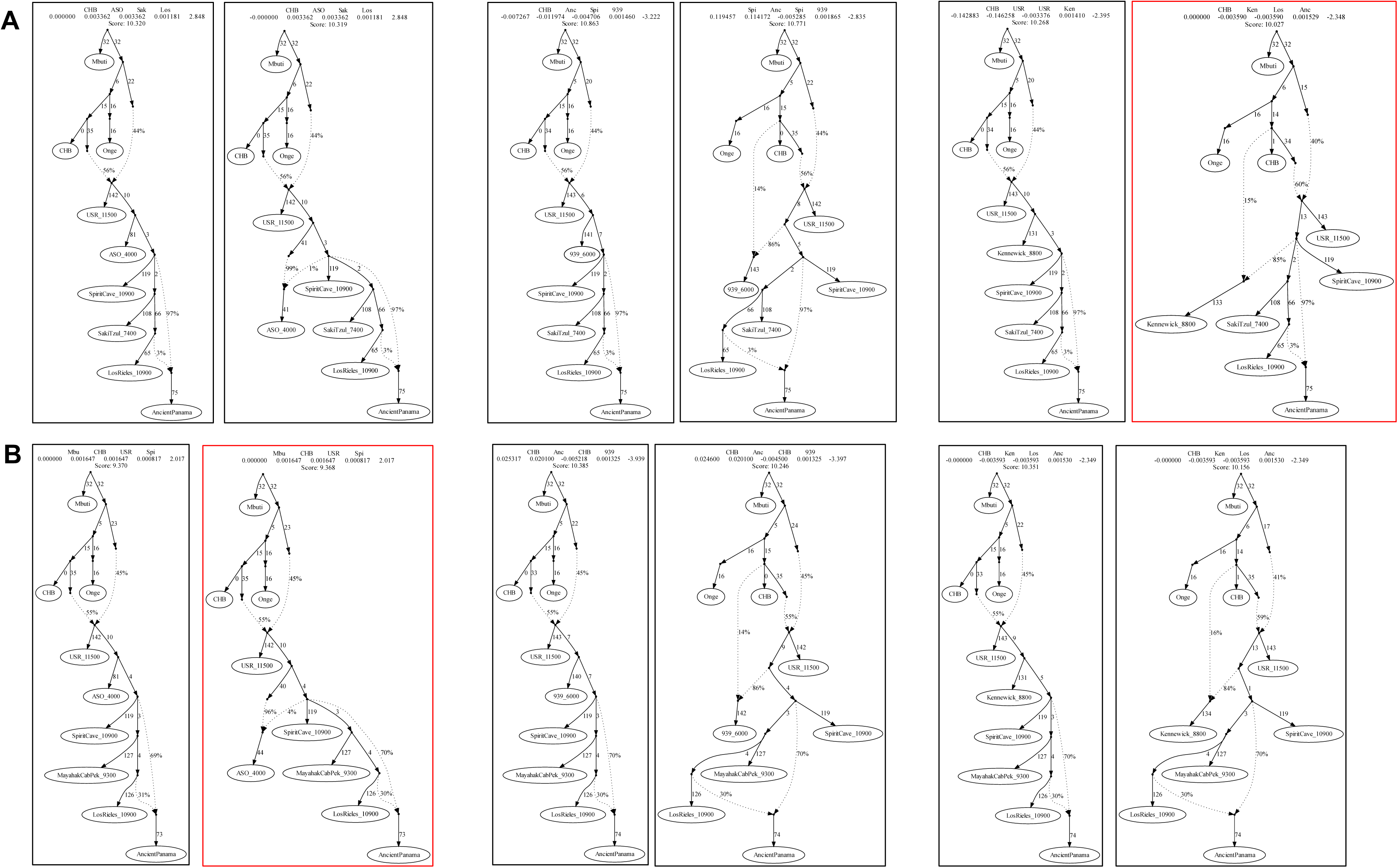
Admixture Graphs Modelling Ancient Panamanians Linked to SNA2 and NNA Ancient Genomes. Related to Figure 6B. Possible extensions of the tree in Figure S33 adding NNA genomes (ASO, Kennewick and 939) to ancient Panama with Saki Tzul (**A**) or with Mayahak Cab Pek (**B**). The best fitting topologies are highlighted in red. See the legend of Figures 6B for further details.

**Figure S35.**
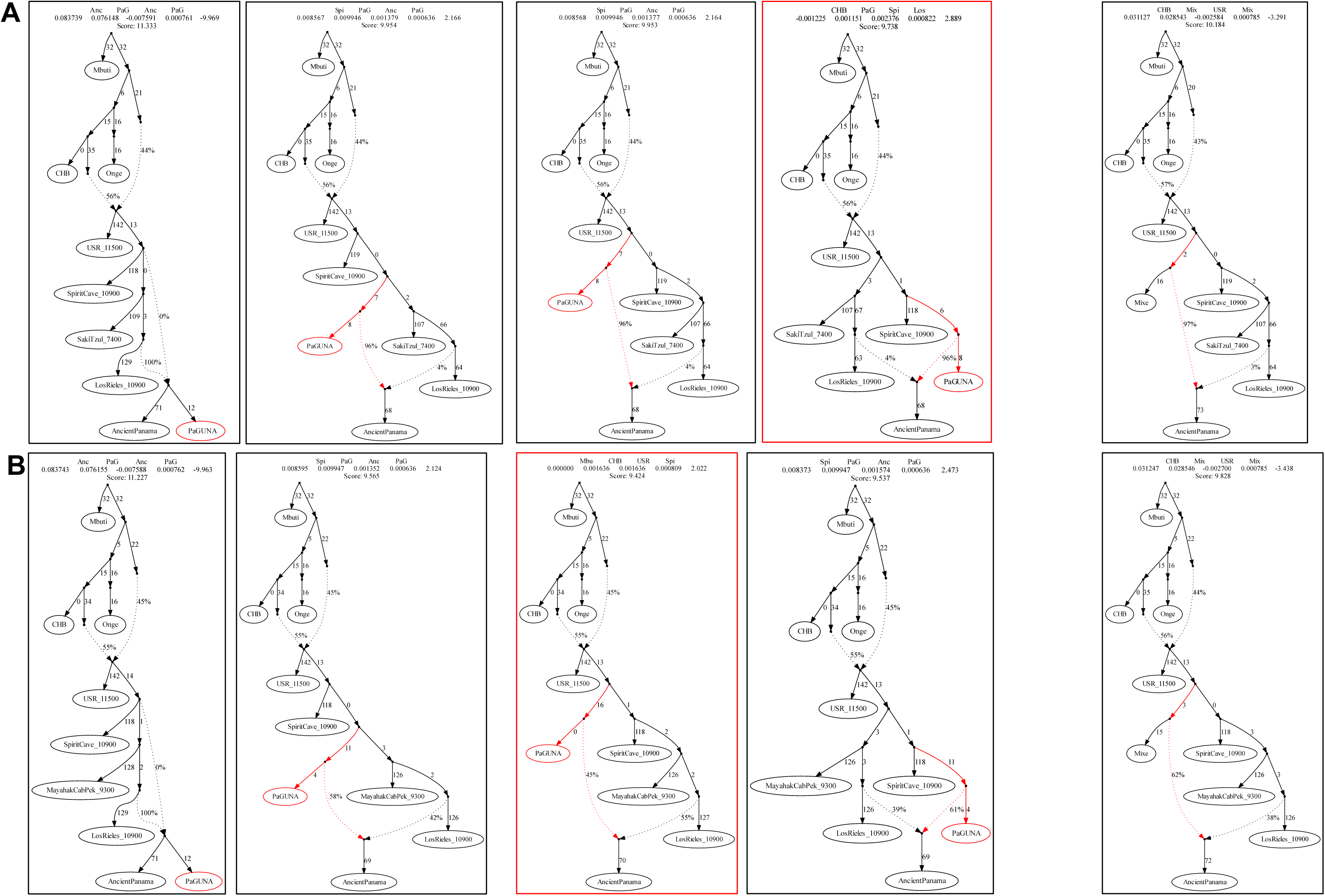
Admixture Graphs Modelling the Panama Genetic History Linked to SNA2 Ancient Genomes. Related to Figure 6B. Possible extensions of the best trees in Figure S33 adding Guna to Saki Tzul (**A**) or Mayahak Cab Pek (**B**). The best fitting topologies are highlighted in red. In the rightmost graphs Guna (UPopI) have been replaced with Mixe (UPopA). See the legend of Figures 6B for further details.

**Figure S36.**
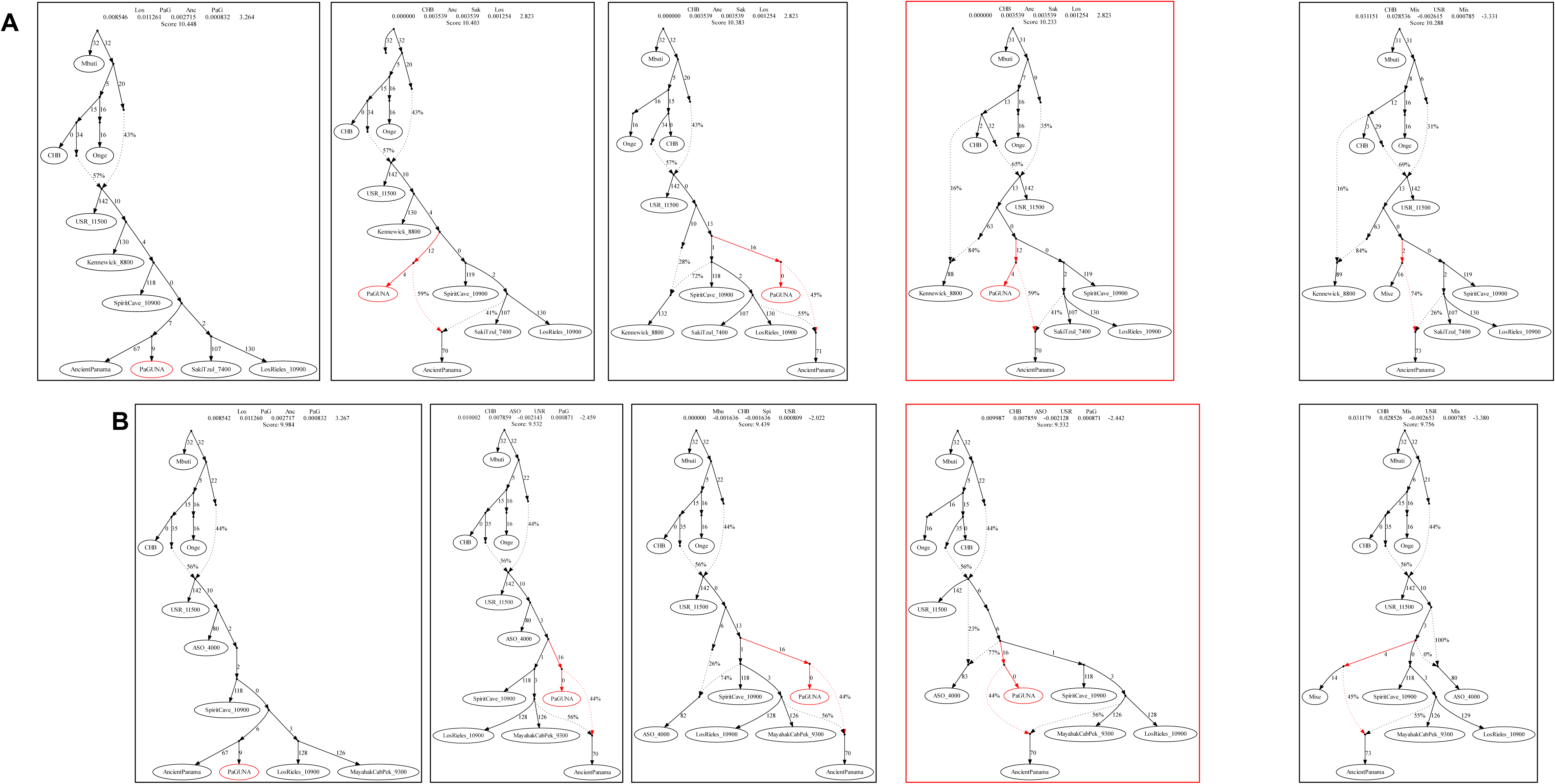
Admixture Graph Modelling the Panama Genetic History Linked to SNA2 and NNA Ancient Genomes. Related to Figure 6B. Possible extensions of the best trees in Figure S34 adding Guna to Saki Tzul with Kennewick (**A**) and Mayahak Cab Pek with ASO (**B**). Best fitting graphs are highlighted in red. In the rightmost graphs Guna (UPopI) has been replaced with Mixe (UPopA). See the legend of Figures 6B for further details.

**Figure S37.**
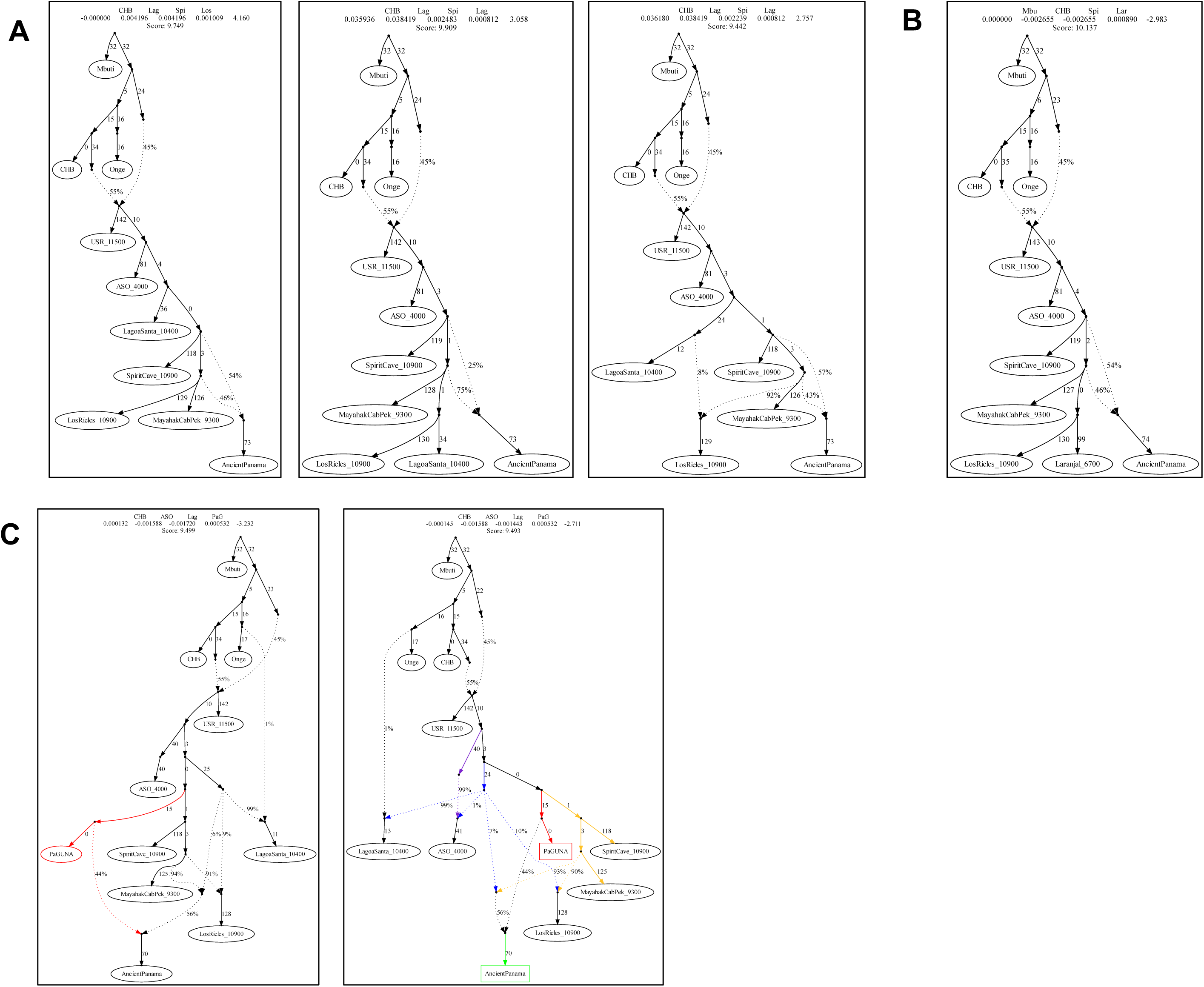
Admixture Graph Modelling the Panama Genetic History Linked to NNA, SNA1 and SNA2. Related to Figure 6B. Possible extensions of the best trees in Figure S34 adding Lagoa Santa (SNA1) together with Ancient Panama (**A**), testing Laranjal instead of Lagoa Santa (**B**) and finally modelling Guna as representative of UPopI (**C**). Best fitting graphs are highlighted in red. In the rightmost graphs we tested Mixe (UPopA) instead of Guna (UPopI) and Anzick-1 instead of Spirit Cave. See the legend of Figures 6B for further details.

**Figure S38.**
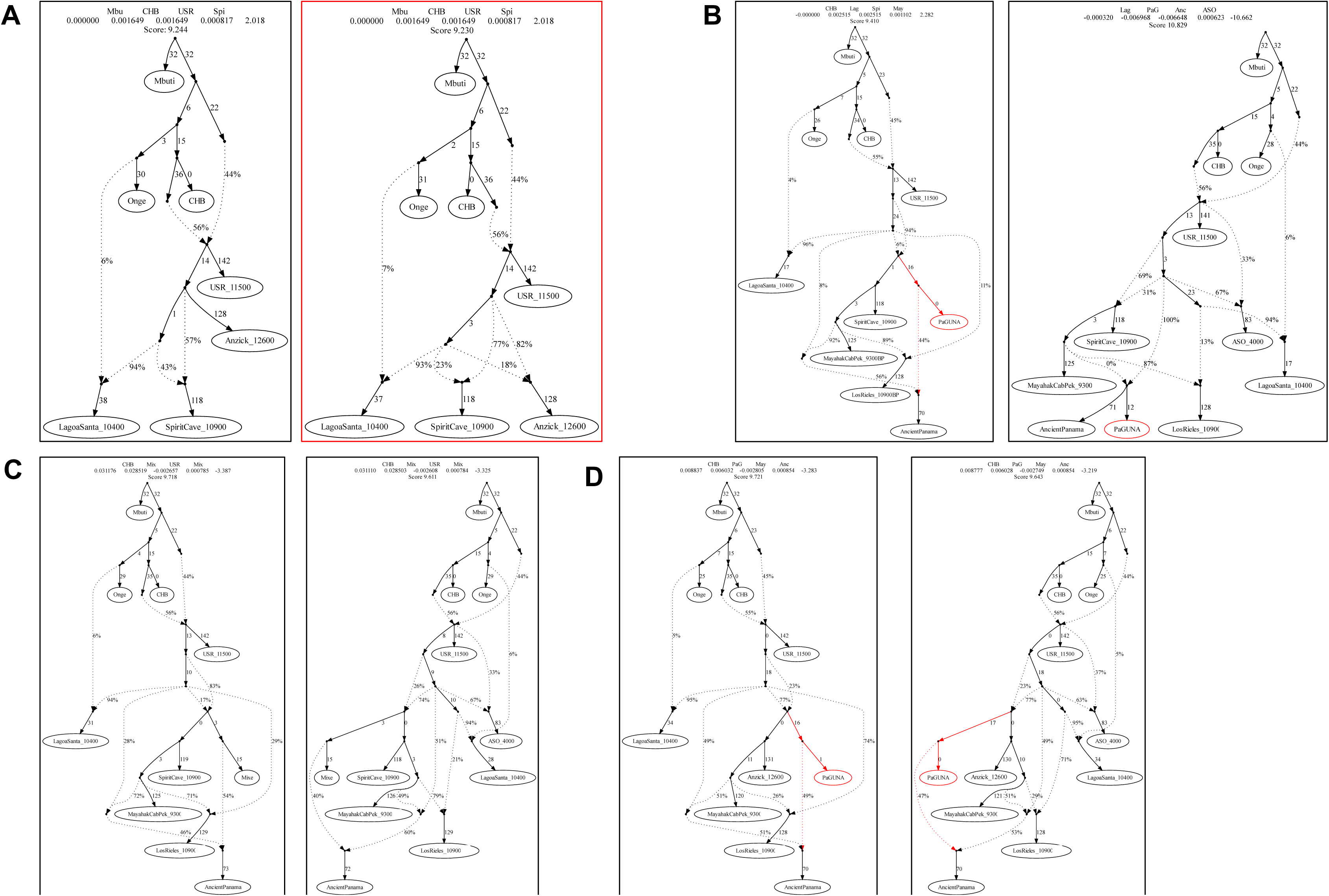
Admixture Graph Modelling the Panama Genetic History Linked to NNA, SNA1 and SNA2. Related to Figure 6B. Possible extensions of the best basal tree (**A**) and the final trees with UPopI (**B**) considering an early admixture between the SNA2 source and SNA1. We also tested Mixe (UPopA) instead of Guna (UPopI) (**C**) and Anzick-1 instead of Spirit Cave (**D**). Best fitting topologies are highlighted in red. See the legend of Figure 6B for further details.

## Notes

### Competing Interest Statement

The authors have declared no competing interest.

